# Screening and identification of key biomarkers associated with endometriosis using bioinformatics and next generation sequencing data analysis

**DOI:** 10.1101/2024.05.06.592657

**Authors:** Basavaraj Vastrad, Chanabasayya Vastrad

**Author notes:** Chanabasayya Vastrad, Chanabasava Nilaya, Bharthinagar, Dharwad 580001, Karanataka, India.

## Abstract

Endometriosis is a common cause of endometrial-type mucosa outside the uterine cavity with symptoms such as painful periods, chronic pelvic pain, pain with intercourse and infertility. However, the early diagnosis of endometriosis is still restricted. The purpose of this investigation is to identify and validate the key biomarkers of endometriosis. Next generation sequencing (NGS) dataset GSE243039 was obtained from the Gene Expression Omnibus (GEO) database, and differentially expressed genes (DEGs) between endometriosis and normal control samples were identified. After screening of DEGs, gene ontology (GO) and REACTOME pathway enrichment analyses were performed. Furthermore, a protein-protein interaction (PPI) network was constructed and modules were analysed using the Human Integrated Protein-Protein Interaction rEference (HIPIE) database and Cytoscape software, and hub genes were identified. Subsequantely, a network between miRNAs and hub genes, and network between TFss and hub genes were constructed using the miRNet and NetworkAnalyst tool, and possible key miRNAs and TFs were predicted. Finally, receiver operating characteristic curve (ROC) analysis was used to validate the hub genes. A total of 958 DEGs, including 479 up regulated genes and 479 down regulated genes, were screened between endometriosis and normal control samples. GO and REACTOME pathway enrichment analyses of the 958 DEGs showed that they were mainly involved in multicellular organismal process, developmental process, signaling by GPCR and muscle contraction. Further analysis of the PPI network and modules identified 10 hub genes, including VCAM1, SNCA, PRKCB, ADRB2, FOXQ1, MDFI, ACTBL2, PRKD1, DAPK1 and ACTC1. Possible target miRNAs, including hsa-mir-3143 and hsa-mir-2110, and target TFs, including TCF3 and CLOCK, were predicted by constructing a miRNA-hub gene regulatory network and TF-hub gene regulatory network. This investigation used bioinformatics techniques to explore the potential and novel biomarkers. These biomarkers might provide new ideas and methods for the early diagnosis, treatment, and monitoring of endometriosis.

## Introduction

Endometriosis is one of the most important chronic inflammatory disease and has become the chief cause of serious reproductive and general health condition [1]. Endometriosis is characterized by presence of endometrial-type mucosa outside the uterine cavity [2]. The clinical incidence of endometriosis is high, and its main features include dysmenorrhea, dyspareunia, chronic pelvic pain, irregular uterine bleeding and infertility [3], which places a great burden on the economy of health and reduces quality of life in worldwide, 10% of women of reproductive age are diagnosed with endometriosis each year [4]. These patients have higher risks of gynecological cancer (ovarian, endometrial and cervical cancers) [5], polycystic ovary syndrome [6], cardiovascular diseases [7], obesity [8], gestational diabetes mellitus [9], diabetes mellitus [10] and hypertension [11]. Studies have revealed that the progression of endometriosis is related to genetic risk factors [12] as well as environmental factors [13]. Because of this disorder complex pathogenesis, it is mainly treated by gynecological surgery [14], oral contraceptives [15], progestins [16], nonsteroidal anti-inflammatory drugs [17], and gonadotropin-releasing hormone agonists [18]. In current years, molecular biomarkers were demonstrated highly useful as clinical tools for endometriosis diagnosis [19]. Thus, it is essential to find new methods for early detection and treatment for better outcomes.

The underlying complex molecular mechanisms in endometriosis pose a special challenge to daily clinical practice. Genes including CYR61 [20], ESR2 and CYP19A1 [21], HOXA10 [22], FOXD3 [23], and LOXL1 and HTRA1 [24] as well as signaling pathways include AKT and ERK signaling pathways [25], Wnt/β-catenin signaling pathway [26], PI3K-Akt-mTOR and MAPK signaling pathways [27], notch signaling pathway [28] and MAPK/ERK signal pathway [29], and are involved in the progression of endometriosis. Taken together, current evidence suggests that the genes and signal pathways are closely related to the progression of endometriosis.

The mechanisms of endometriosis at the molecular level are essentially important for treating the disease. With the wide application of next generation sequencing (NGS) technology, endometriosis related genes have been widely identified, which is a key step in exploring the complex pathology of endometriosis and finding drugs that combat the illness. Numerous NGS data of gene expression have been published in public databases such as NCBI Gene Expression Omnibus (GEO) [https://www.ncbi.nlm.nih.gov/geo/] [30] during the past few years, and they are being increasingly used in bioinformatics and NGS data analysis to explore target genes or proteins associated in various diseases [31–32].

Bioinformatics and network analysis of NGS data is an effective way to explore gene expression profiles in the pathogenesis of disease. Therefore, this investigation aimed to use bioinformatics analysis to identify hub genes and molecular pathways involved in endometriosis, to identify key diagnostic or therapeutic biomarkers. We obtained DEGs between endometriosis and normal control samples from GSE243039, a gene expression profile downloaded from the GEO database. Immediately after, we performed gene ontology (GO) and REACTOME pathway enrichment analysis on these DEGs. By constructing PPI networks, we screened for the significant modules and hub genes. We constructed miRNA-hub gene regulatory network and TF-hub gene regulatory network, we screened for the miRNAs, TFs and hub genes. To validate that these hub genes can serve as molecular markers of endometriosis, we determined hub genes by using receiver operating characteristic curve (ROC) analysis. This investigation will improve our understanding of the molecular pathogenesis of endometriosis and provide genomic-targeted therapy options for endometriosis.

## Materials and Methods

### Next generation sequencing (NGS) data source

The NGS dataset GSE243039 was obtained from the GEO database. The GSE243039 dataset included 20 endometriosis samples and 20 normal control samples. The platform used was the GPL24676 Illumina NovaSeq 6000 (Homo sapiens).

### Identification of DEGs

The limma R/Bioconductor software package [33] was used to perform the identification of DEGs between endometriosis samples and normal control samples. We adjusted p-value to correct the false positive error caused by the multiple tests and determined it by the Benjamini & Hochberg method [34], which is the common tools to minimize the false discovery rate. The cutoff criteria were |logFC| > 1.304 (log2 fold change) for up regulated genes, |logFC| > 1.304 (log2 fold change) < -1.2644 for down regulated genes and a adj.P.Val<0.05. Thereafter, we used R packages “ggplot2” and “gplot” to show the DEGs with up regulated and down regulated expression in volcano plot and heatmap, respectively.

### GO and pathway enrichment analyses of DEGs

GO enrichment analysis (http://www.geneontology.org) [35] was frequently used to annotate the degree of gene function terms in DEGs, which included biological process (BP), cellular component (CC), and molecular function (MF). REACTOME (https://reactome.org/) [36] pathway enrichment analysis was used to demonstrate enriched signaling pathways in DEGs. The g:Profiler (http://biit.cs.ut.ee/gprofiler/) [37] was used to perform GO and REACTOME pathway enrichment analysis of DEGs. P<0.05 was considered to represent statistical significance.

### Construction of the PPI network and module analysis

To ensure the optimal graphical display of protein interactions of DEGs, Human Integrated Protein-Protein Interaction rEference (HIPIE) (https://cn.string-db.org/) [38] was used to generate the PPI network. The software Cytoscape (version 3.10.1) (http://www.cytoscape.org/) [39] was used to visualize the PPI network. The Network Analyzer in Cytoscape was utilized to calculate node degree [40], betweenness [41], stress [42] and closeness [43]. The PEWCC Cytoscape software plugin [44] was used to create modules in the PPI network of endometriosis.

### Construction of the miRNA-hub gene regulatory network

The significant miRNAs were identified from miRNA-hub gene regulatory network analysis through the TarBase, miRTarBase, miRecords, miRanda (S mansoni only), miR2Disease, HMDD, PhenomiR, SM2miR, PharmacomiR, EpimiR, starBase, TransmiR, ADmiRE and TAM 2 via miRNet database (https://www.mirnet.ca/) [45]. This networks was visualized with Cytoscape [39] and the significant hub genes and miRNAs were selected via the Network Analyzer plugin in Cytoscape based on the degree connectivity.

### Construction of the TF-hub gene regulatory network

The significant transcription factors (TFs) were identified from TF-hub gene regulatory network analysis through the CHEA via NetworkAnalyst database (https://www.networkanalyst.ca/) [46]. This networks was visualized with Cytoscape [39] and the significant hub genes and TFs were selected via the Network Analyzer plugin in Cytoscape based on the degree connectivity.

### Receiver operating characteristic curve (ROC) analysis

ROC analysis was performed to predict the diagnostic effectiveness of biomarkers by pROC package of R software [47]. The area under the curve (AUC) value was utilized to determine the diagnostic effectiveness in discriminating endometriosis from normal control samples.

## Results

### Identification of DEGs

The DEGs were screened by “limma” package (|logFC| > 1.304 (log2 fold change) for up regulated genes, |logFC| > 1.304 (log2 fold change) < -1.2644 for down regulated genes and adj.P.Val<0.05). The GSE243039 dataset contained 958 DEGs, including 479 up regulated genes and 479 down regulated genes and are listed in Table 1. The volcano plot (Fig.1) was used to show the expression pattern of DEGs in endometriosis. The heatmap of the DEGs is shown in Fig. 2.

**Fig. 1.**
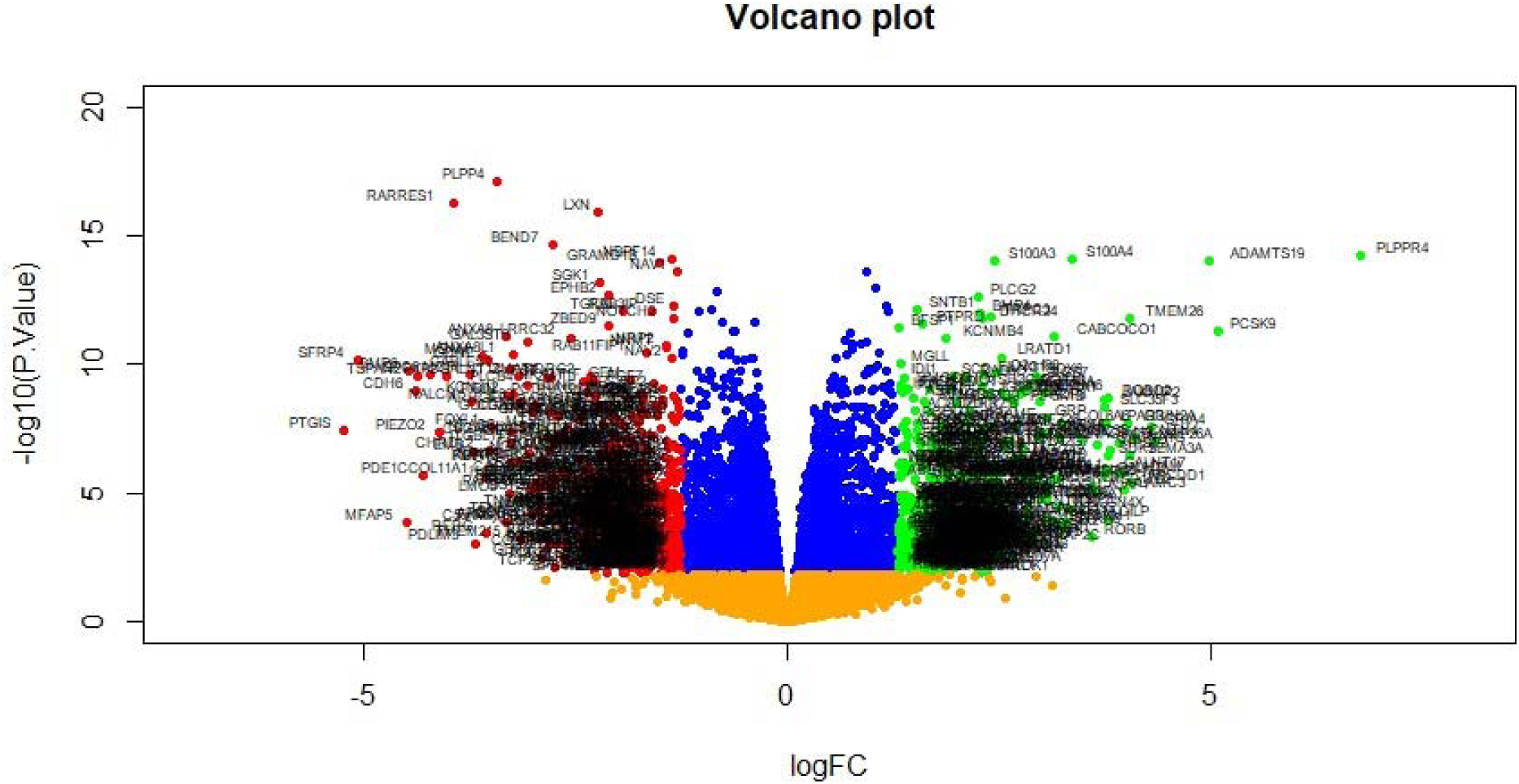
Volcano plot of differentially expressed genes. Genes with a significant change of more than two-fold were selected. Green dot represented up regulated significant genes and red dot represented down regulated significant genes.

**Fig. 2.**
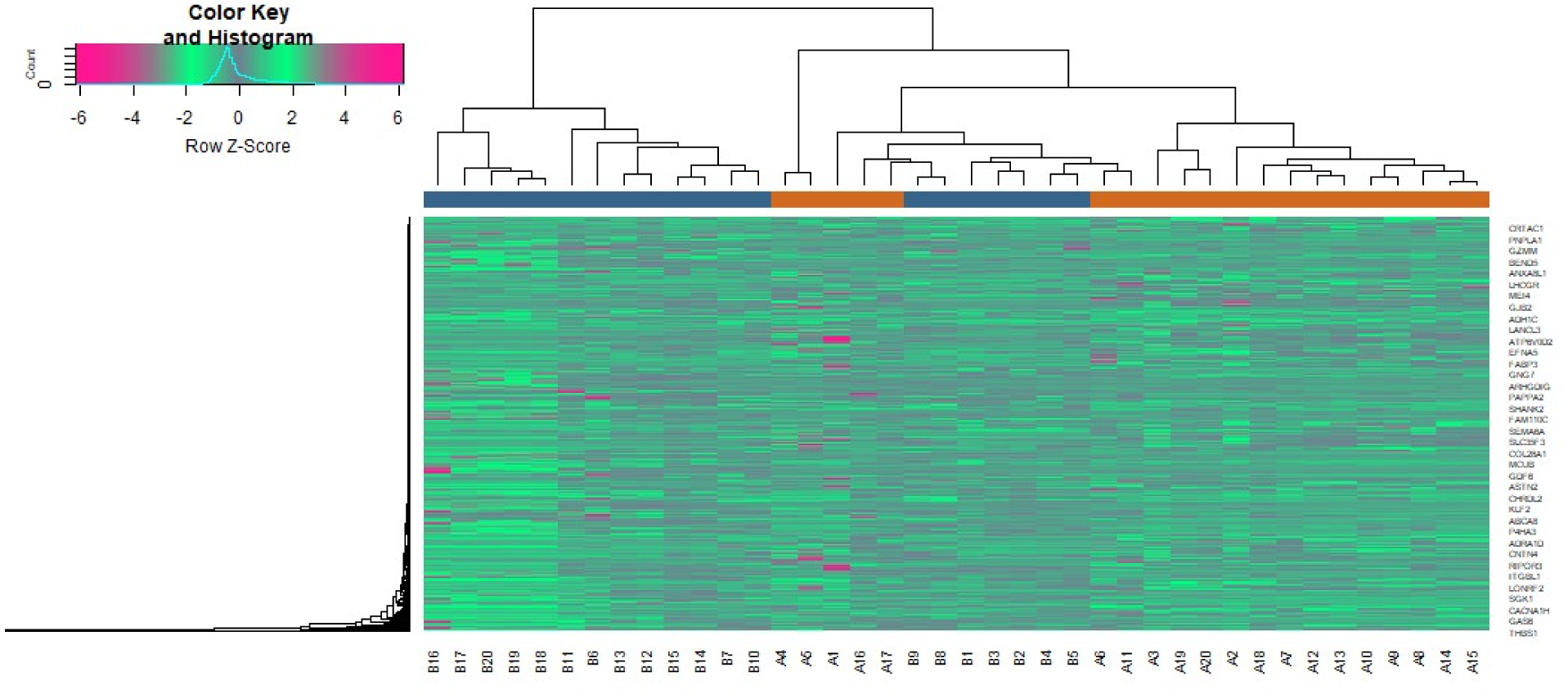
Heat map of differentially expressed genes. Legend on the top left indicate log fold change of genes. (A1 – A20 = Endometriosis samples; B1 – B20 = Normal control samples)

**Table 1.**
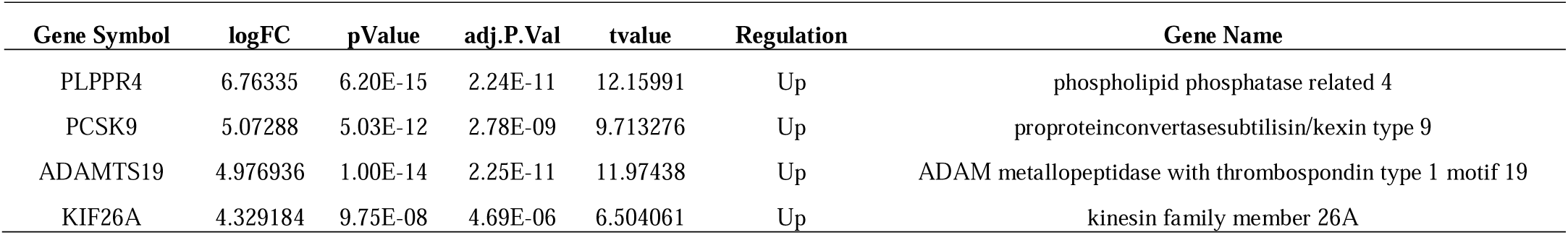
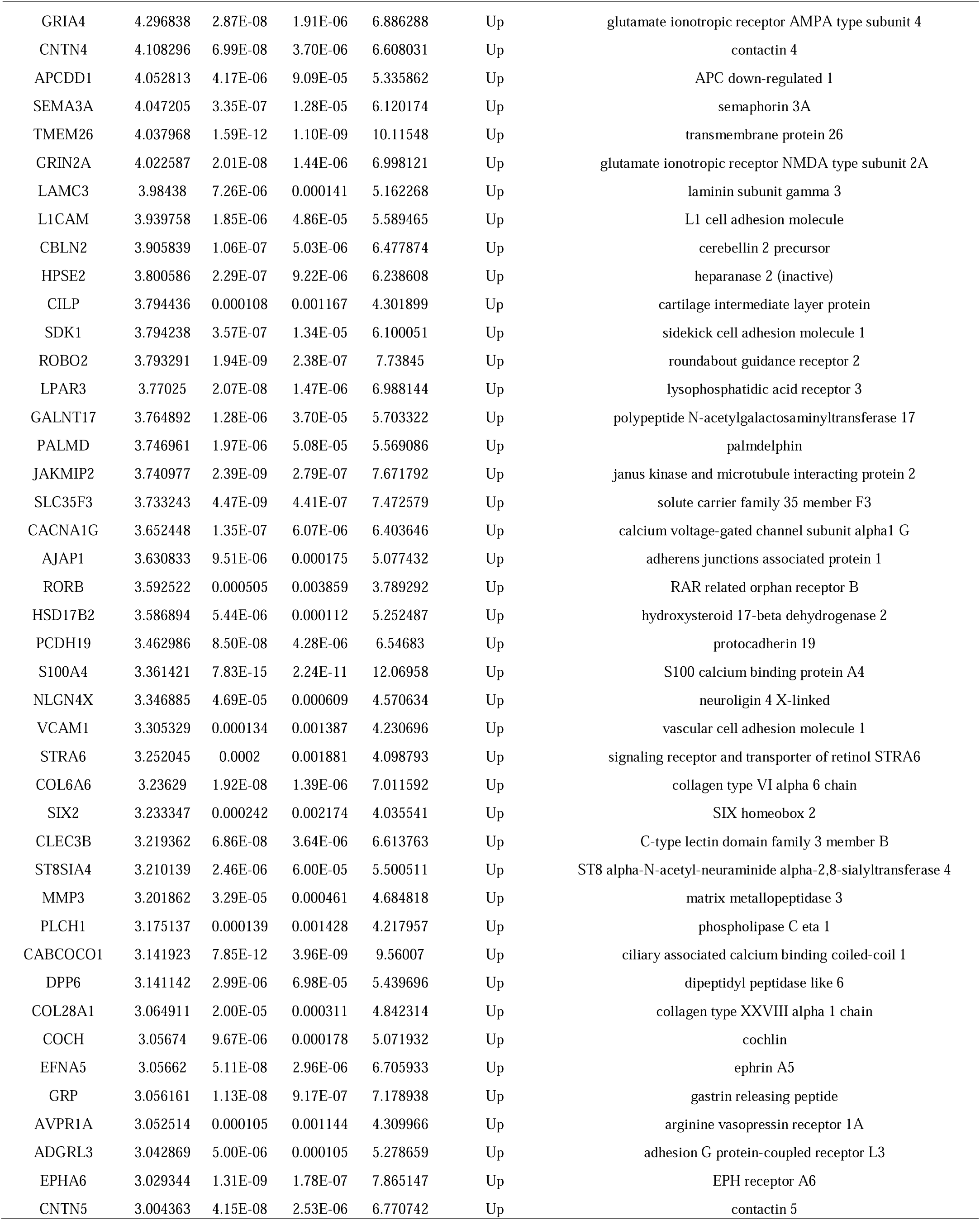
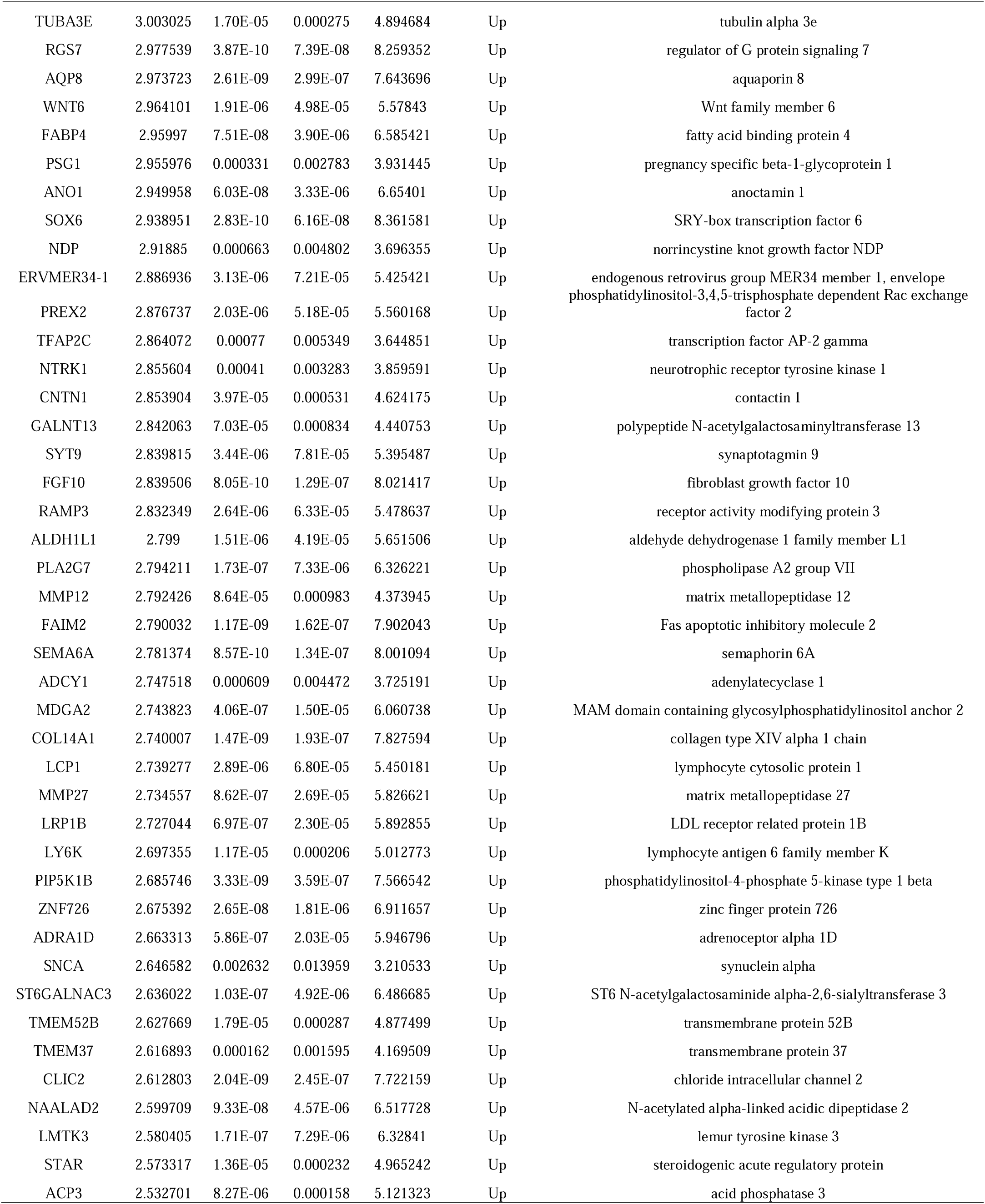
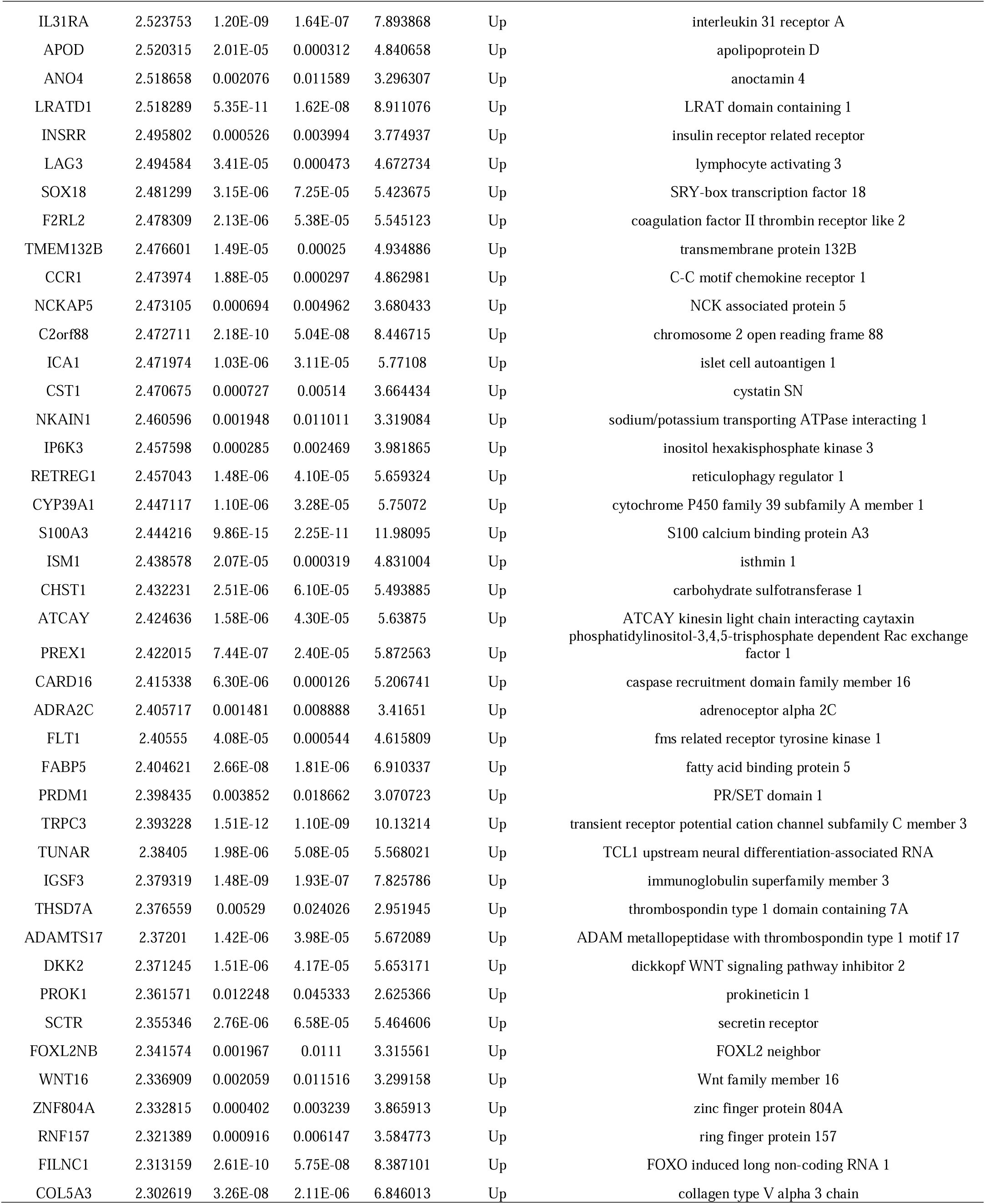
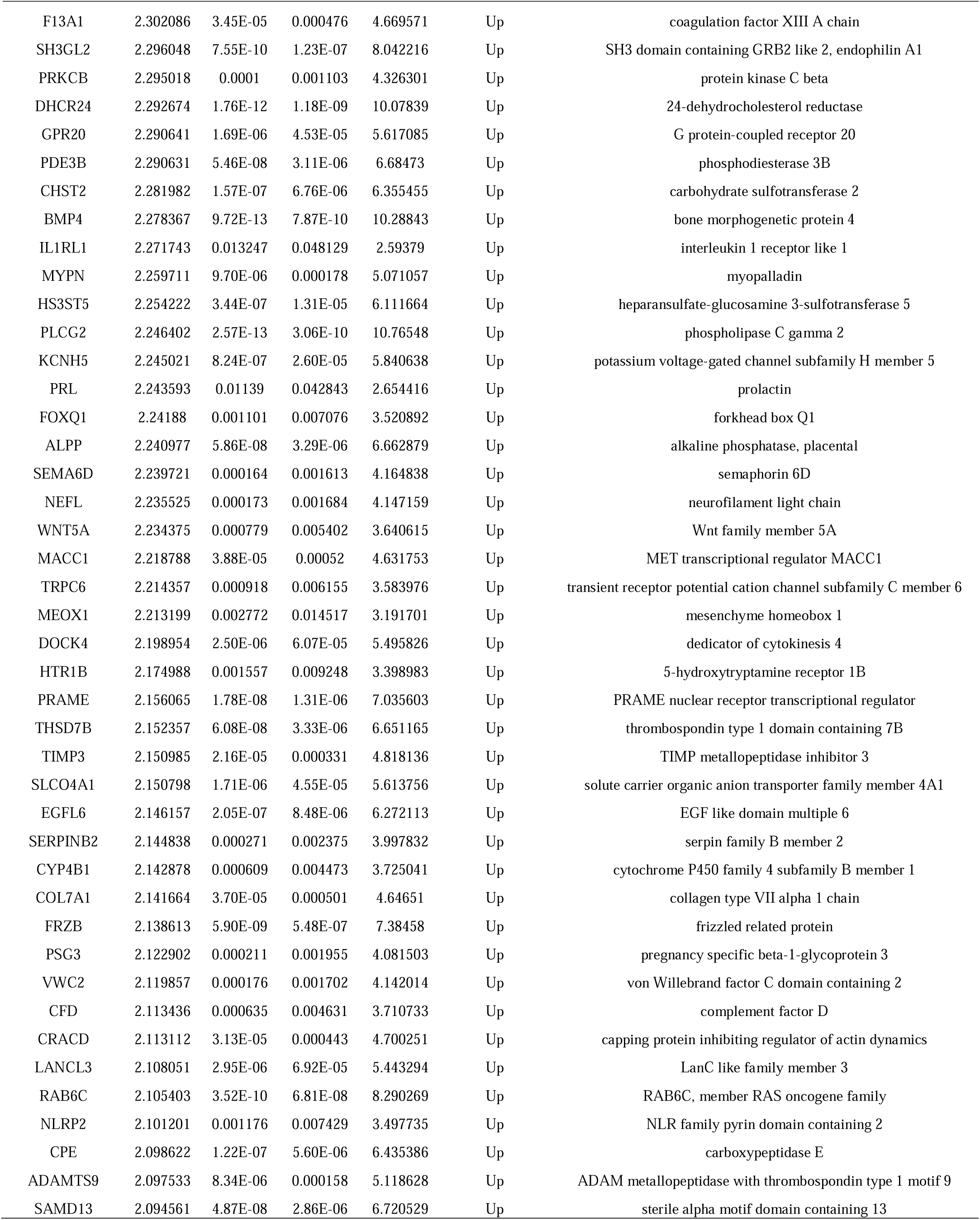
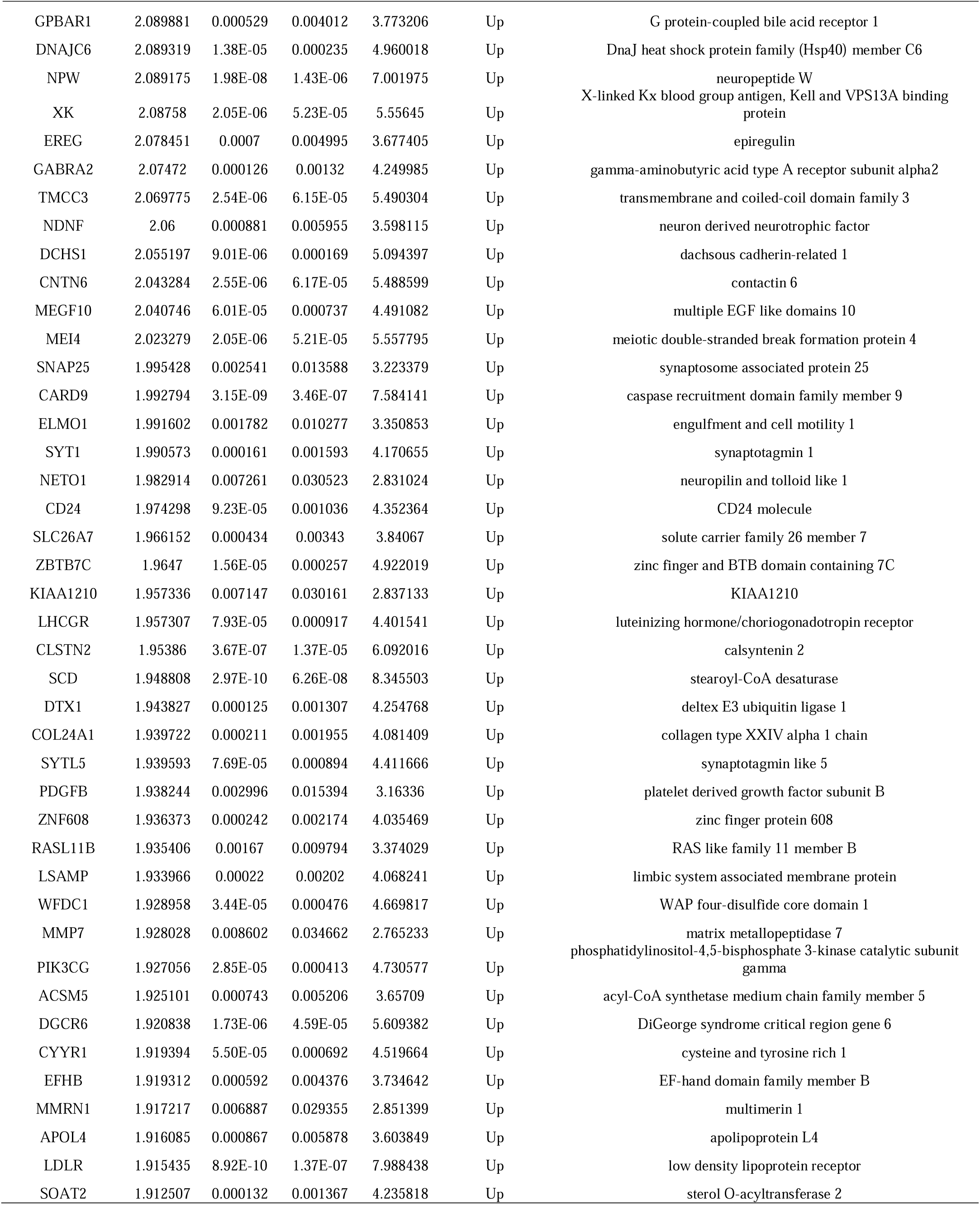
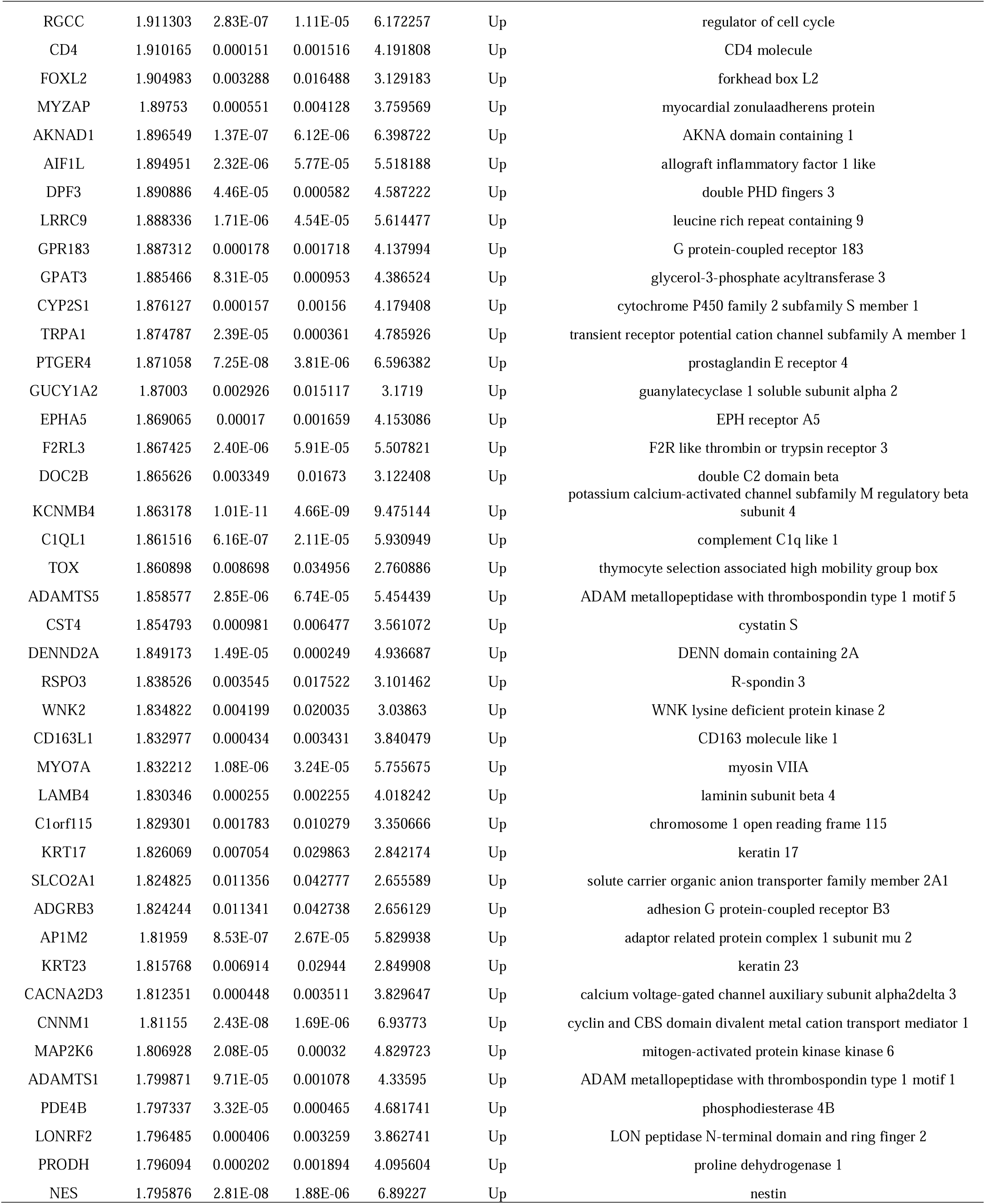
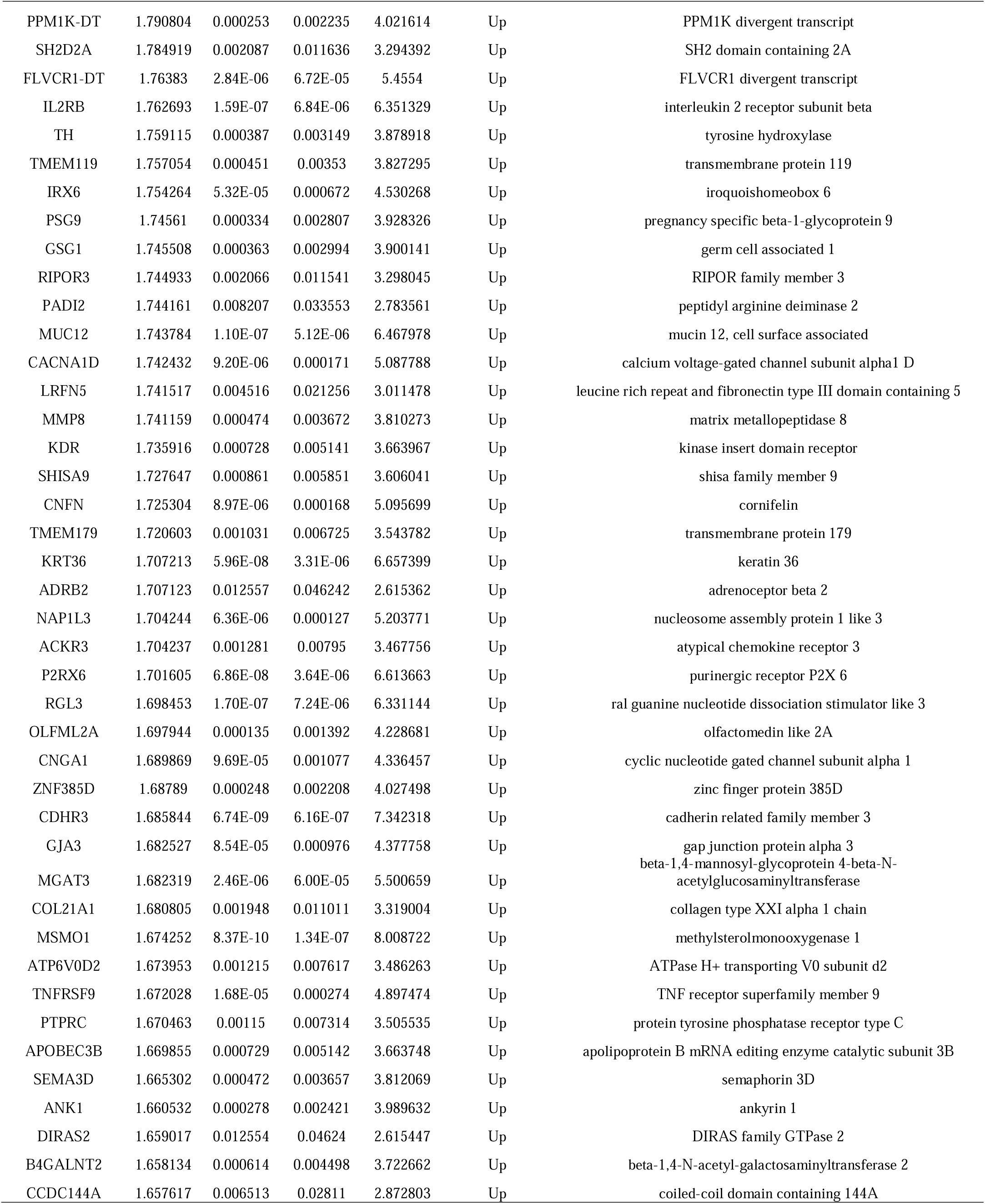
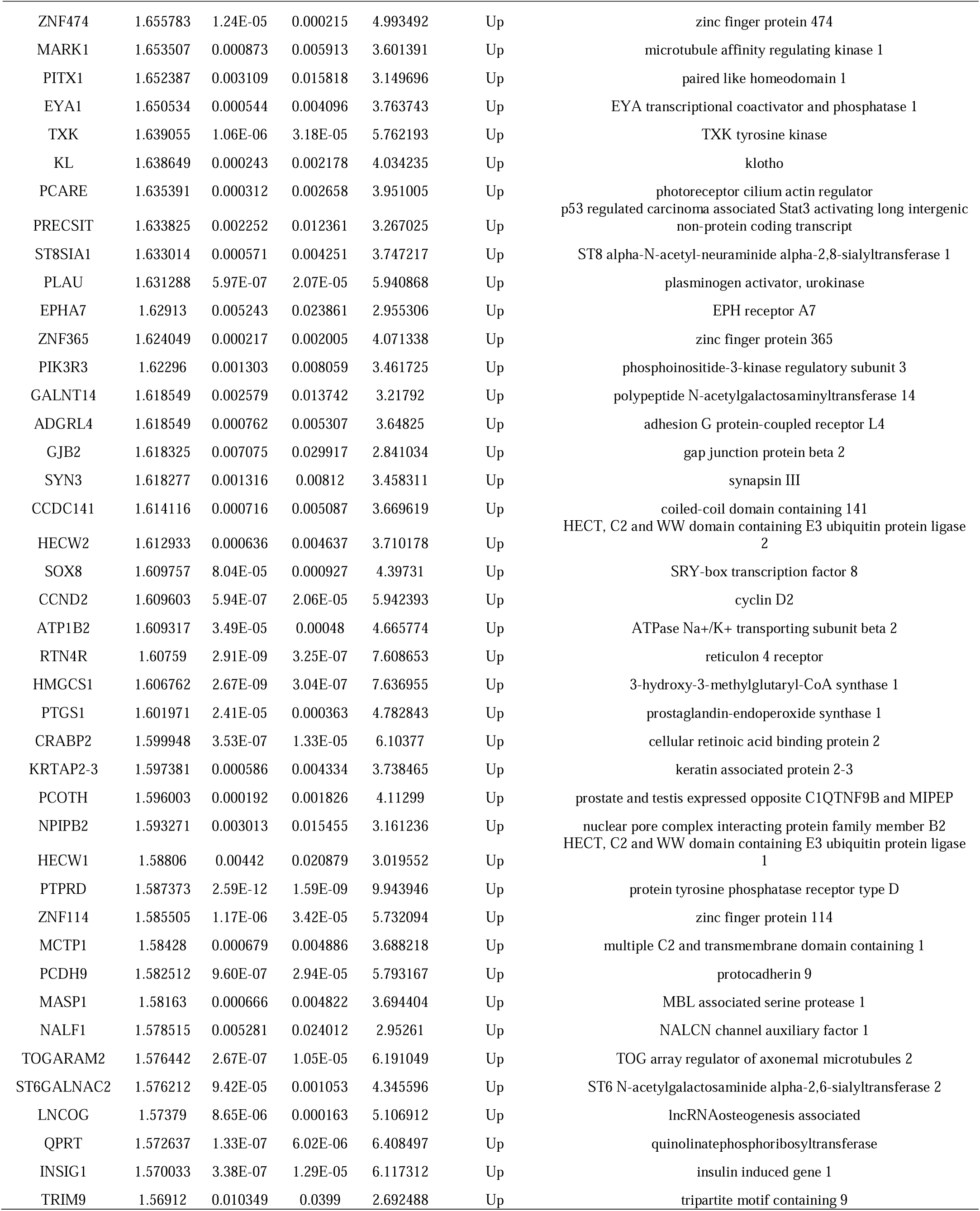
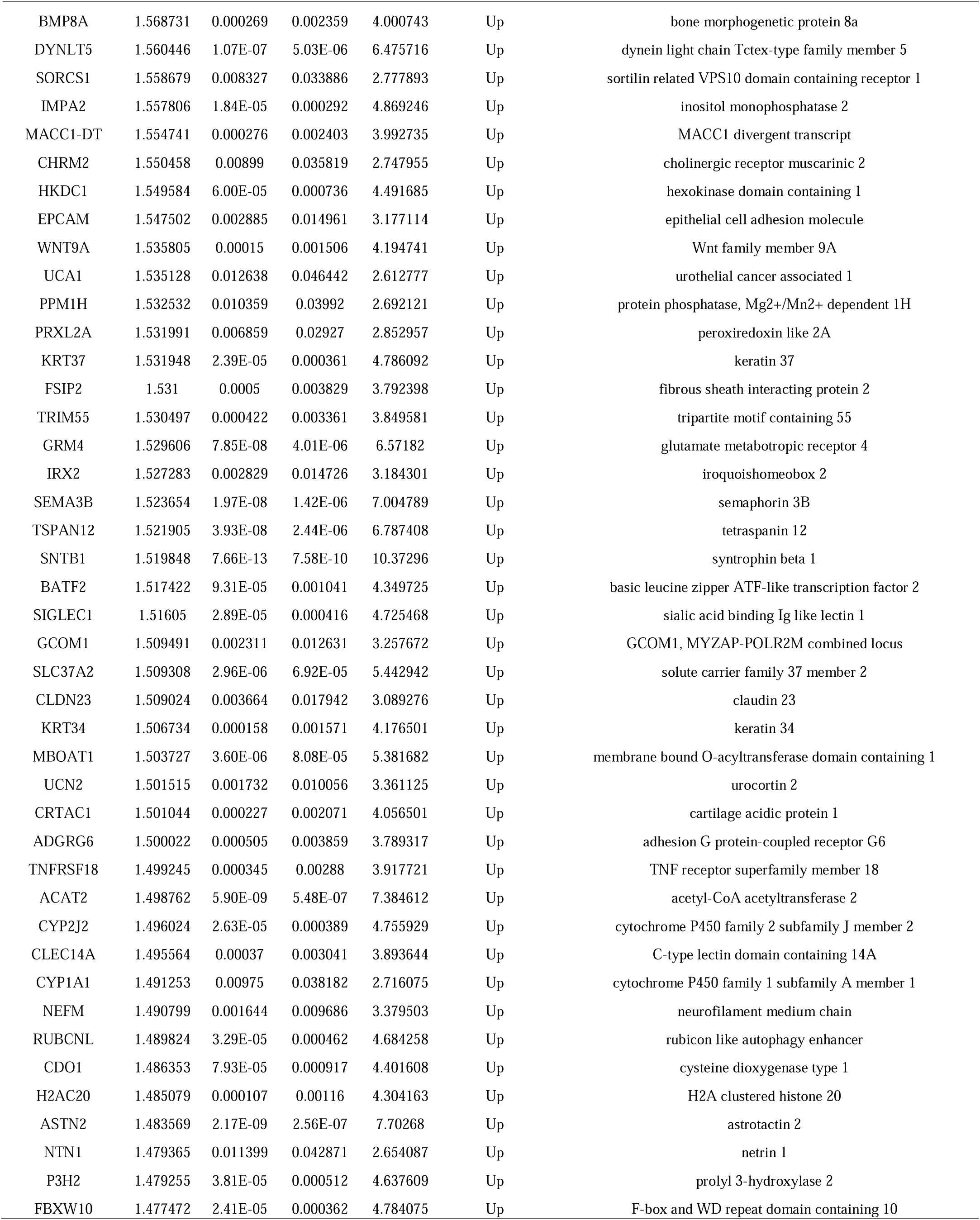
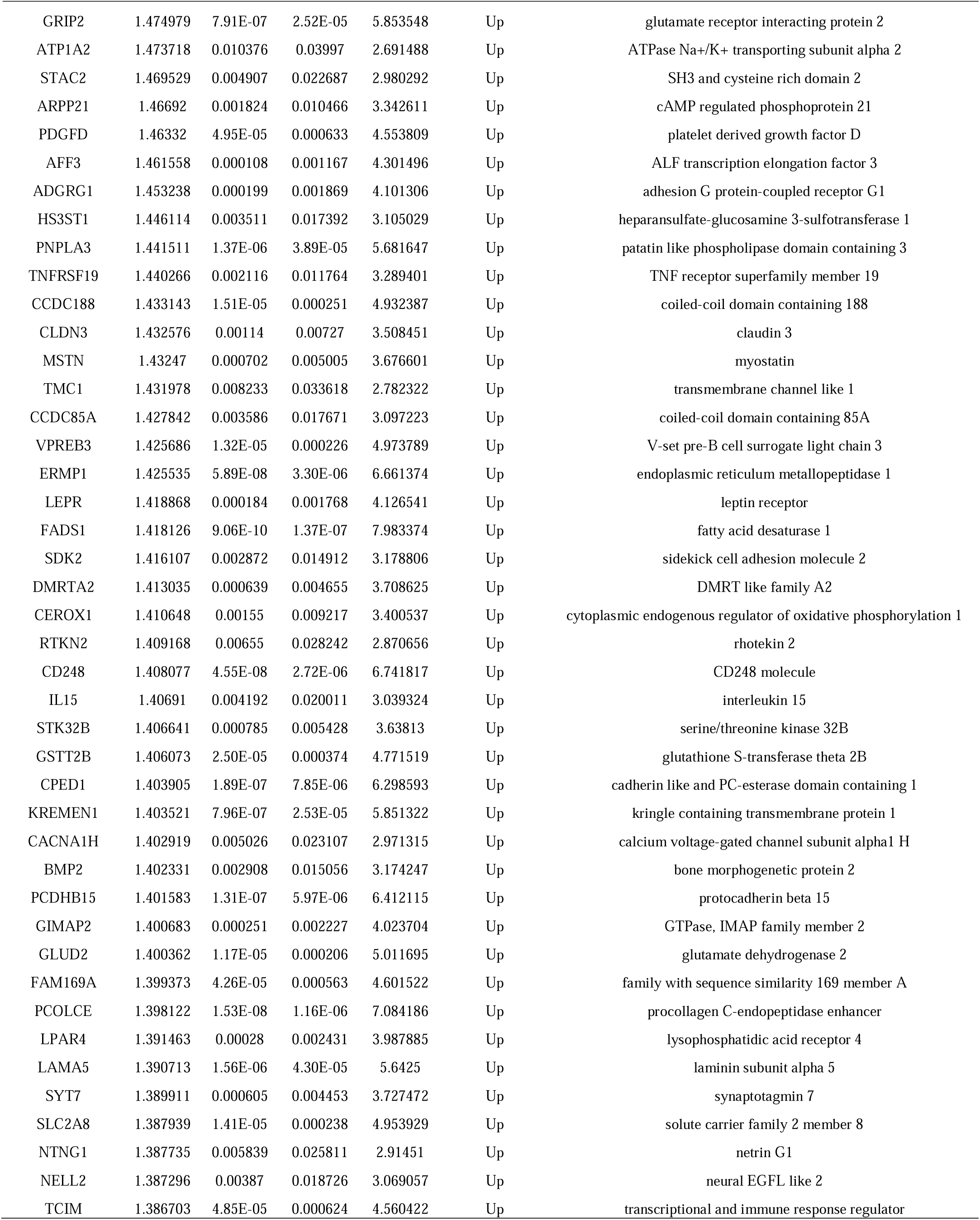
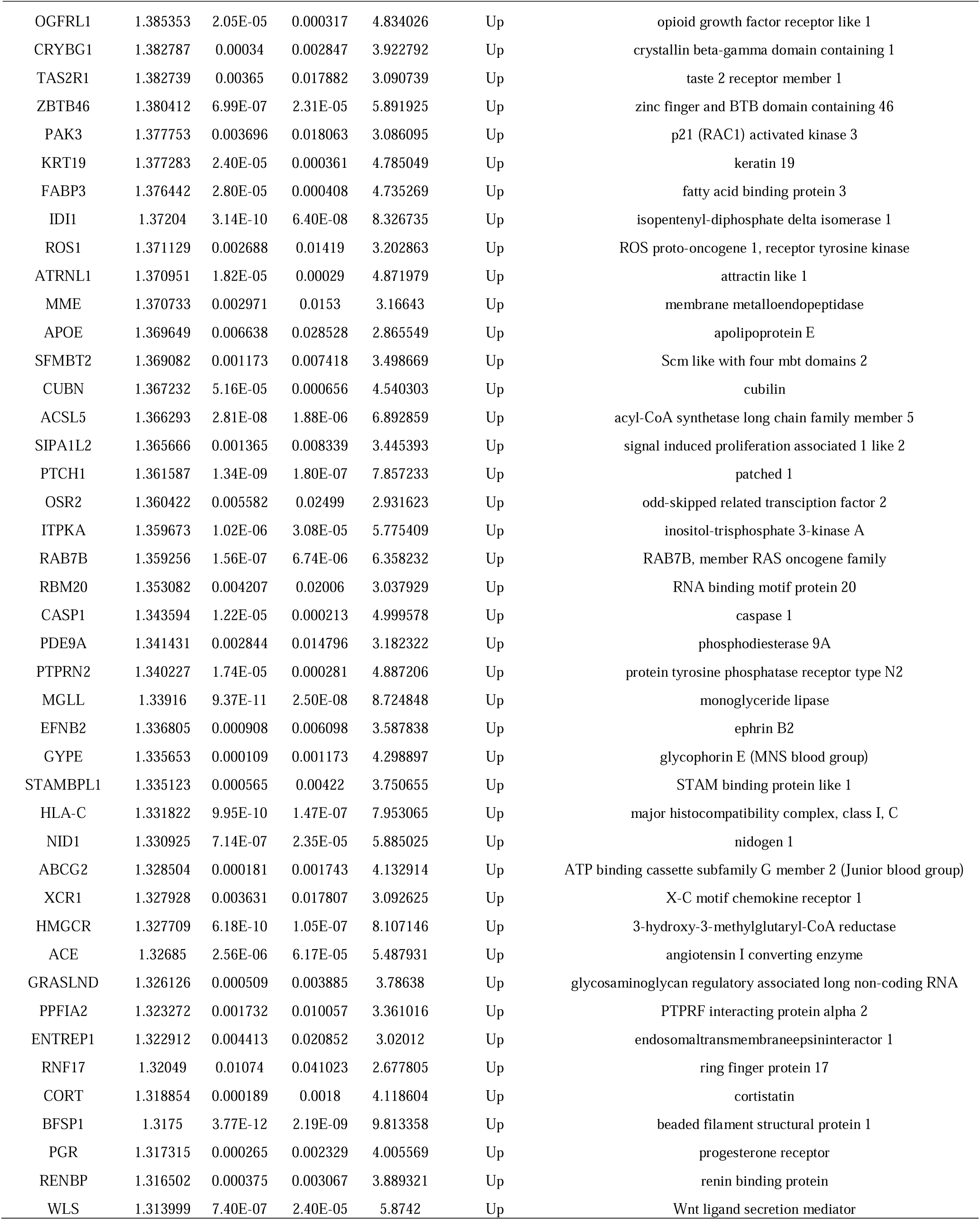
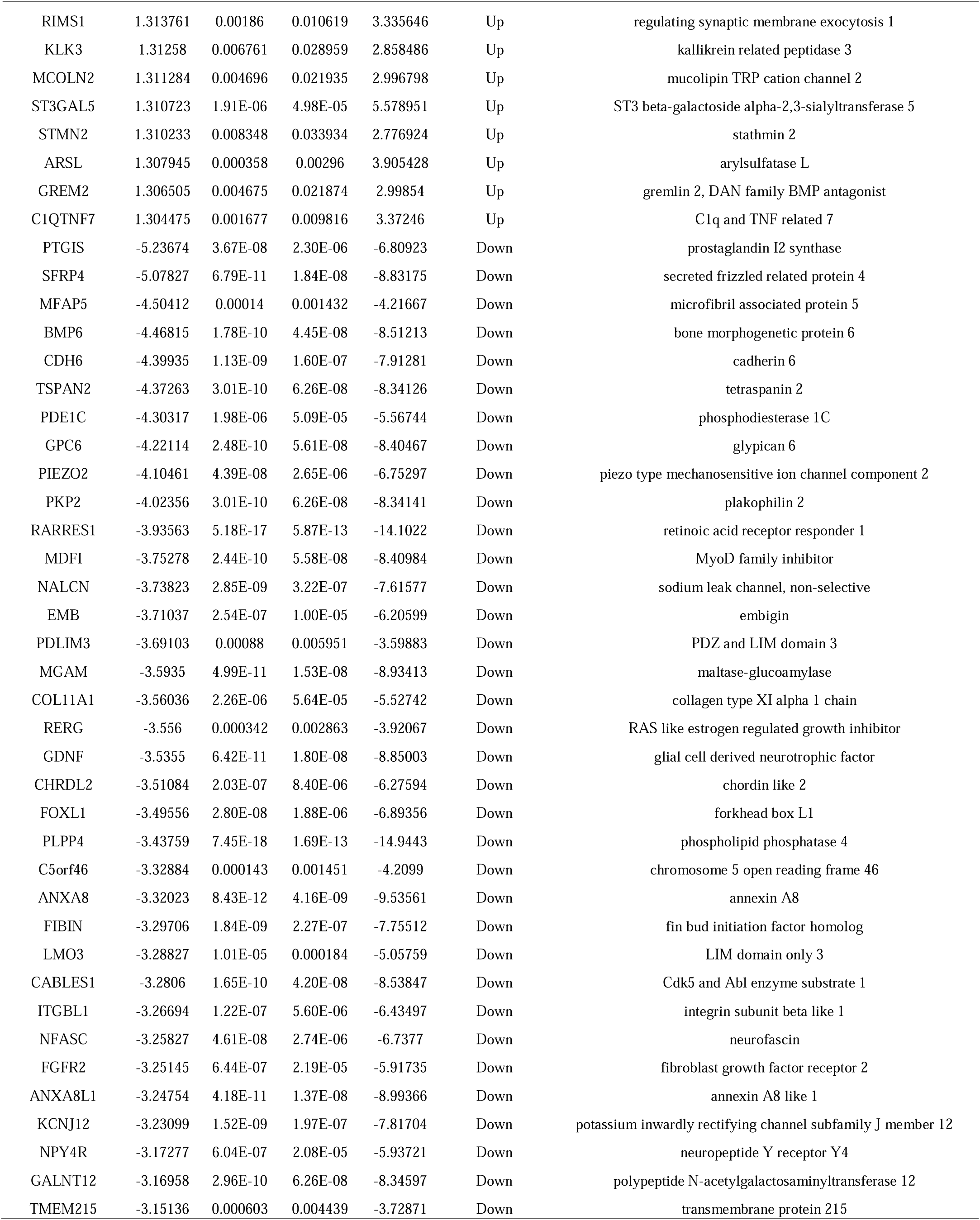
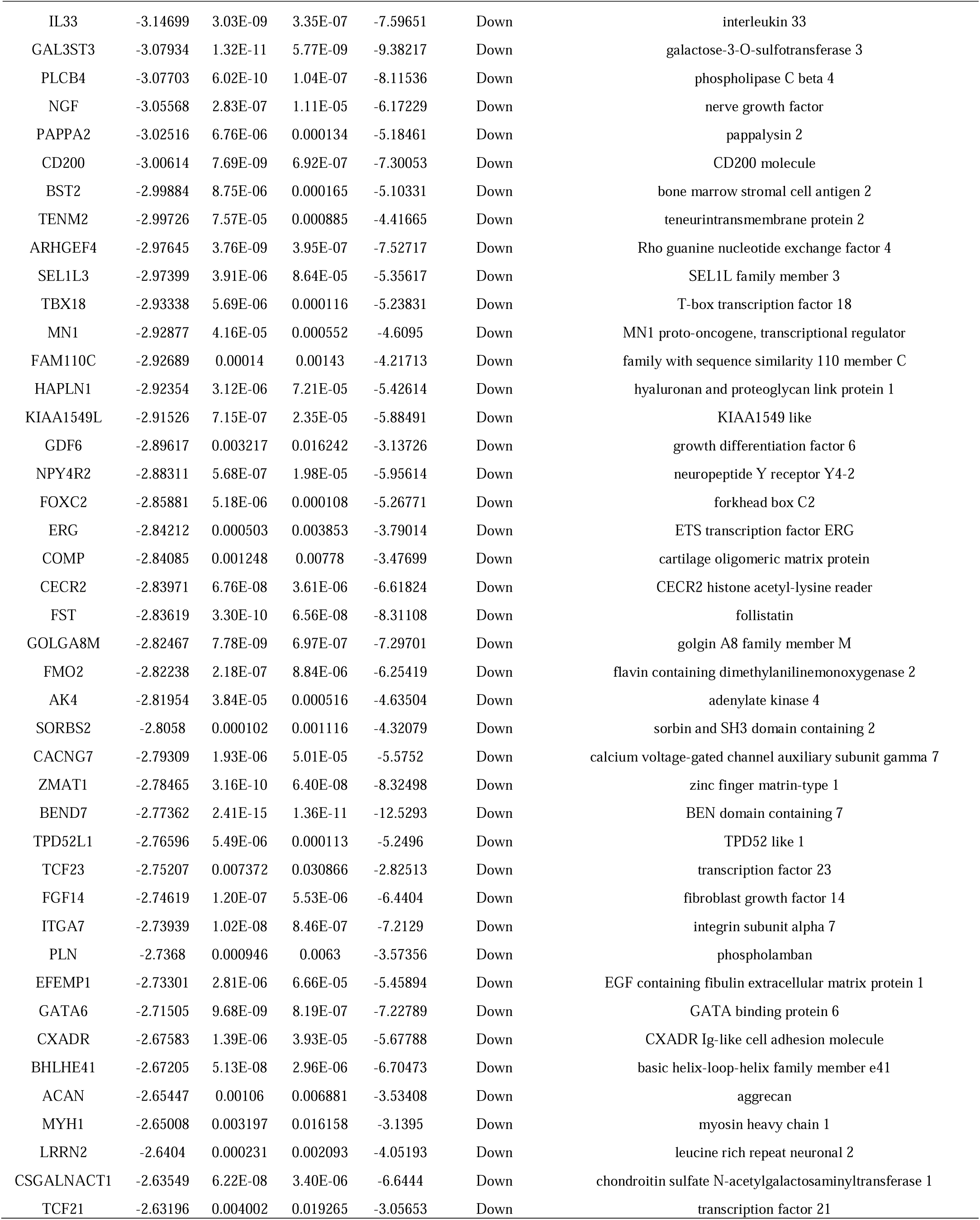
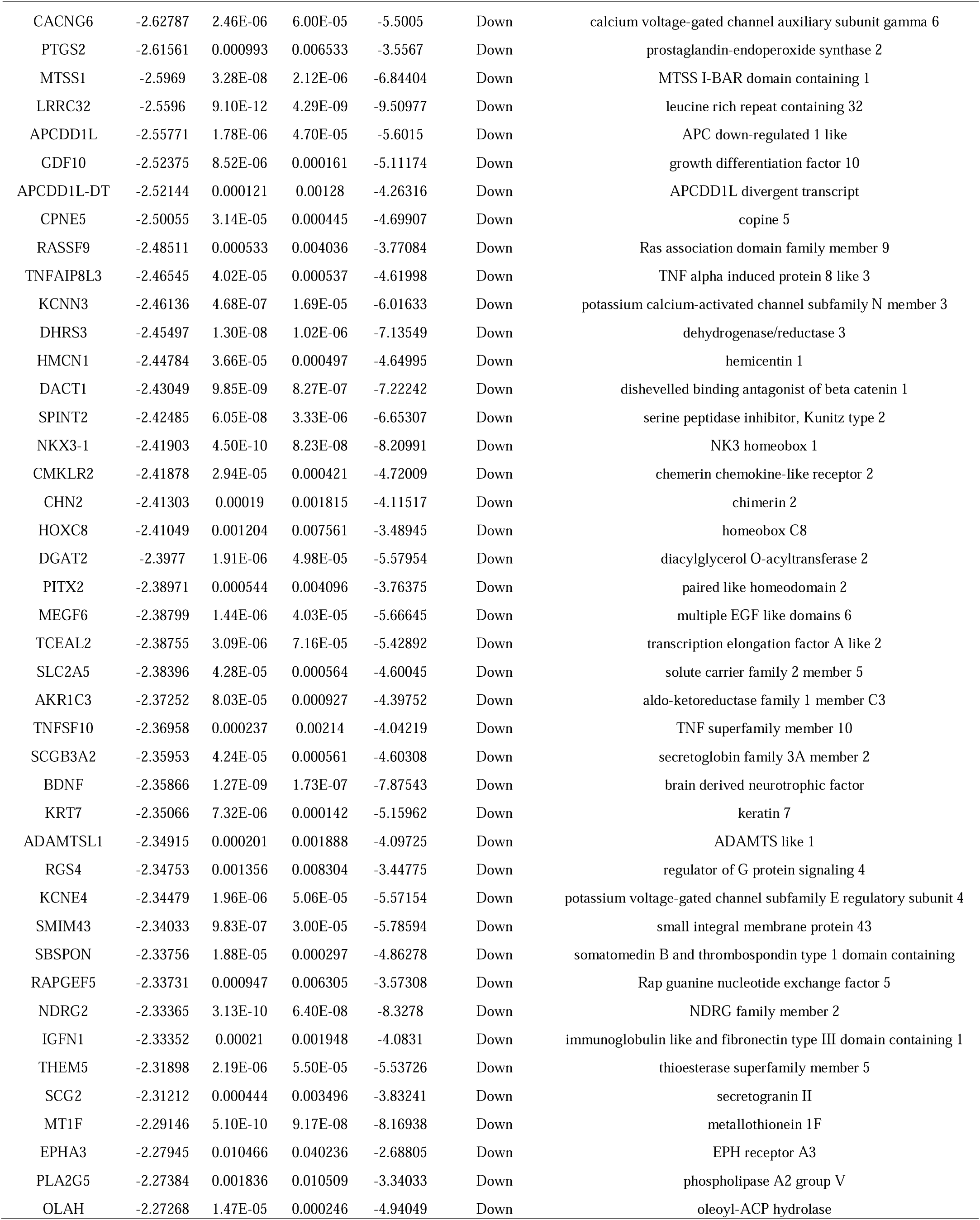
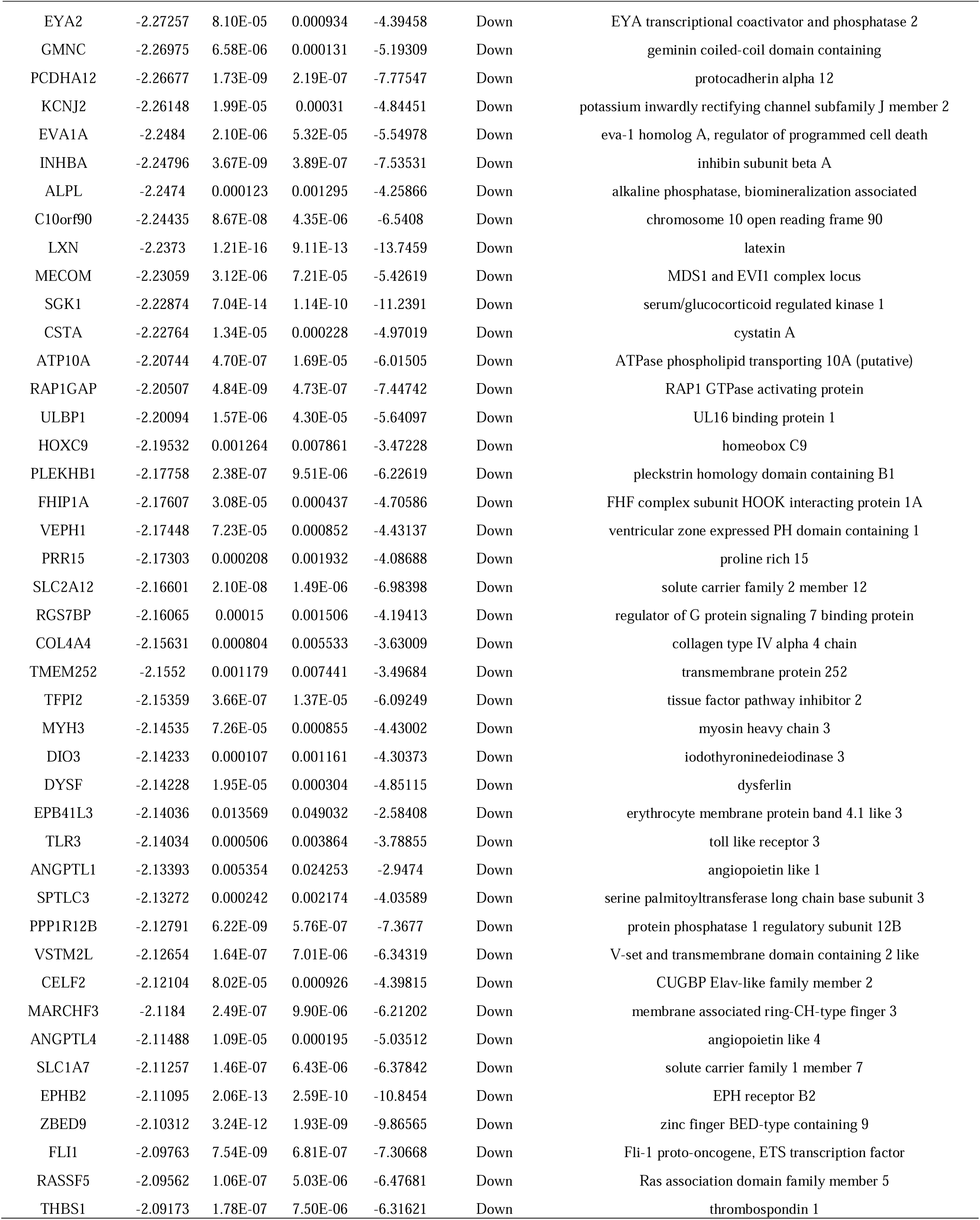
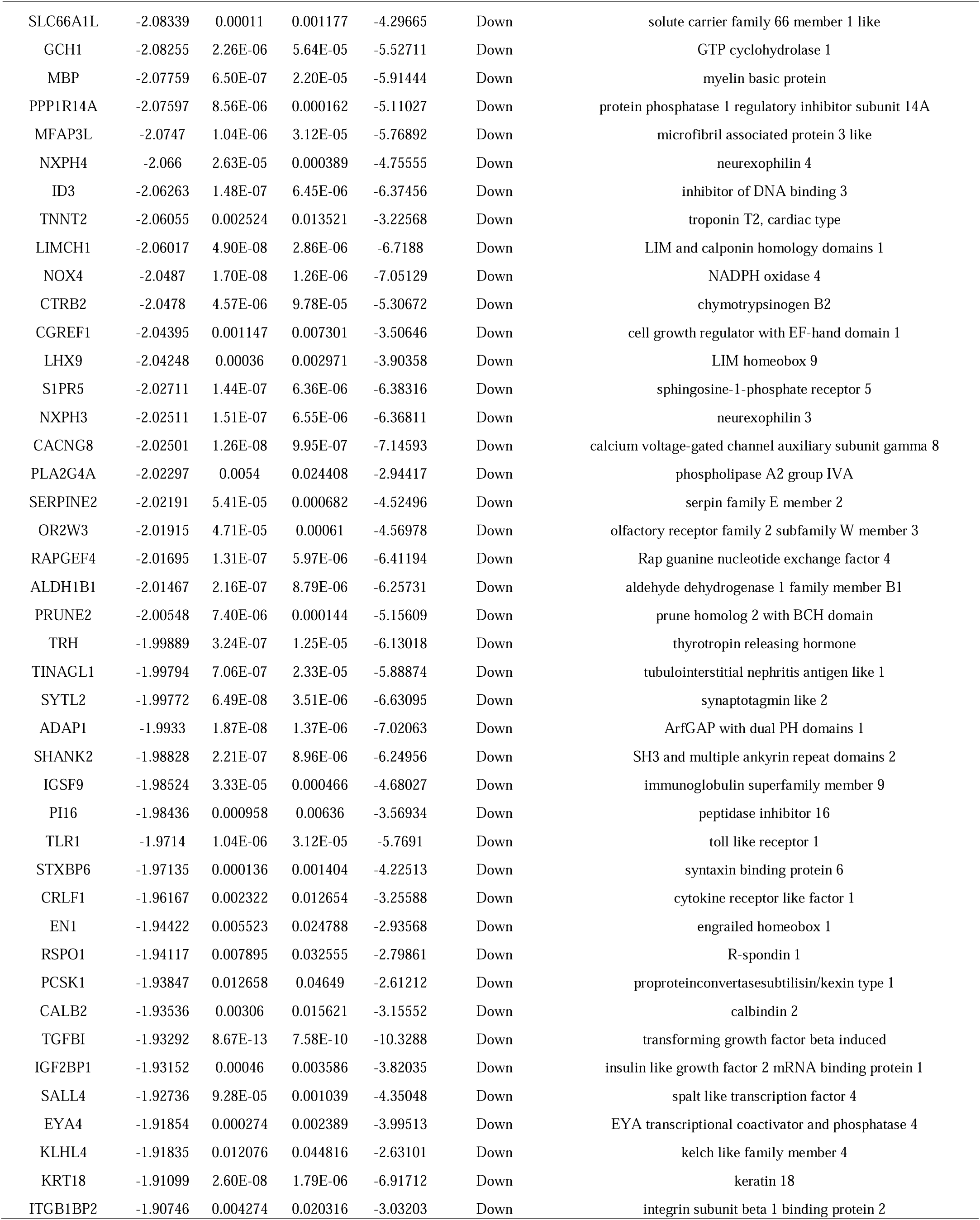
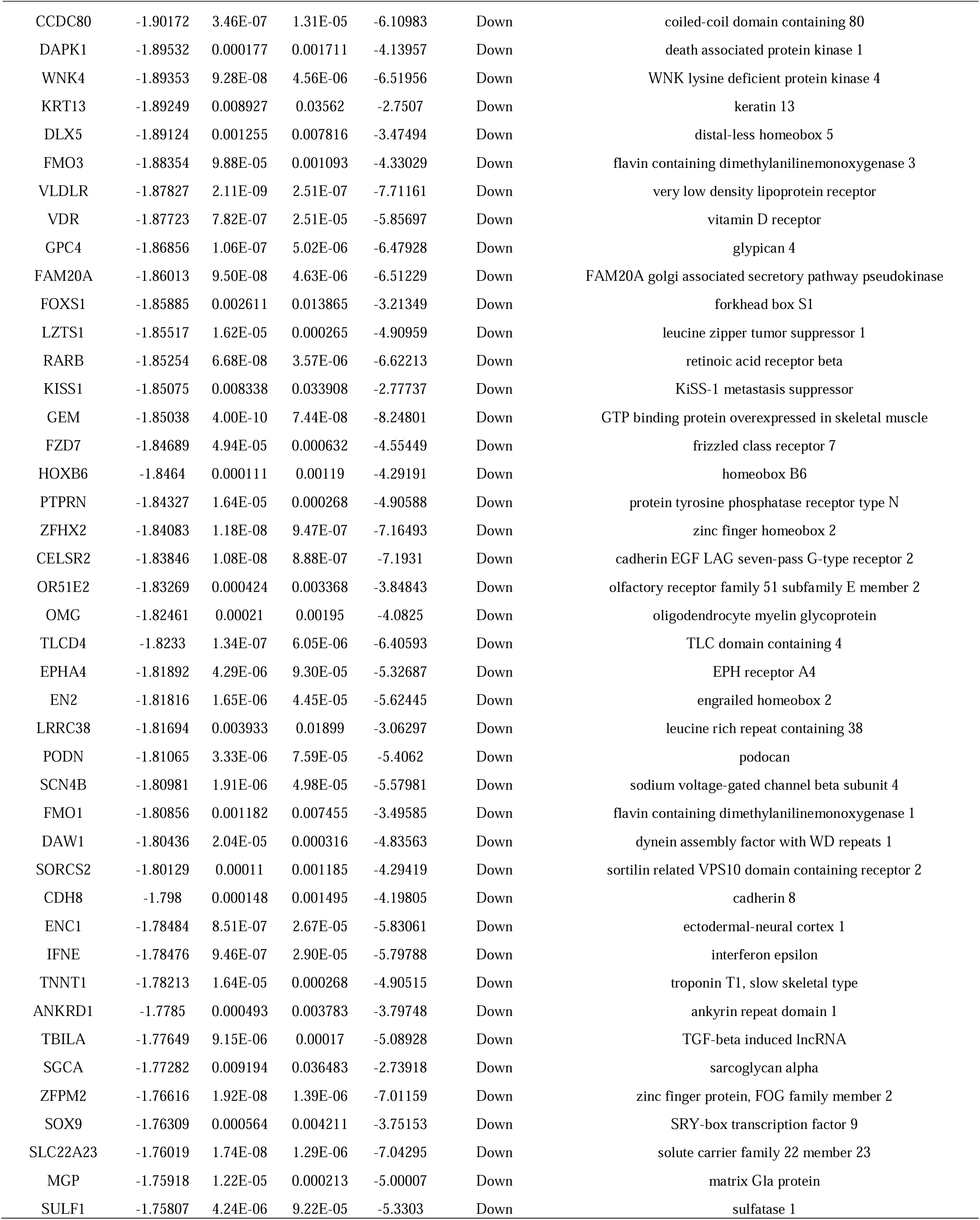
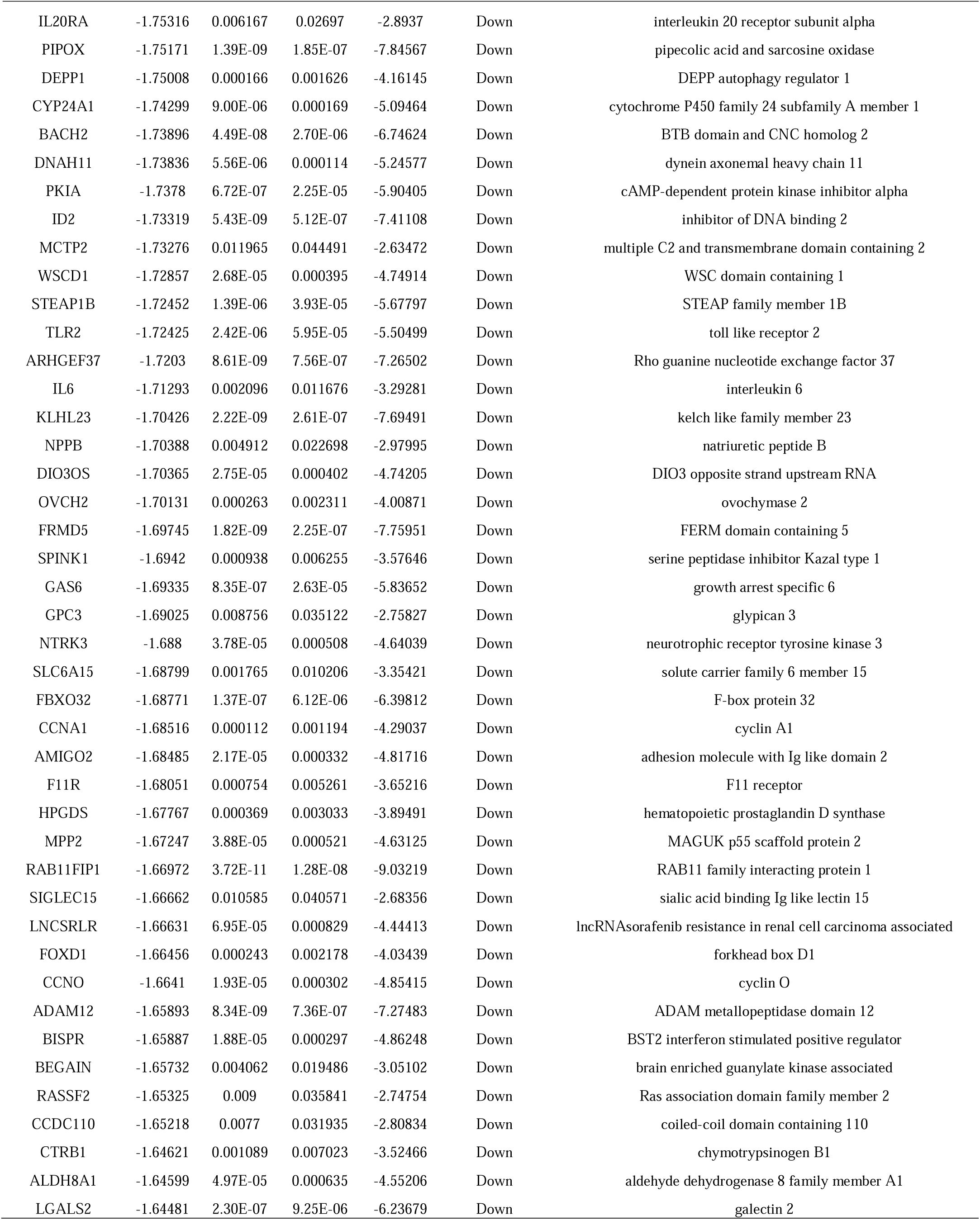
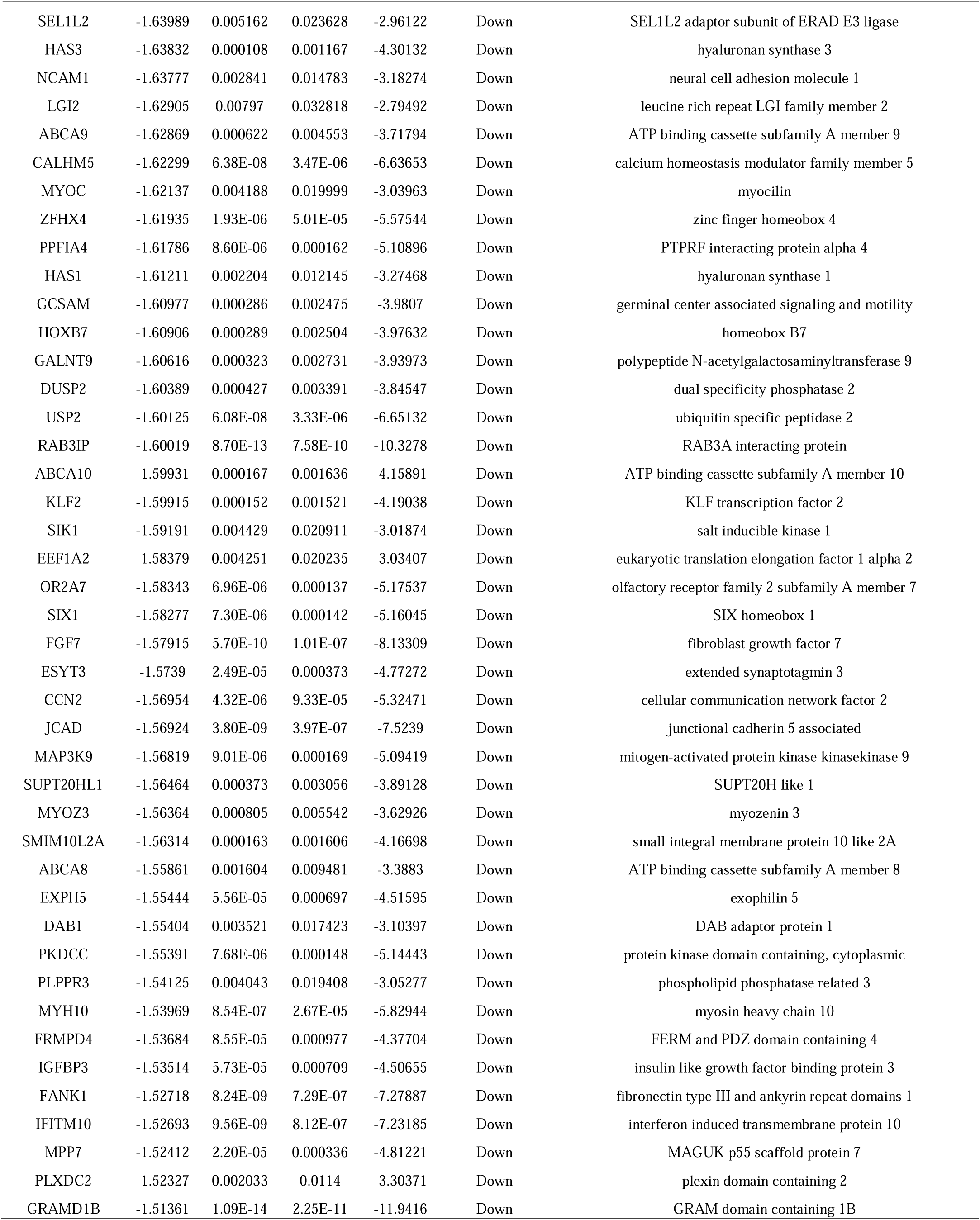
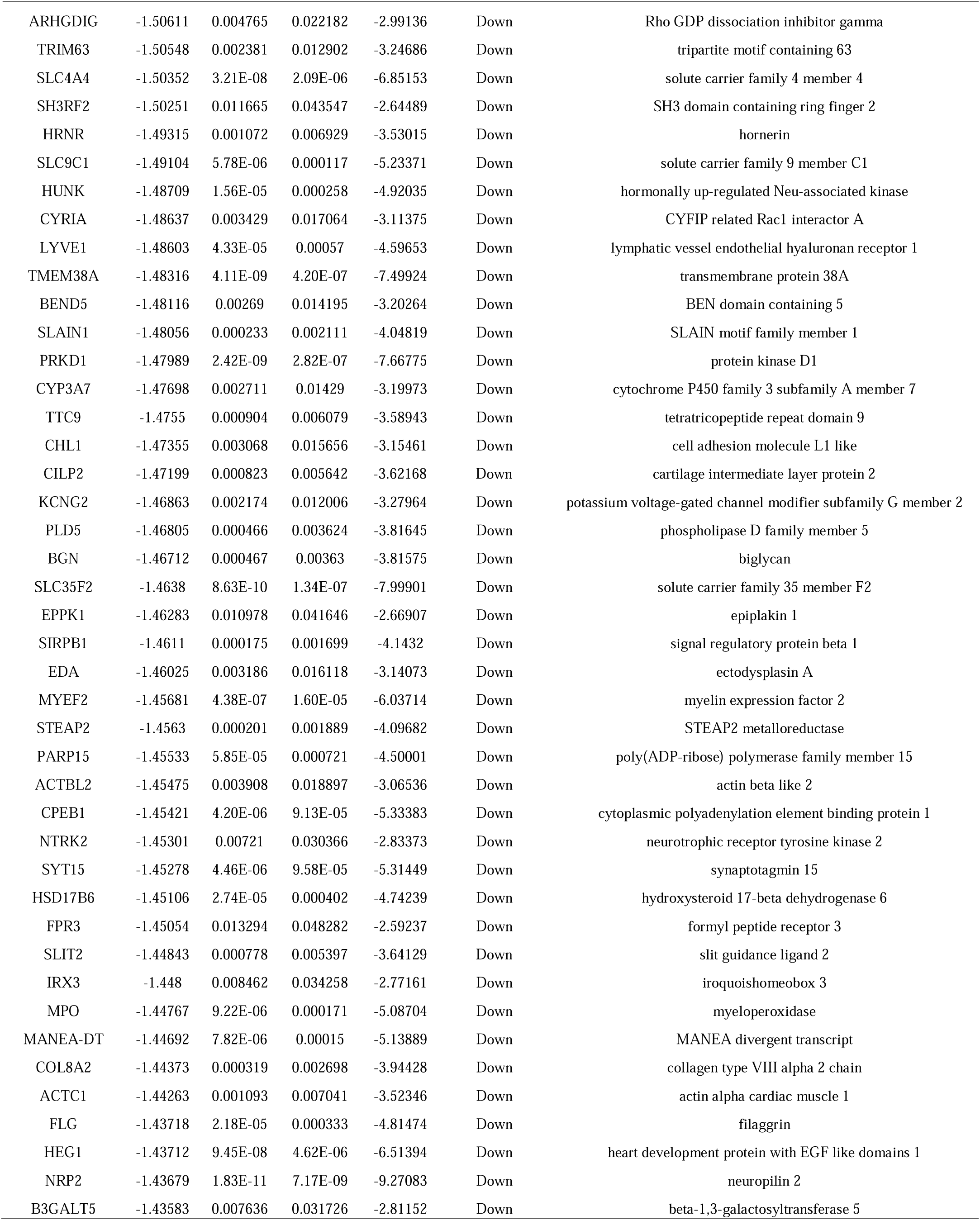
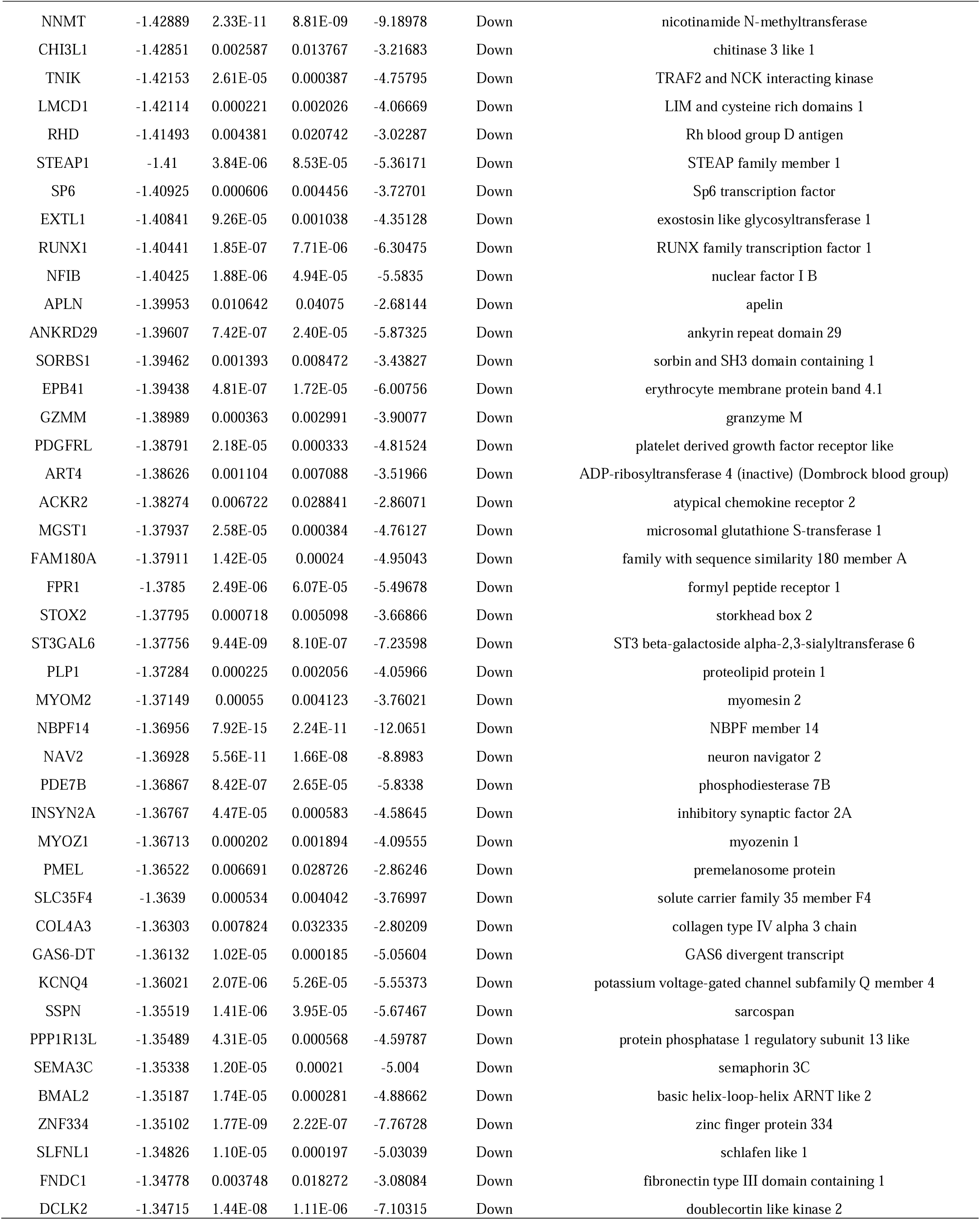
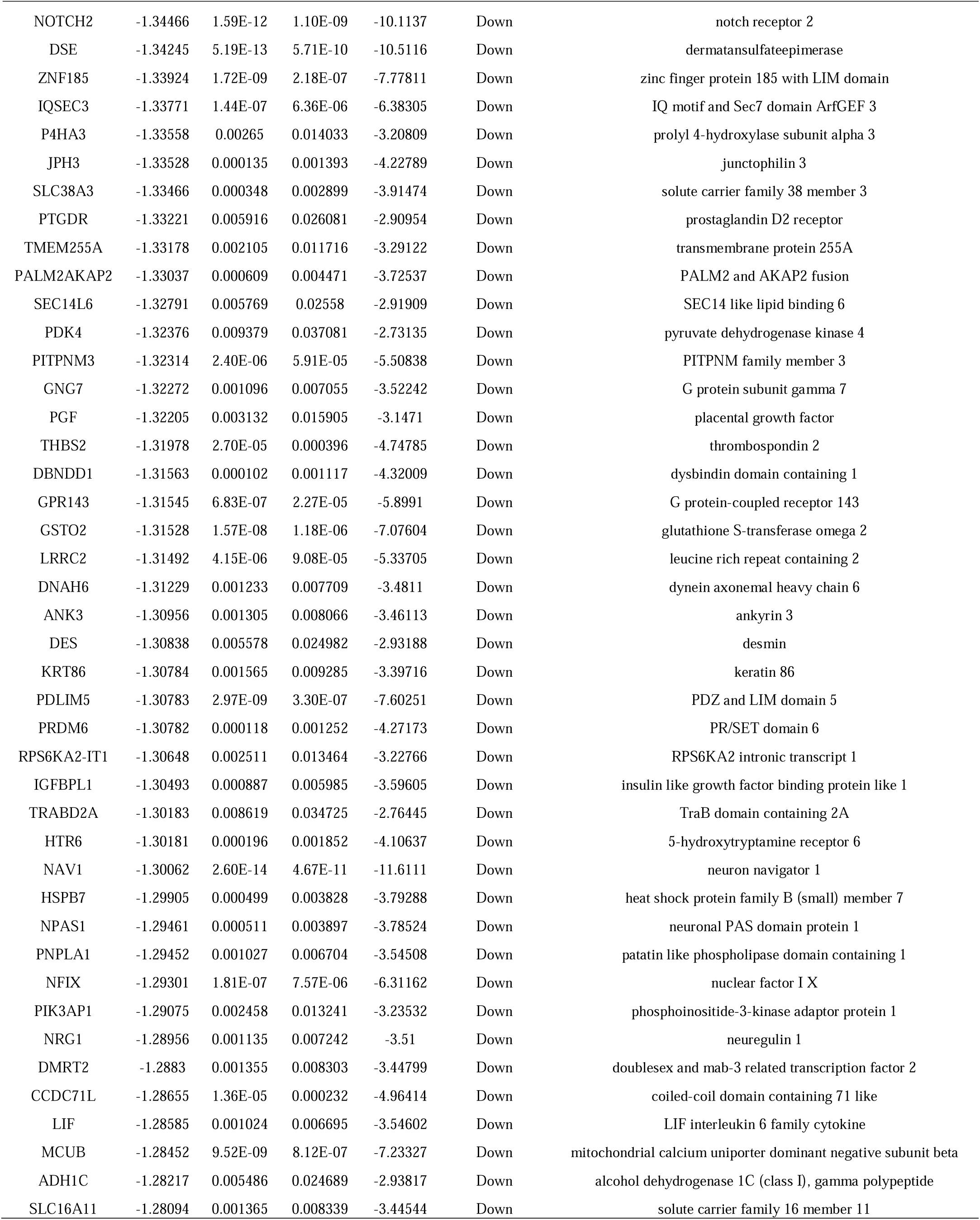
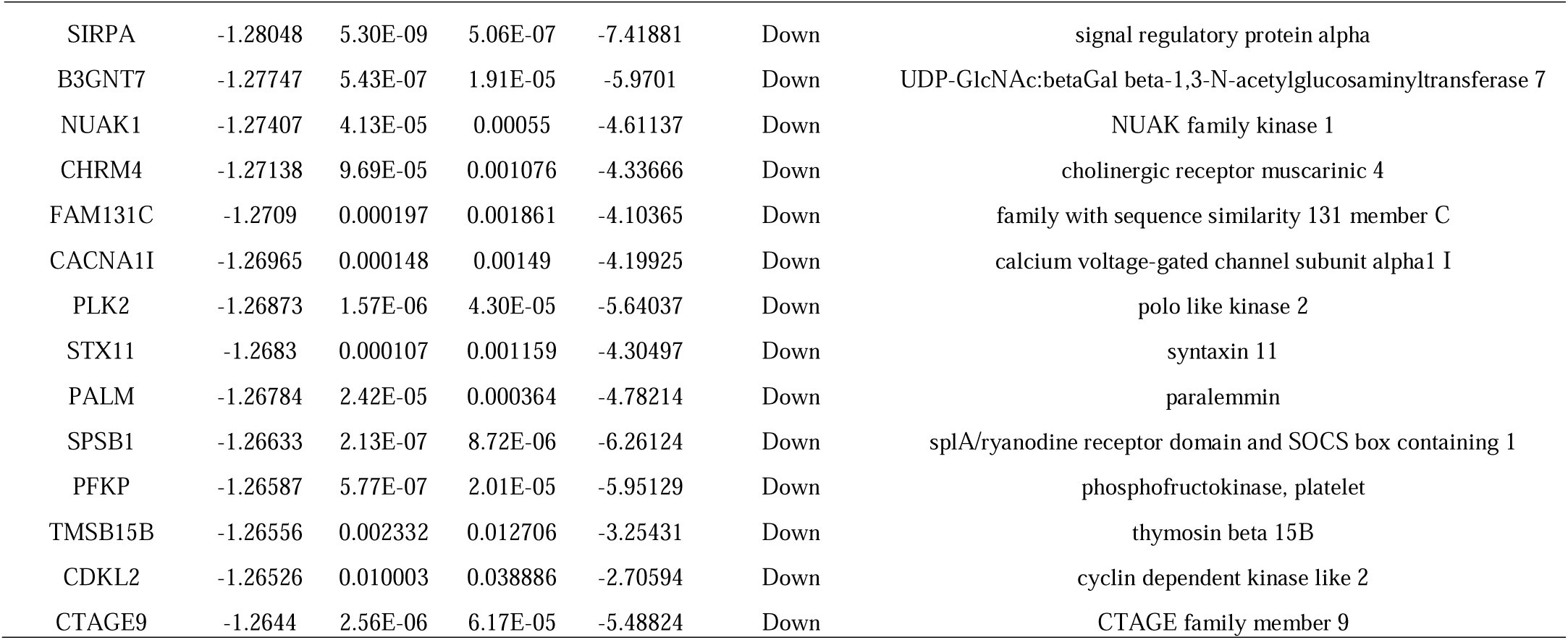
The statistical metrics for key differentially expressed genes (DEGs)

### GO and pathway enrichment analyses of DEGs

GO enrichment and REACTOME pathway enrichment analysis were performed on the DEGs using the g:Profiler database. GO enrichment analysis covers three aspects: BP, CC and MF (Table 2). The up regulated genes were mainly related to multicellular organismal process, regulation of biological process, membrane, extracellular region, signaling receptor binding and molecular transducer activity ; while the down regulated genes were mainly involved in developmental process, biological regulation, cell periphery, cytoplasm, molecular function regulator activity and calcium ion binding. The REACTOME pathway enrichment analysis showed that the genes up regulated genes in endometriosis were enriched in signaling by GPCR, extracellular matrix organization, muscle contraction and glycosaminoglycan metabolism (Table 3).

**Table 2.**
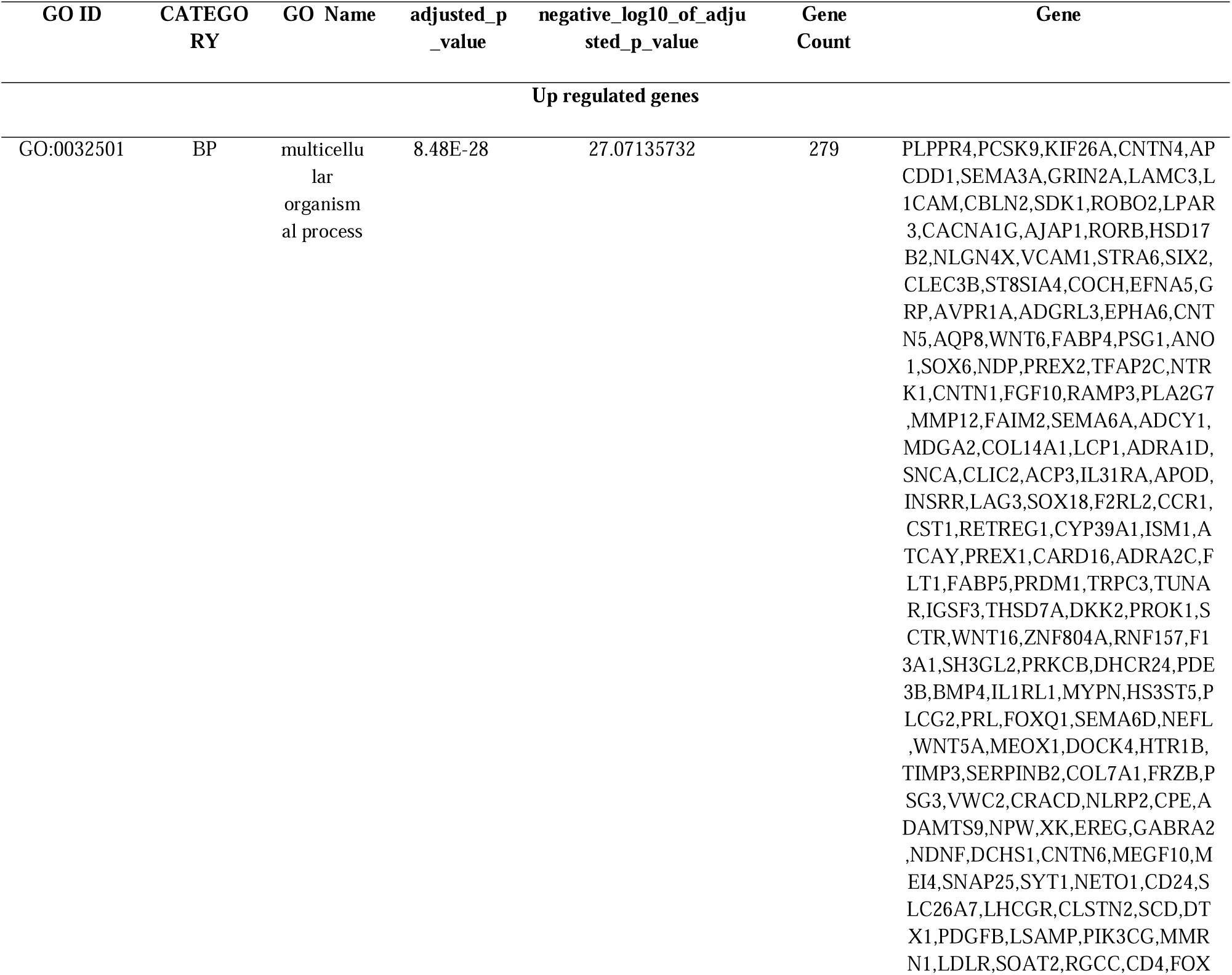
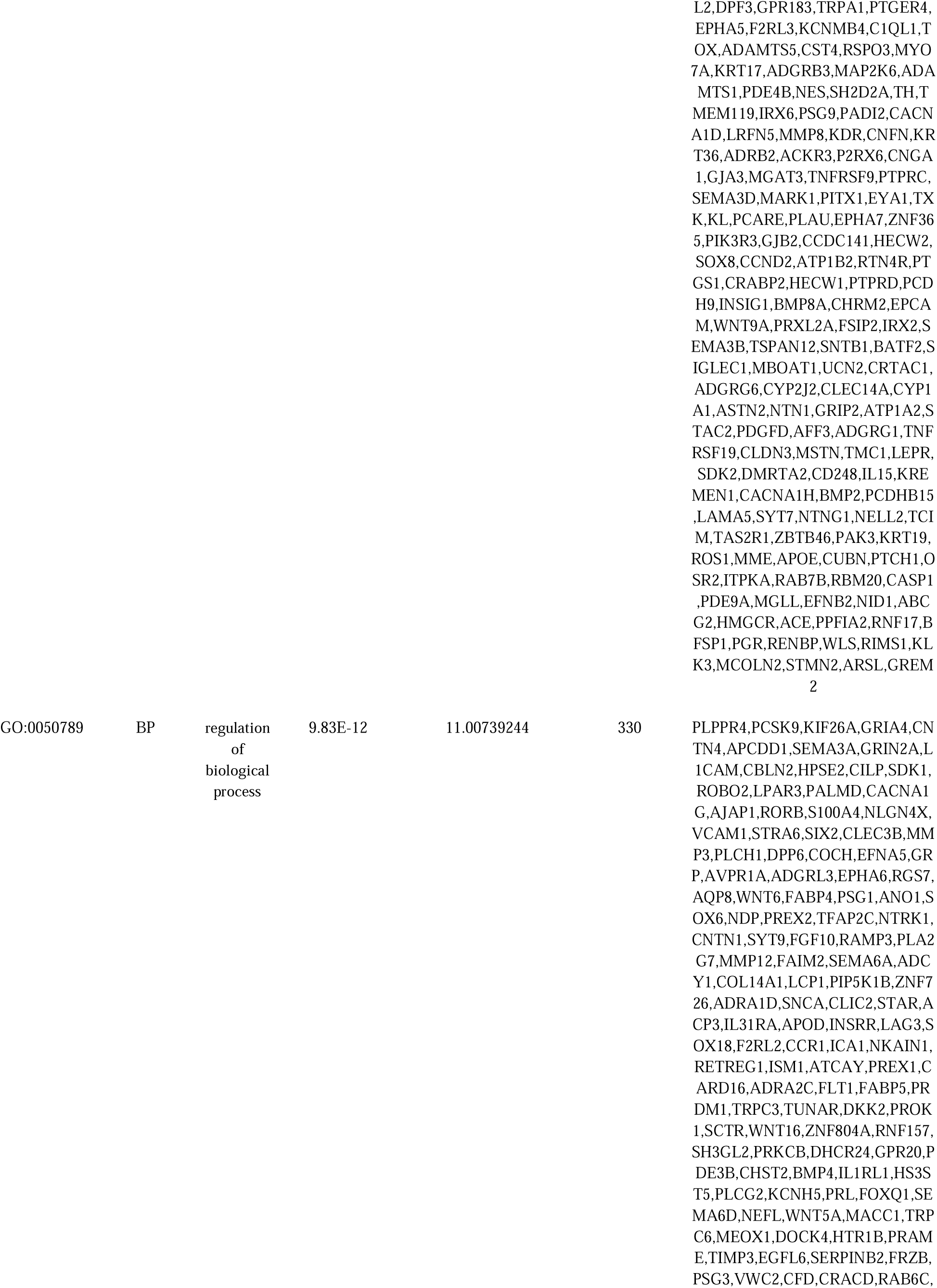
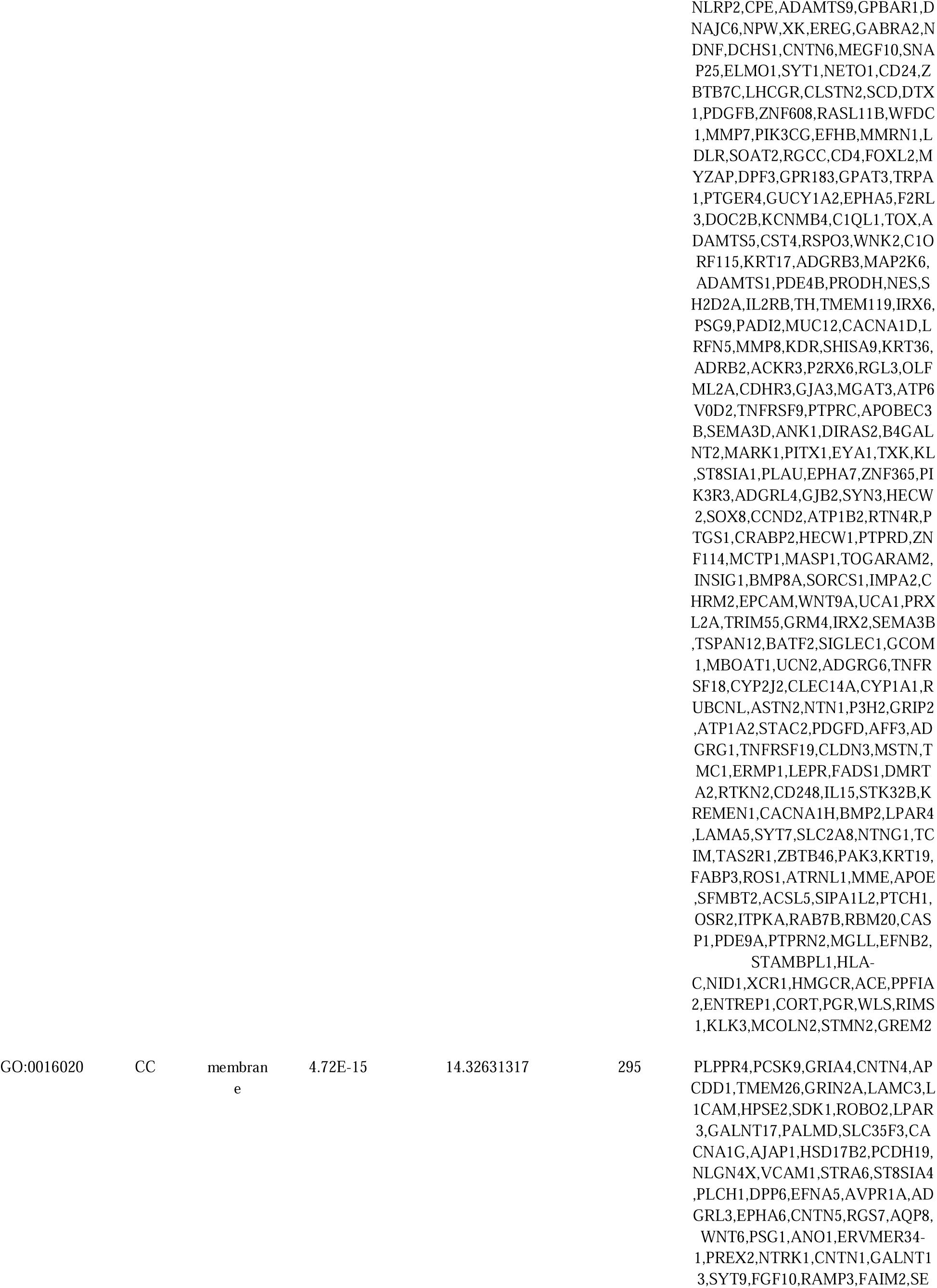
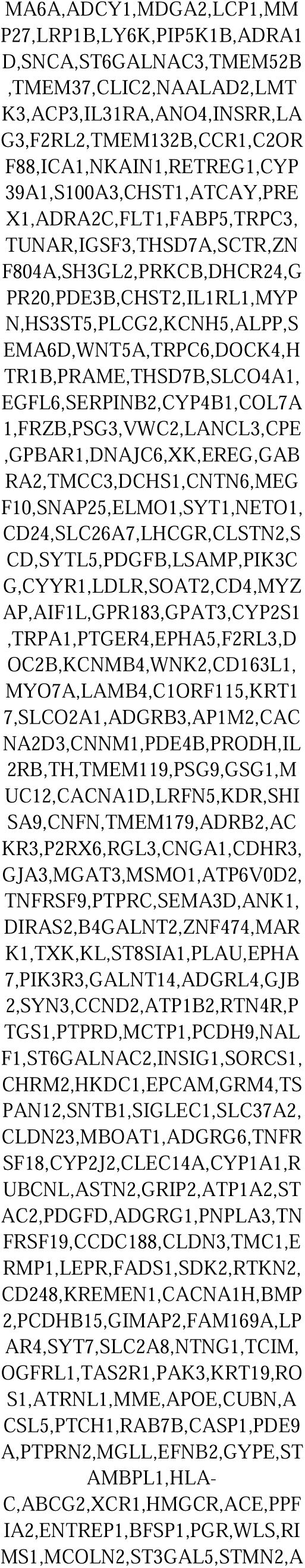
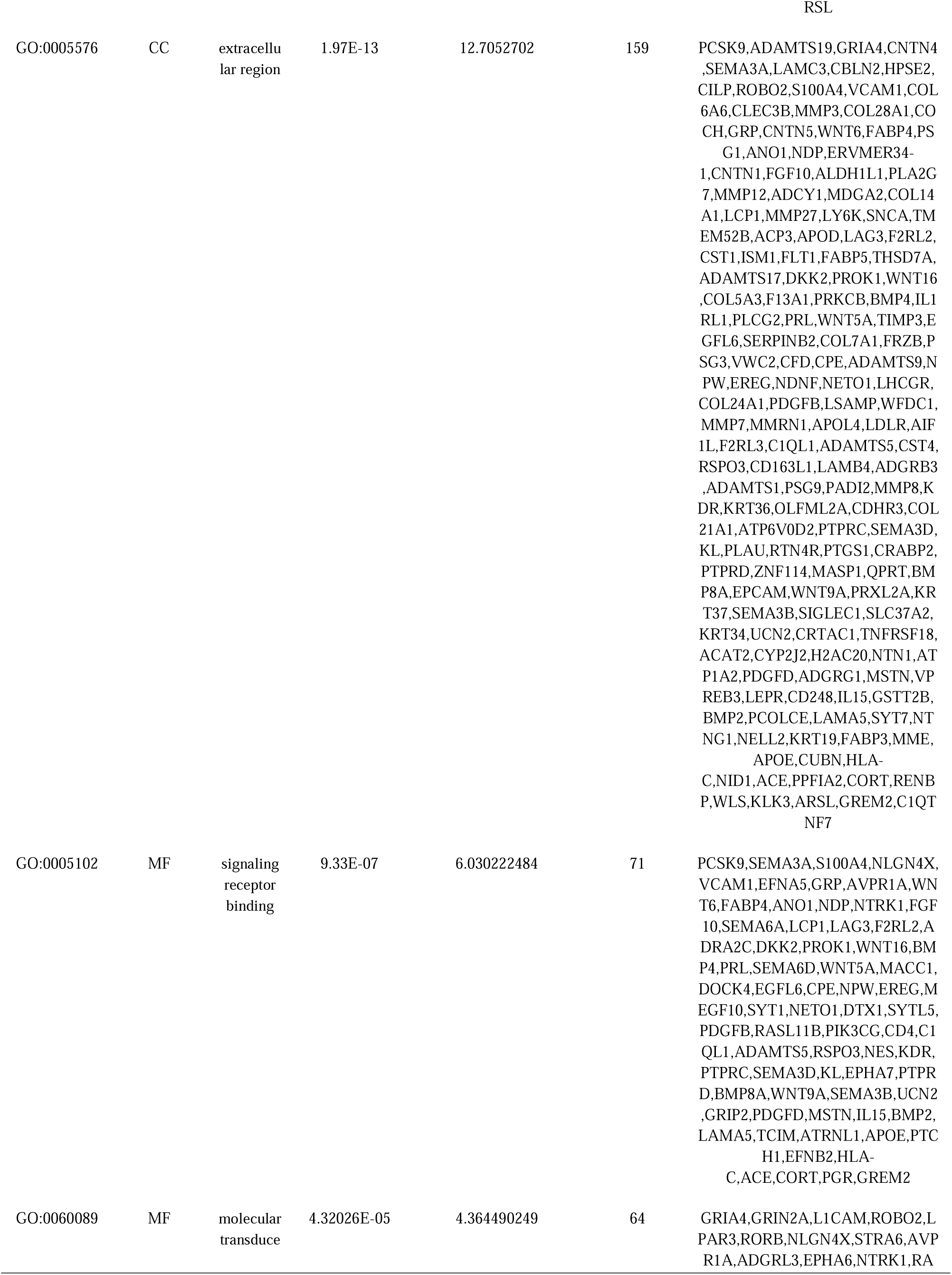
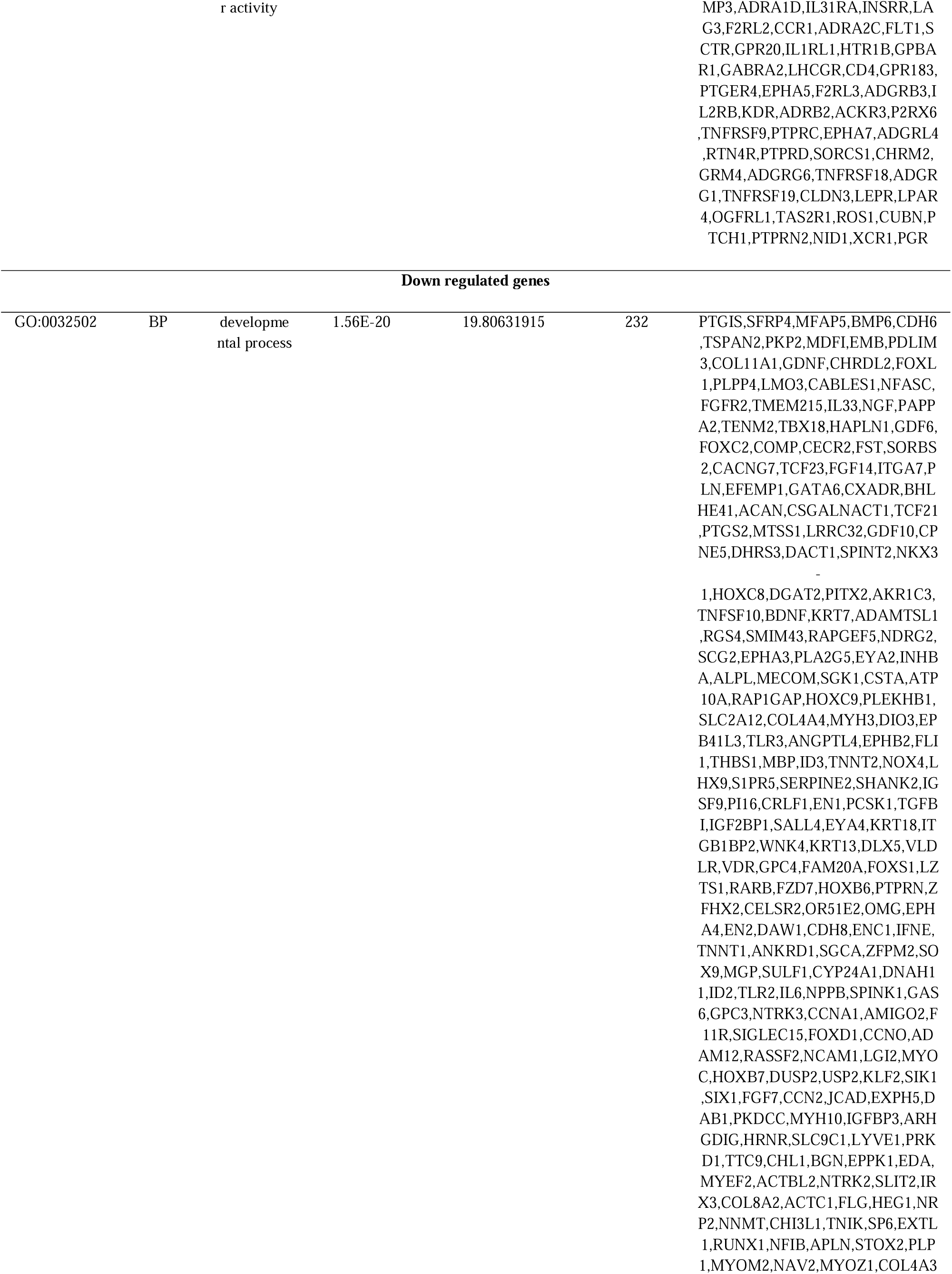
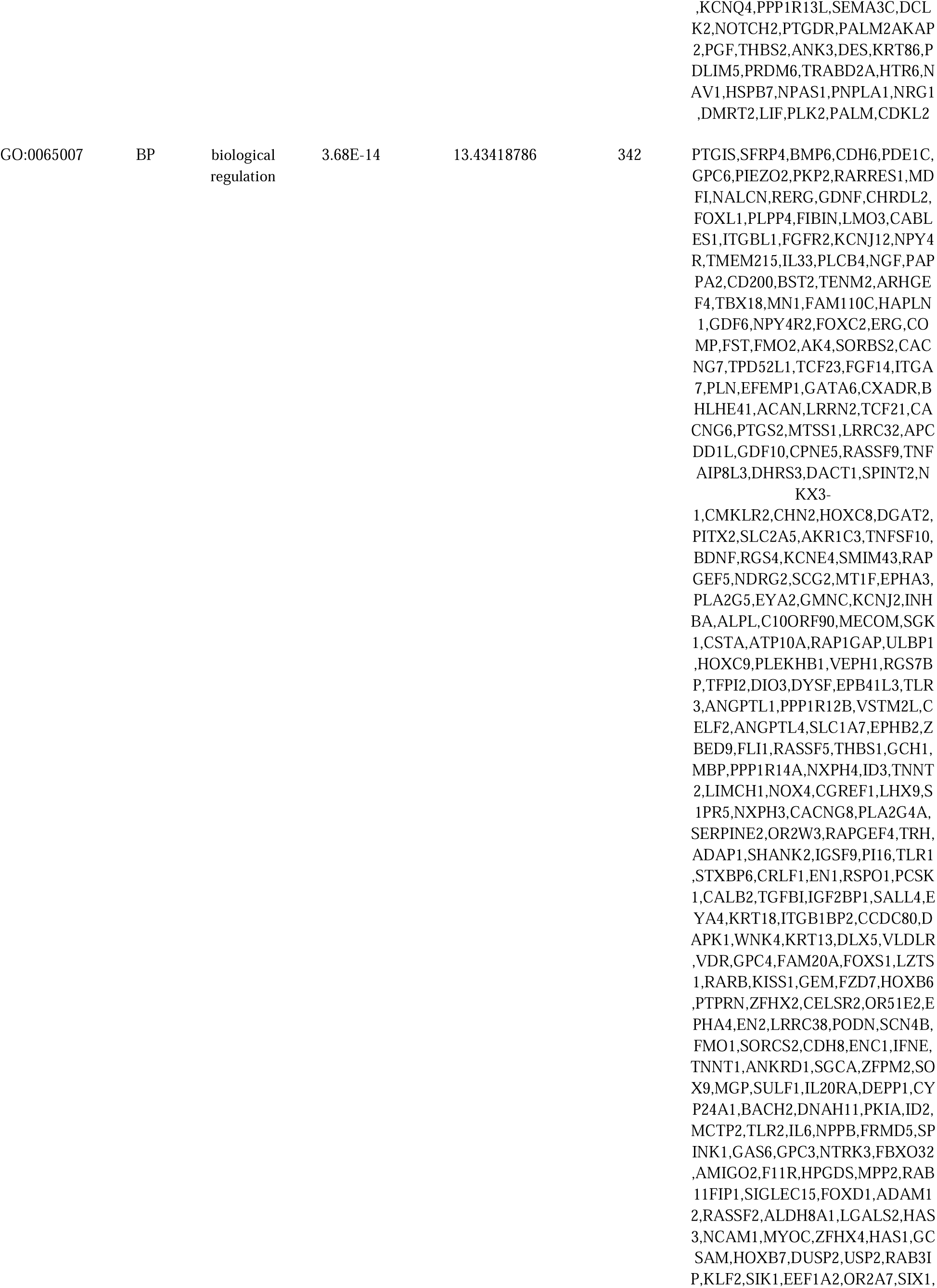
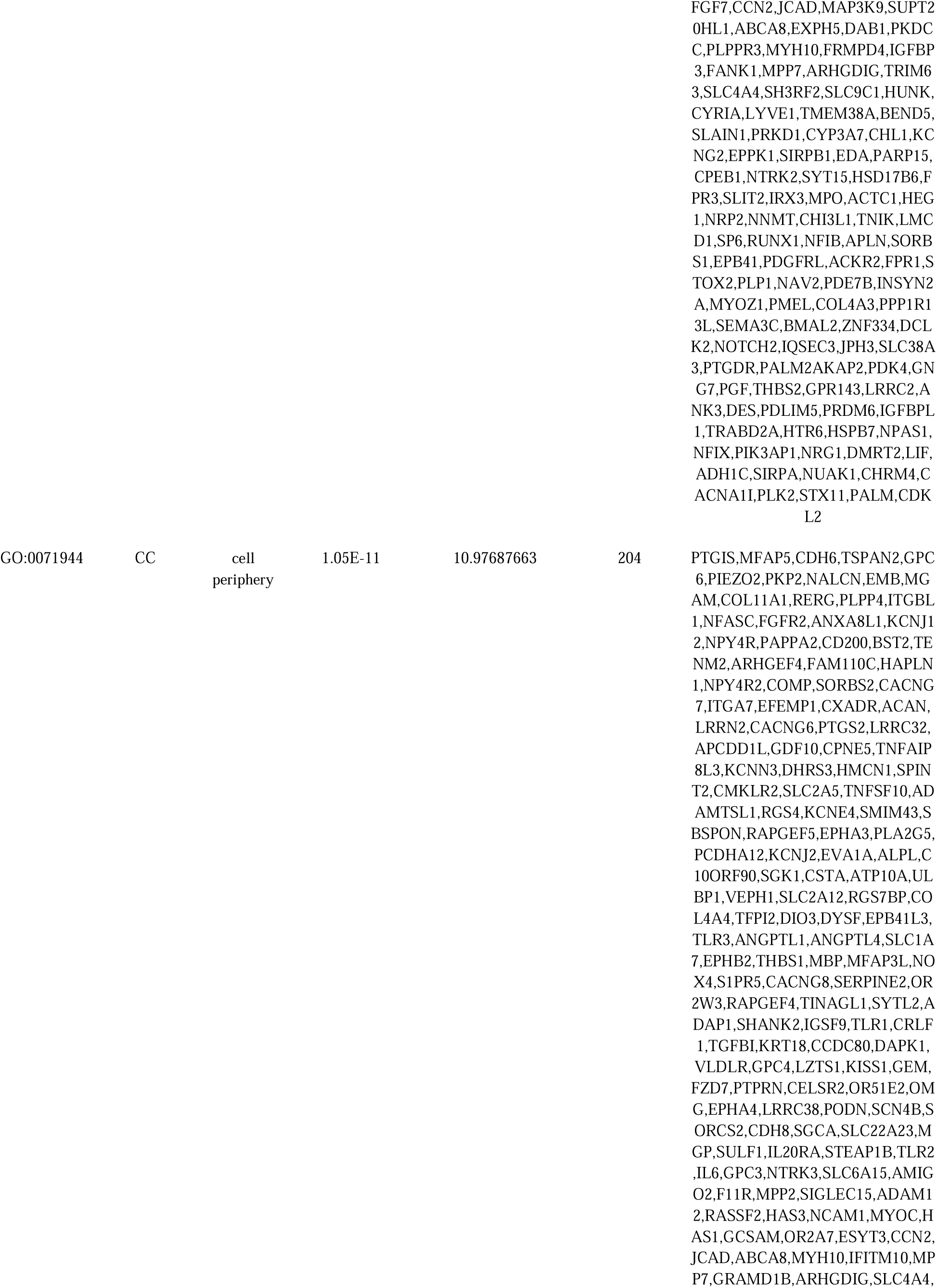
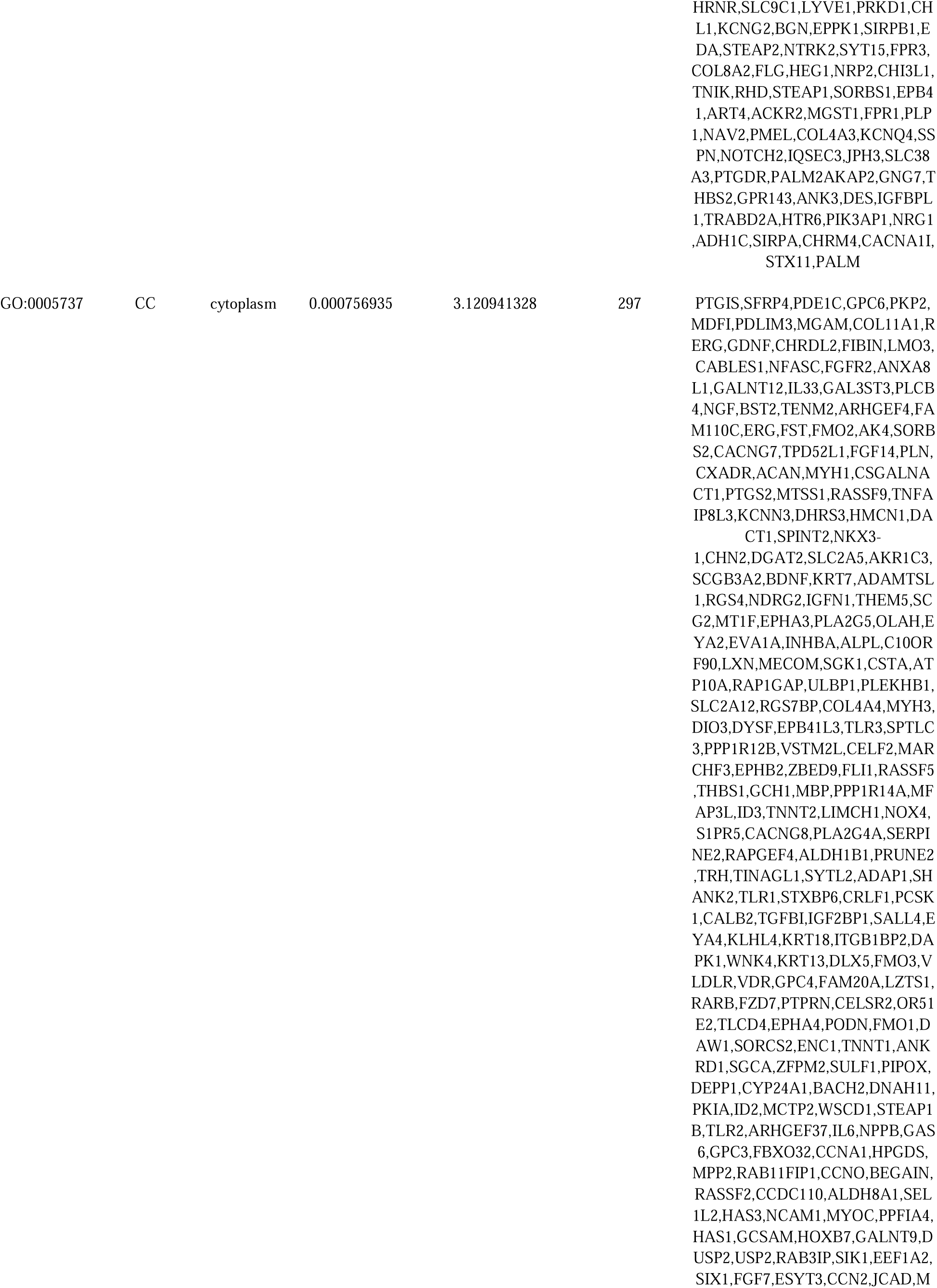
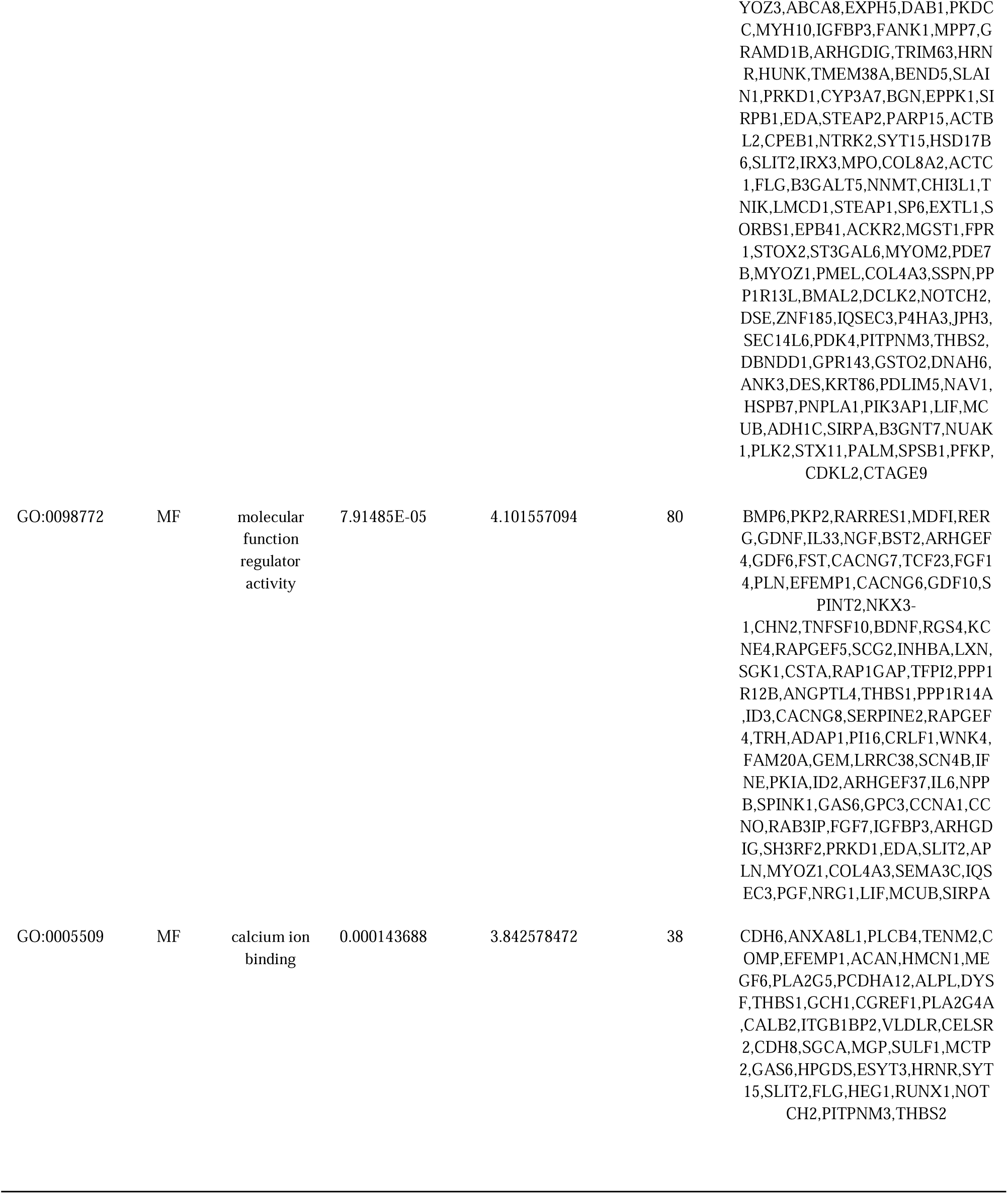
The enriched GO terms of the up and down regulated differentially expressed genes.

**Table 3.**
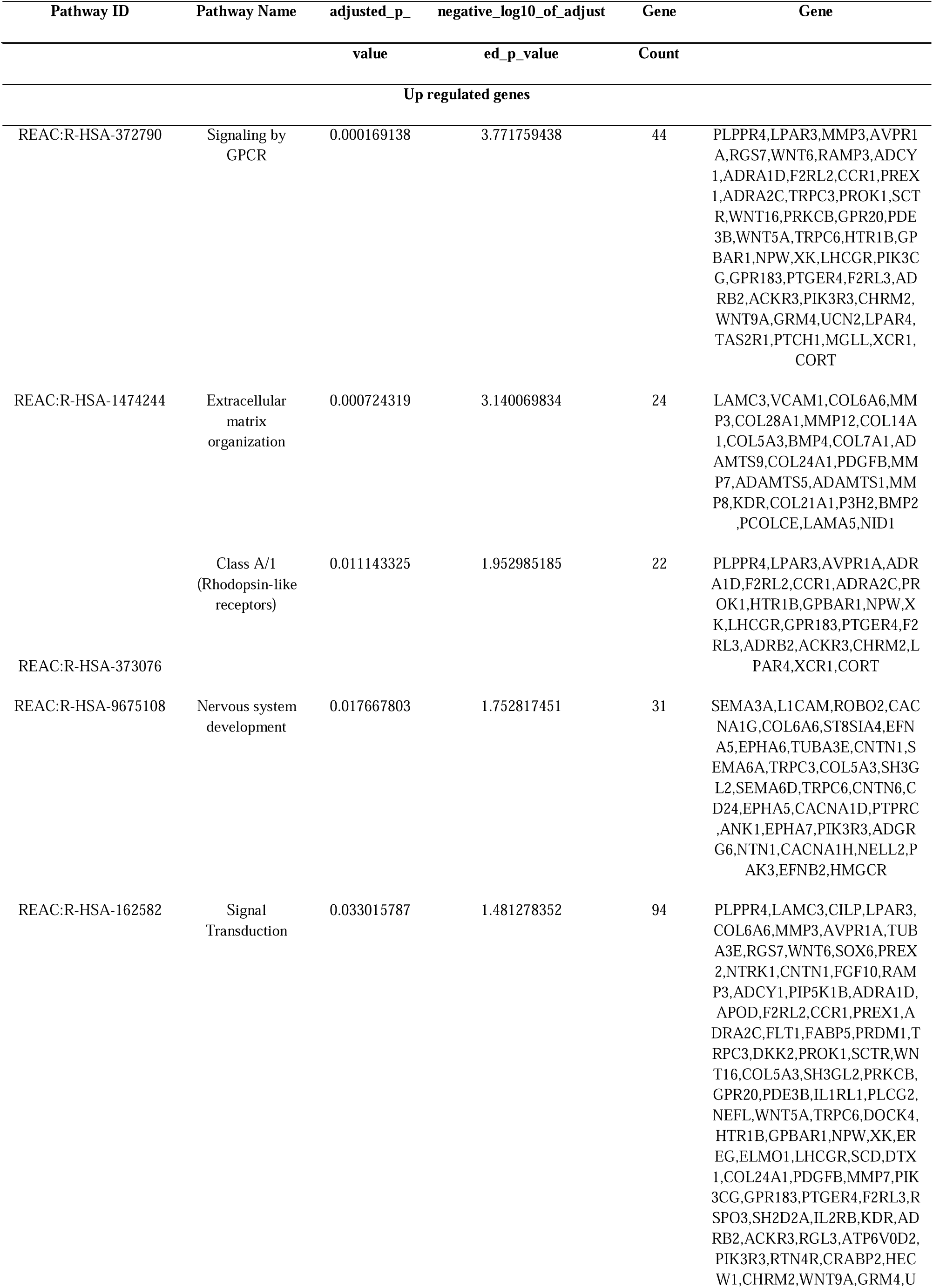
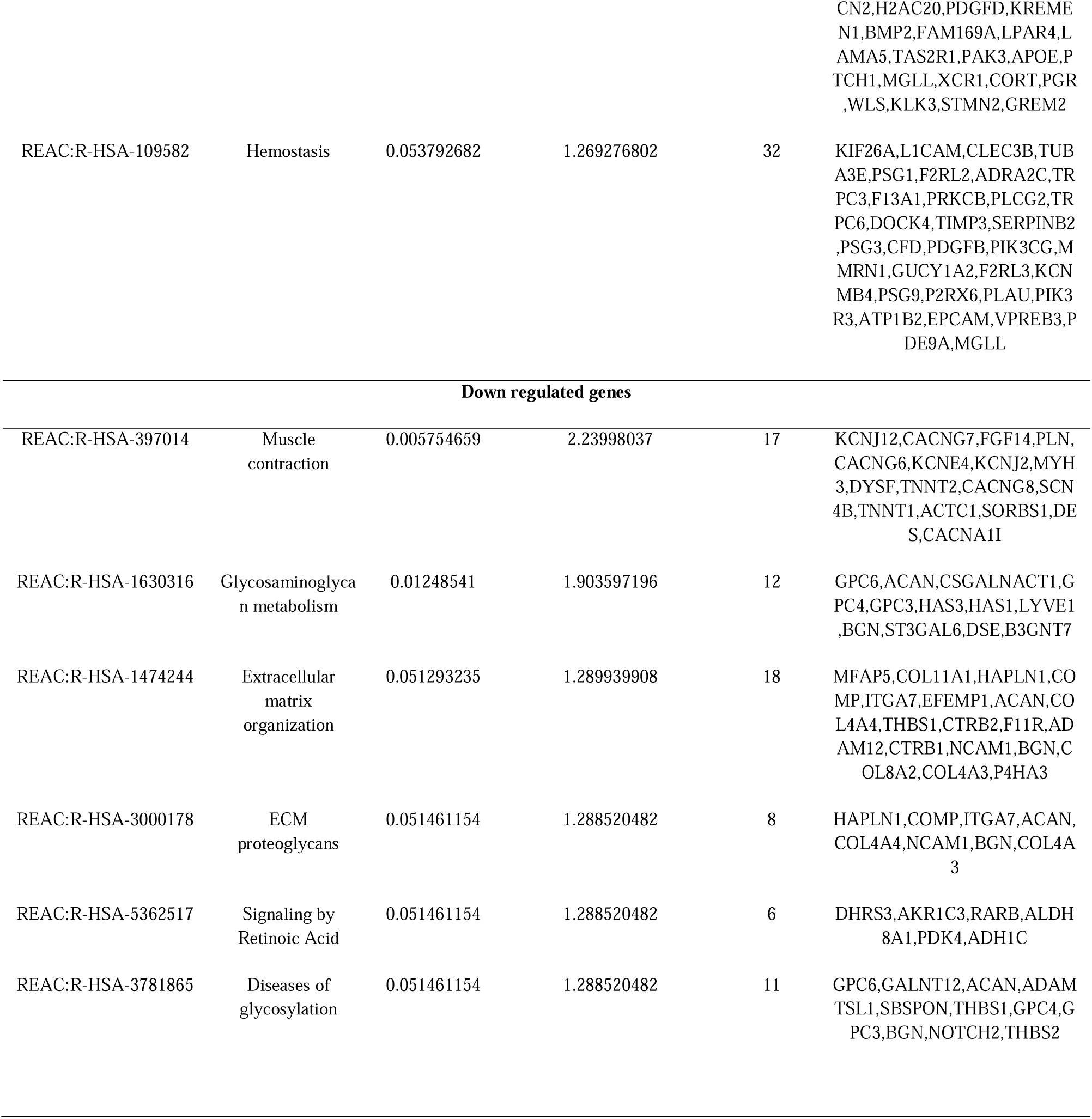
The enriched pathway terms of the up and down regulated differentially expressed genes.

### Construction of the PPI network and module analysis

Considering the critical role of protein interactions in protein function, we used the HIPIE database and Cytoscape software to generate PPI network once we had identified the 958 DEGs. The results showed that there were dense regions in PPI, that is, genes closely related to endometriosis. A total of 4871 nodes and 8009 edges were selected to plot the PPI network (Fig. 3). The Network Analyzer plugin of Cytoscape was used to score each node gene by 4 selected algorithms, including node degree, betweenness, stress and closeness. Finally, we identified ten hub genes (VCAM1, SNCA, PRKCB, ADRB2, FOXQ1, MDFI, ACTBL2, PRKD1, DAPK1 and ACTC1) and are listed Table 4. The top two significant modules from PEWCC were selected for future analysis. Module 1 included 22 nodes and 41 edges (Fig. 4A). The functional enrichment analysis of genes in module 1 were conducted by g:Profiler. These genes were significantly enriched in multicellular organismal process and regulation of biological process. Module 2 included 8 nodes and 14 edges (Fig. 4B). The functional enrichment analysis of genes in module 2 were conducted by g:Profiler. These genes were significantly enriched in developmental process and biological regulation.

**Fig. 3.**
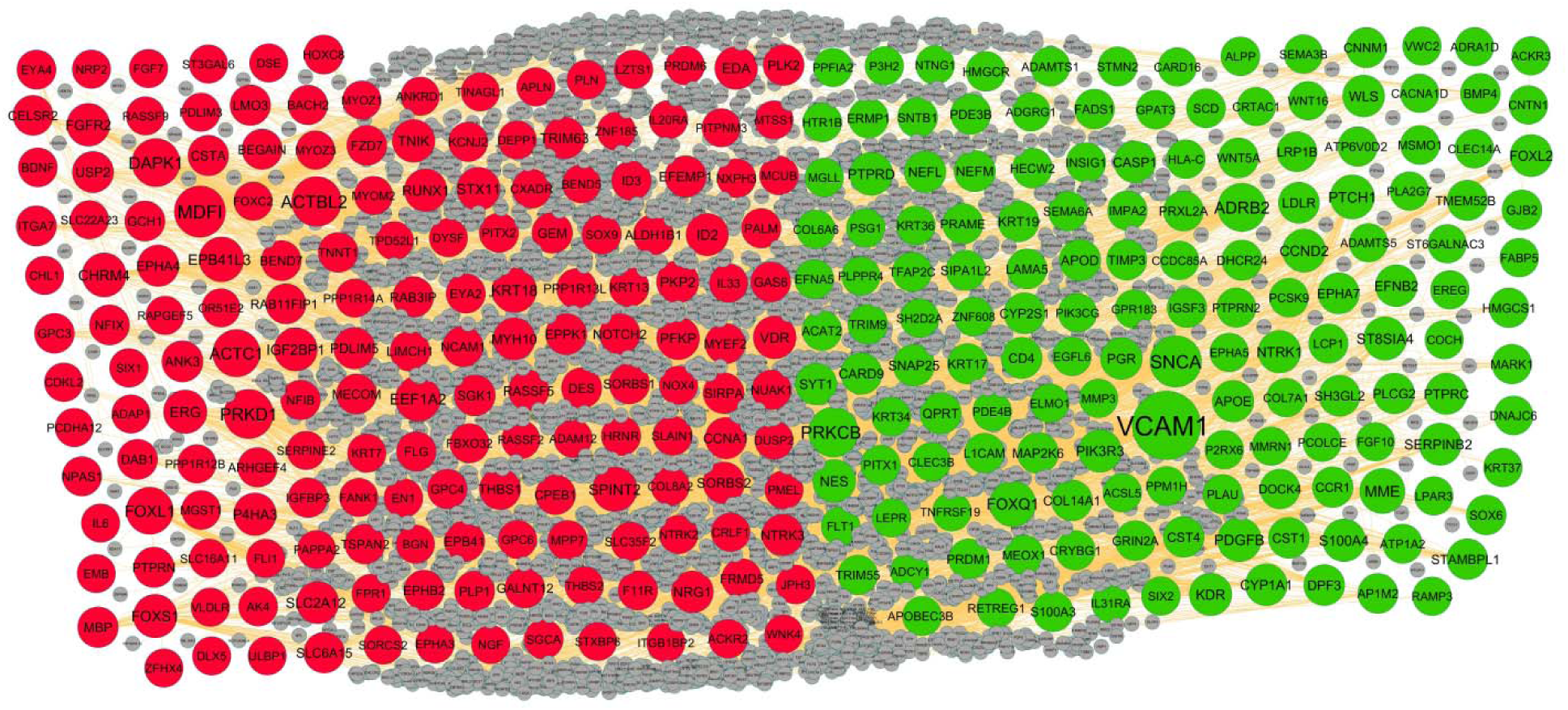
PPI network of DEGs. Up regulated genes are marked in parrot green; down regulated genes are marked in red.

**Fig. 4.**
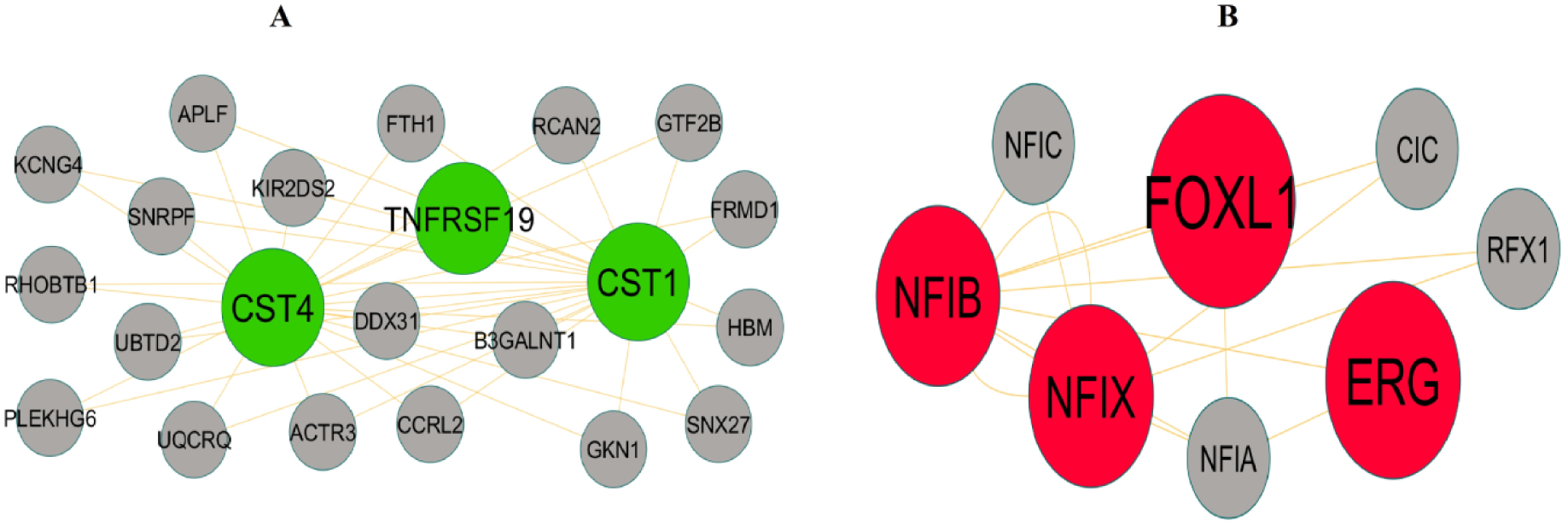
Modules selected from the PPI network. (A) The most significant module was obtained from PPI network with 22 nodes and 41 edges for up regulated genes (B) The most significant module was obtained from PPI network with 8 nodes and 14 edges for down regulated genes. Up regulated genes are marked in parrot green; down regulated genes are marked in red.

**Table 4.**
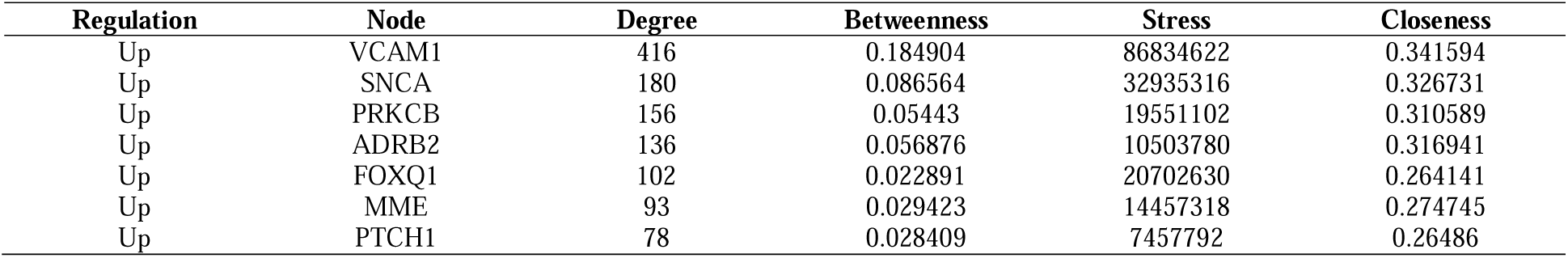
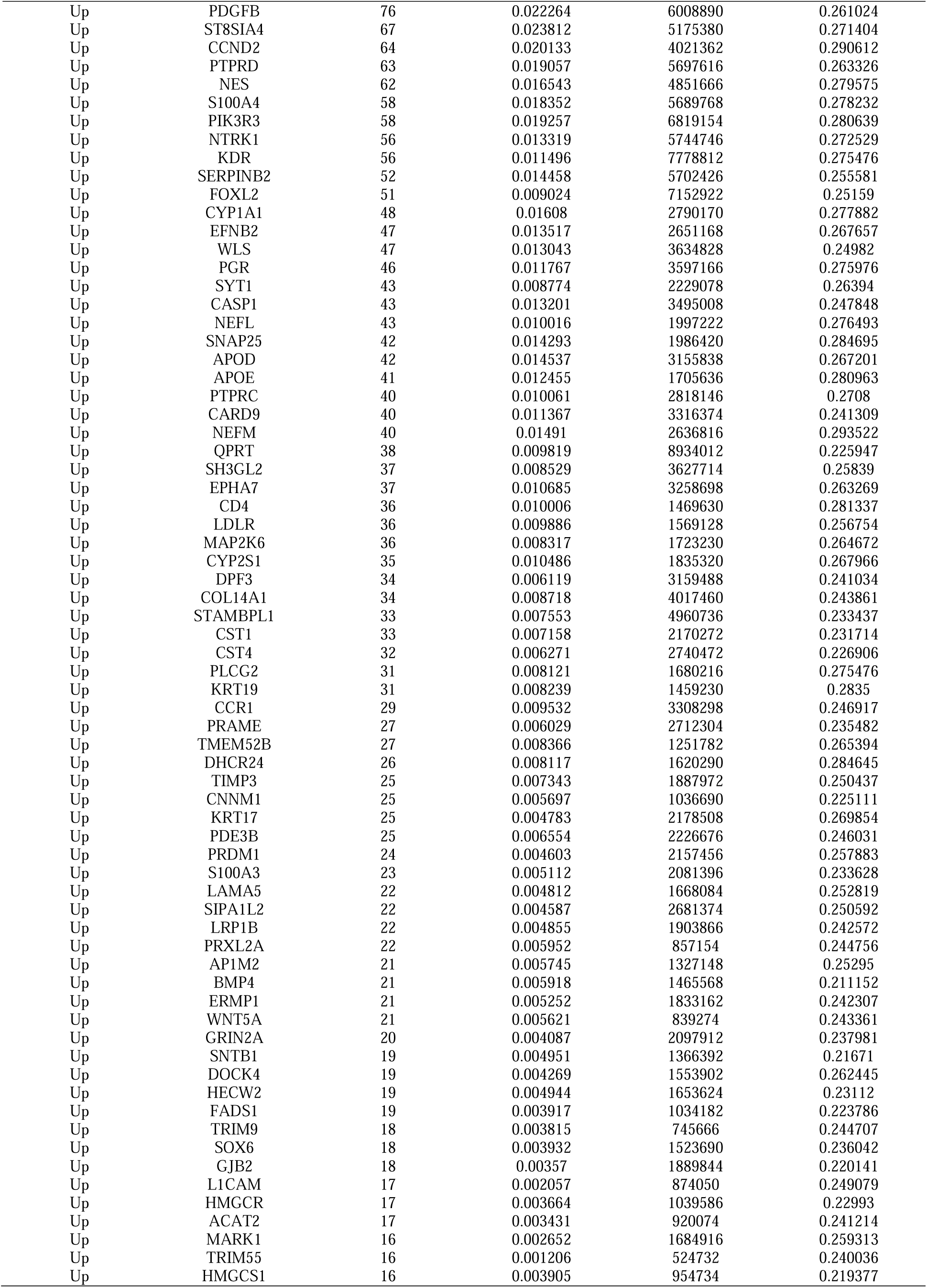
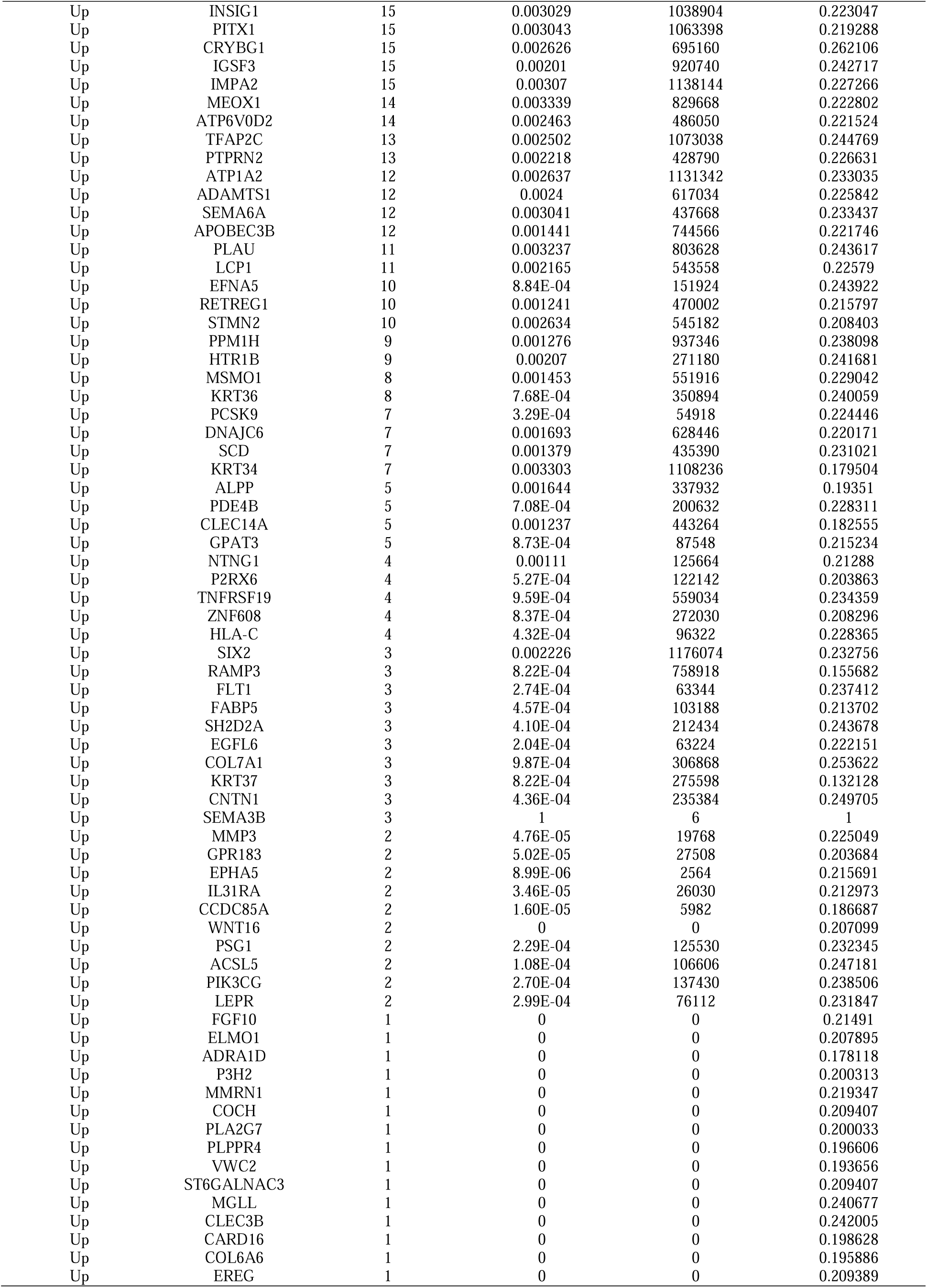
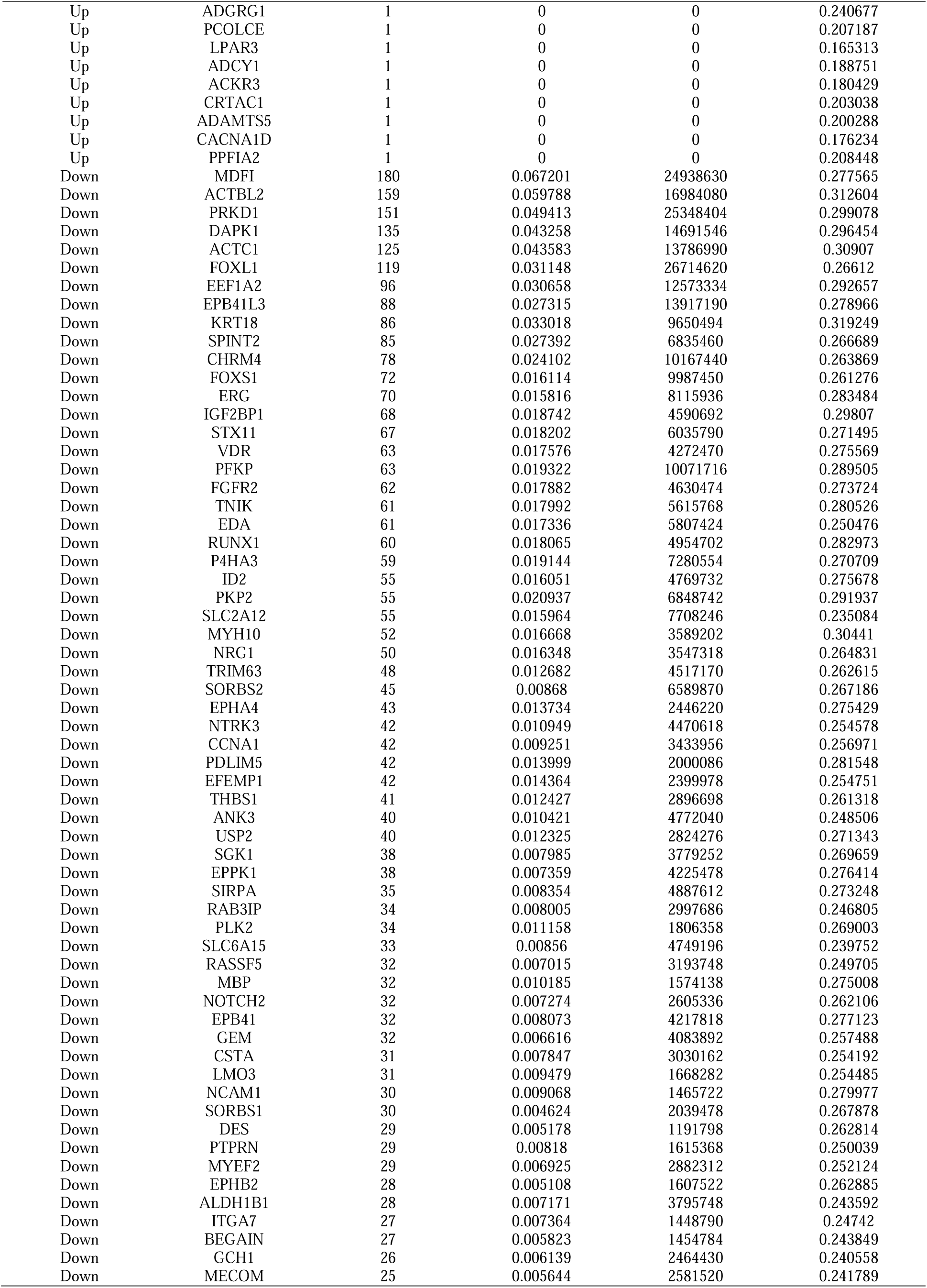
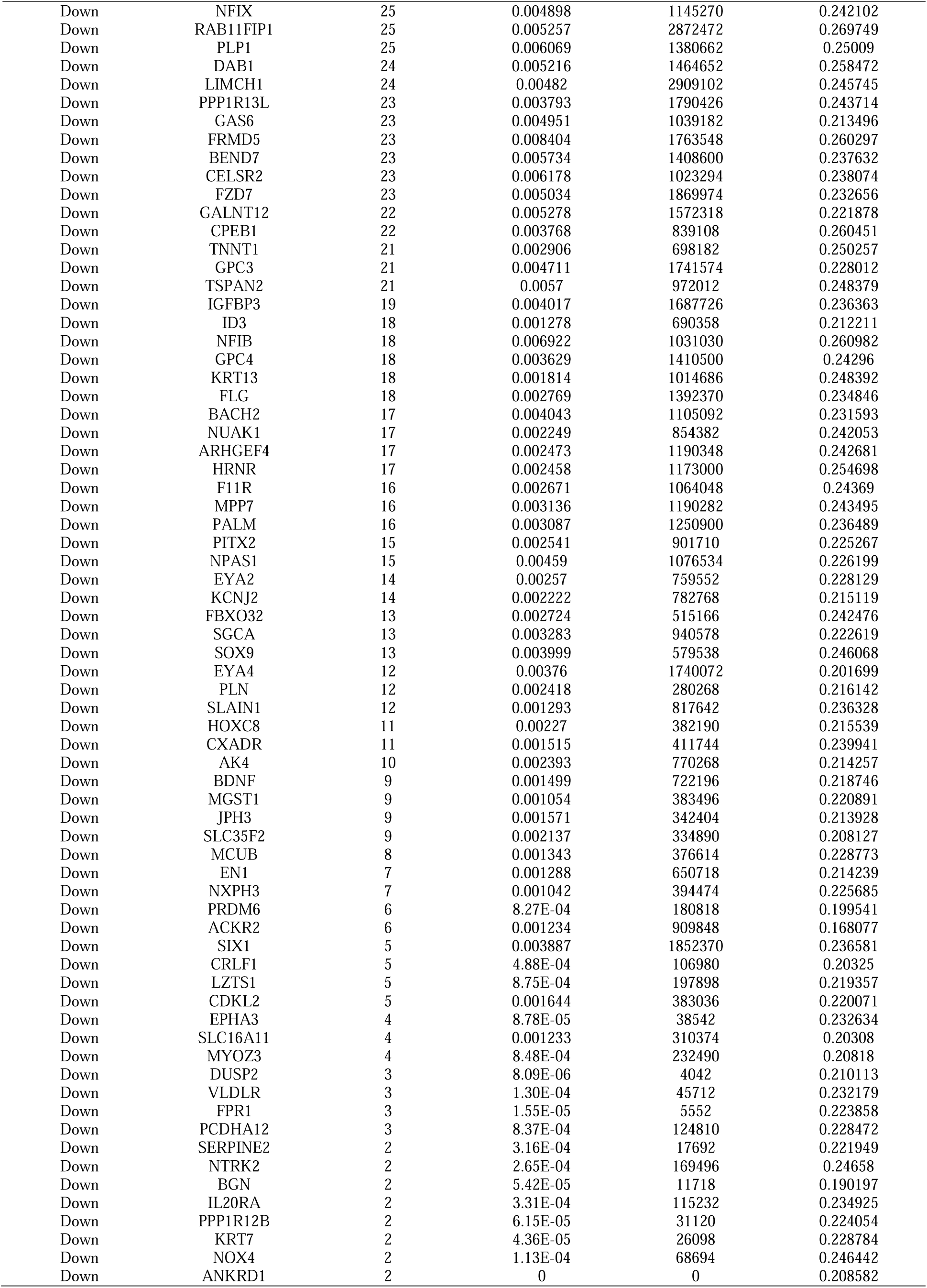
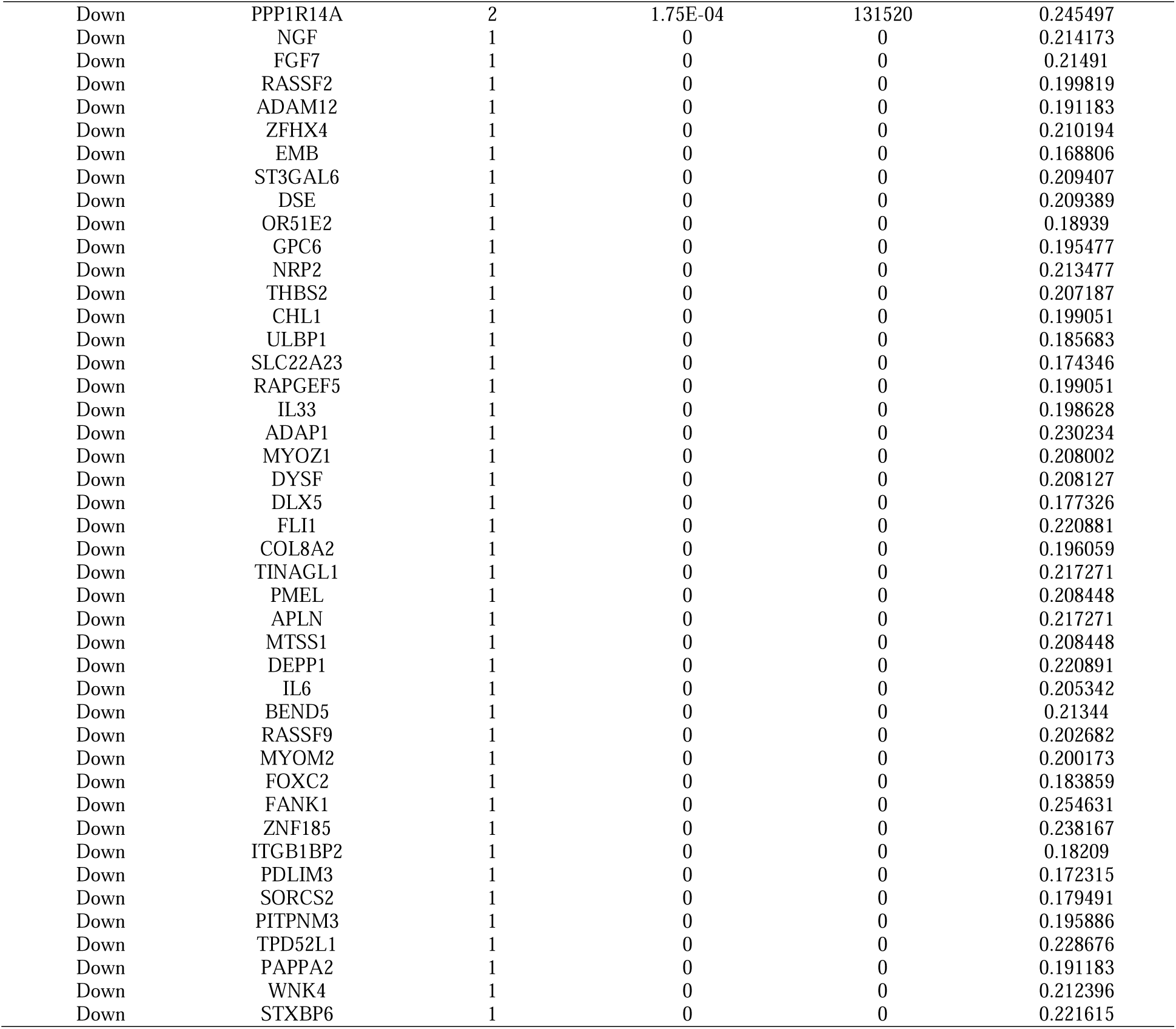
Topology table for up and down regulated genes.

### Construction of the miRNA-hub gene regulatory network

We searched for target-regulated hub gene miRNAs using miRNet database and then used the results of this database. By constructing miRNA-hub gene regulatory network networks, we found 2495 nodes (miRNA: 2168; Hub Gene: 327) and 14692 edges (Fig. 5). We identified 365 miRNAs (ex; hsa-mir-3143) targeting regulation of CCND2, 102 miRNAs (ex; hsa-mir-6888-5p) targeting regulation of VCAM1, 89 miRNAs (ex; hsa-mir-200a-3p) targeting regulation of PTPRD, 88 miRNAs (ex; hsa-mir-3122) targeting regulation of PDGFB, 81 miRNAs (ex; hsa-mir-17-5p) targeting regulation of PRKCB, 241 miRNAs (ex; hsa-mir-2110) targeting regulation of IGF2BP1, 77 miRNAs (ex; hsa-mir-4432) targeting regulation of ACTC1, 53 miRNAs (ex; hsa-mir-556-3p) targeting regulation of EPB41L3, 48 miRNAs (ex; hsa-mir-10b-5p) targeting regulation of DAPK1 and 41 miRNAs (ex; hsa-mir-1229-5p) targeting regulation of MDFI, and are listed Table 5.

**Fig. 5.**
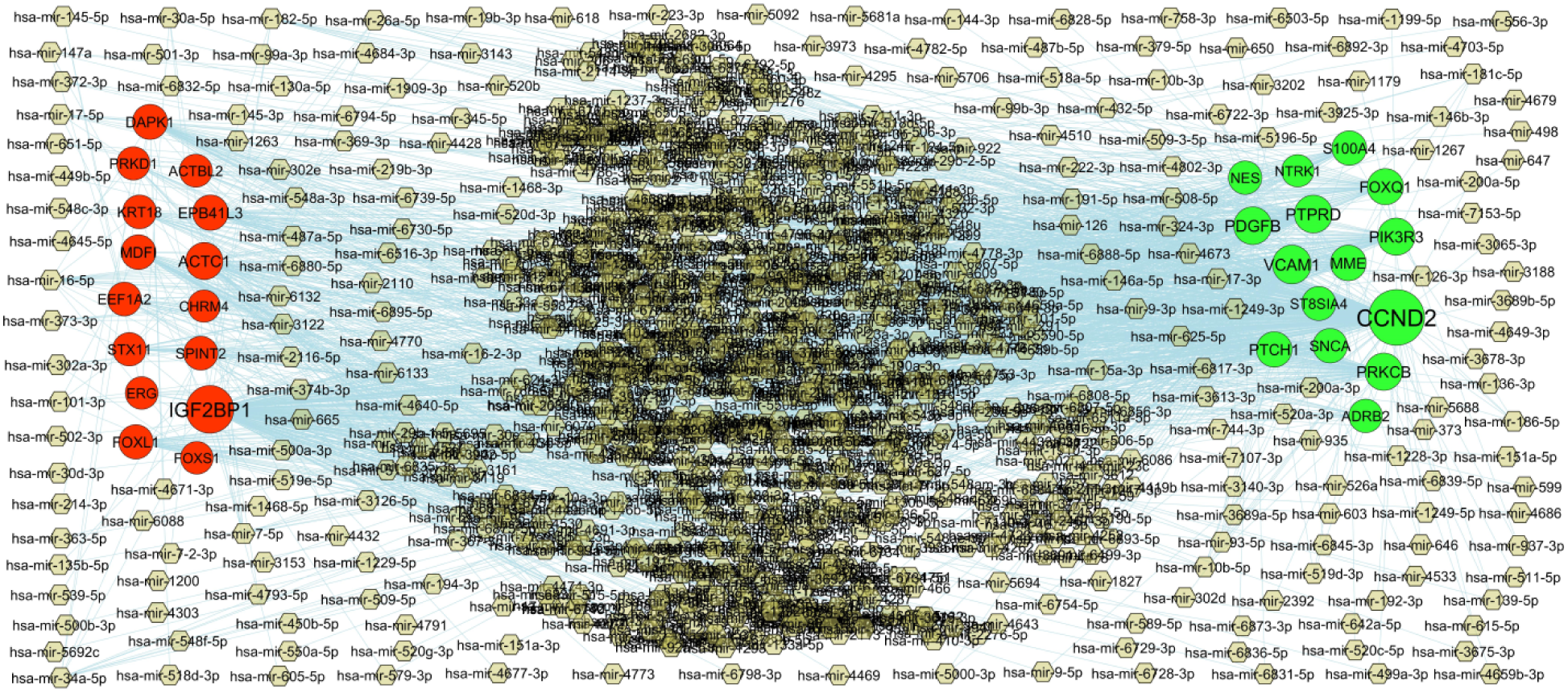
Hub gene - miRNA regulatory network. The olive green color diamond nodes represent the key miRNAs; up regulated genes are marked in green; down regulated genes are marked in red.

**Table 5.** MiRNA - hub gene and TF – hub gene topology table.

| Regulation | Hub Genes | Degree | MicroRNA | Regulation | Hub Genes | Degree | TF |
| --- | --- | --- | --- | --- | --- | --- | --- |
| Up | CCND2 | 365 | hsa-mir-3143 | Up | PTCH1 | 59 | TCF3 |
| Up | VCAM1 | 102 | hsa-mir-6888-5p | Up | CCND2 | 42 | PHC1 |
| Up | PTPRD | 89 | hsa-mir-200a-3p | Up | PRKCB | 37 | NR1I2 |
| Up | PDGFB | 88 | hsa-mir-3122 | Up | ST8SIA4 | 36 | HOXC9 |
| Up | PRKCB | 81 | hsa-mir-17-5p | Up | FOXQ1 | 34 | RNF2 |
| Up | PIK3R3 | 78 | hsa-mir-93-5p | Up | PDGFB | 34 | RELA |
| Up | FOXQ1 | 62 | hsa-mir-5706 | Up | NES | 32 | KLF4 |
| Up | MME | 54 | hsa-mir-186-5p | Up | MME | 31 | BMI1 |
| Up | PTCH1 | 52 | hsa-mir-372-3p | Up | PIK3R3 | 30 | RUNX1 |
| Up | SNCA | 37 | hsa-mir-369-3p | Up | PTPRD | 27 | PAX3 |
| Up | S100A4 | 36 | hsa-mir-194-3p | Up | ADRB2 | 27 | GATA2 |
| Up | ST8SIA4 | 31 | hsa-mir-181c-5p | Up | SNCA | 25 | SCLY |
| Up | ADRB2 | 22 | hsa-mir-10b-5p | Up | S100A4 | 20 | ARNT |
| Up | NES | 21 | hsa-mir-487b-5p | Up | VCAM1 | 16 | SMAD3 |
| Up | NTRK1 | 4 | hsa-mir-147a | Up | NTRK1 | 13 | JARID2 |
| Down | IGF2BP1 | 241 | hsa-mir-2110 | Down | DAPK1 | 44 | CLOCK |
| Down | ACTC1 | 77 | hsa-mir-4432 | Down | IGF2BP1 | 38 | PRDM14 |
| Down | EPB41L3 | 53 | hsa-mir-556-3p | Down | KRT18 | 36 | SMARCA4 |
| Down | DAPK1 | 48 | hsa-mir-10b-5p | Down | STX11 | 34 | TRIM28 |
| Down | MDFI | 41 | hsa-mir-1229-5p | Down | EPB41L3 | 31 | HTT |
| Down | FOXL1 | 36 | hsa-mir-7-5p | Down | FOXL1 | 29 | STAT3 |
| Down | SPINT2 | 29 | hsa-mir-4677-3p | Down | ERG | 28 | TFAP2C |
| Down | KRT18 | 27 | hsa-mir-126-3p | Down | SPINT2 | 26 | NACC1 |
| Down | EEF1A2 | 27 | hsa-mir-16-5p | Down | EEF1A2 | 24 | DNAJC2 |
| Down | STX11 | 21 | hsa-mir-373-3p | Down | CHRM4 | 22 | SOX9 |
| Down | PRKD1 | 19 | hsa-mir-34a-5p | Down | PRKD1 | 21 | MTF2 |
| Down | ACTBL2 | 14 | hsa-mir-599 | Down | ACTBL2 | 18 | YAP1 |
| Down | ERG | 9 | hsa-mir-145-5p | Down | MDFI | 17 | TET1 |
| Down | FOXS1 | 8 | hsa-mir-101-3p | Down | ACTC1 | 15 | TBX5 |
| Down | CHRM4 | 4 | hsa-mir-146a-5p | Down | FOXS1 | 11 | EP300 |

### Construction of the TF-hub gene regulatory network

We searched for target-regulated hub gene TFs using NetworkAnalyst database and then used the results of this database. By constructing TF-hub gene regulatory network networks, we found 520 nodes (TF: 198; Hub Gene: 322) and 8331 edges (Fig. 6). We identified 59 TFs (ex; TCF3) targeting regulation of PTCH1, 42 TFs (ex; PHC1) targeting regulation of CCND2, 37 TFs (ex; NR1I2) targeting regulation of PRKCB, 36 TFs (ex; HOXC9) targeting regulation of ST8SIA4, 34 TFs (ex; RNF2) targeting regulation of FOXQ1, 44 TFs (ex; CLOCK) targeting regulation of DAPK1, 38 TFs (ex; PRDM14) targeting regulation of IGF2BP1, 36 TFs (ex; SMARCA4) targeting regulation of KRT18, 34 TFs (ex; TRIM28) targeting regulation of STX11 and 31 TFs (ex; HTT) targeting regulation of EPB41L3, and are listed Table 5.

**Fig. 6.**
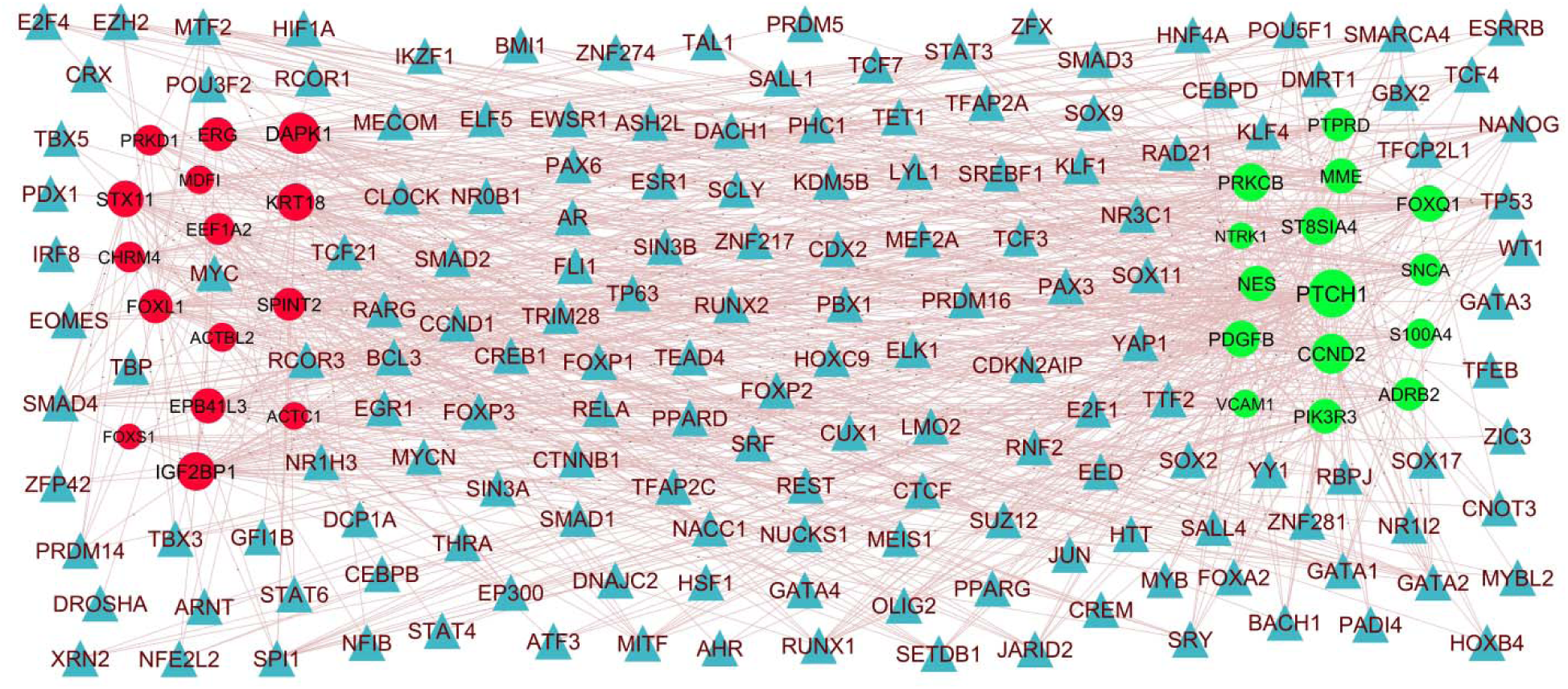
Hub gene - TF regulatory network. The blue color triangle nodes represent the key TFs; up regulated genes are marked in dark green; down regulated genes are marked in dark red.

### Receiver operating characteristic curve (ROC) analysis

The ROC curve was used to evaluate the diagnostic value of hub genes. As shown in Fig. 7, the AUC values of VCAM1, SNCA, PRKCB, ADRB2, FOXQ1, MDFI, ACTBL2, PRKD1, DAPK1 and ACTC1 in endometriosis were 0.904, 0.907, 0.903, 0.926, 0.901, 0.910, 0.923, 0.892, 0.895 and 0.898, respectively. Thus, the hub genes have good diagnostic efficiency in endometriosis and normal control samples.

**Fig. 7.**
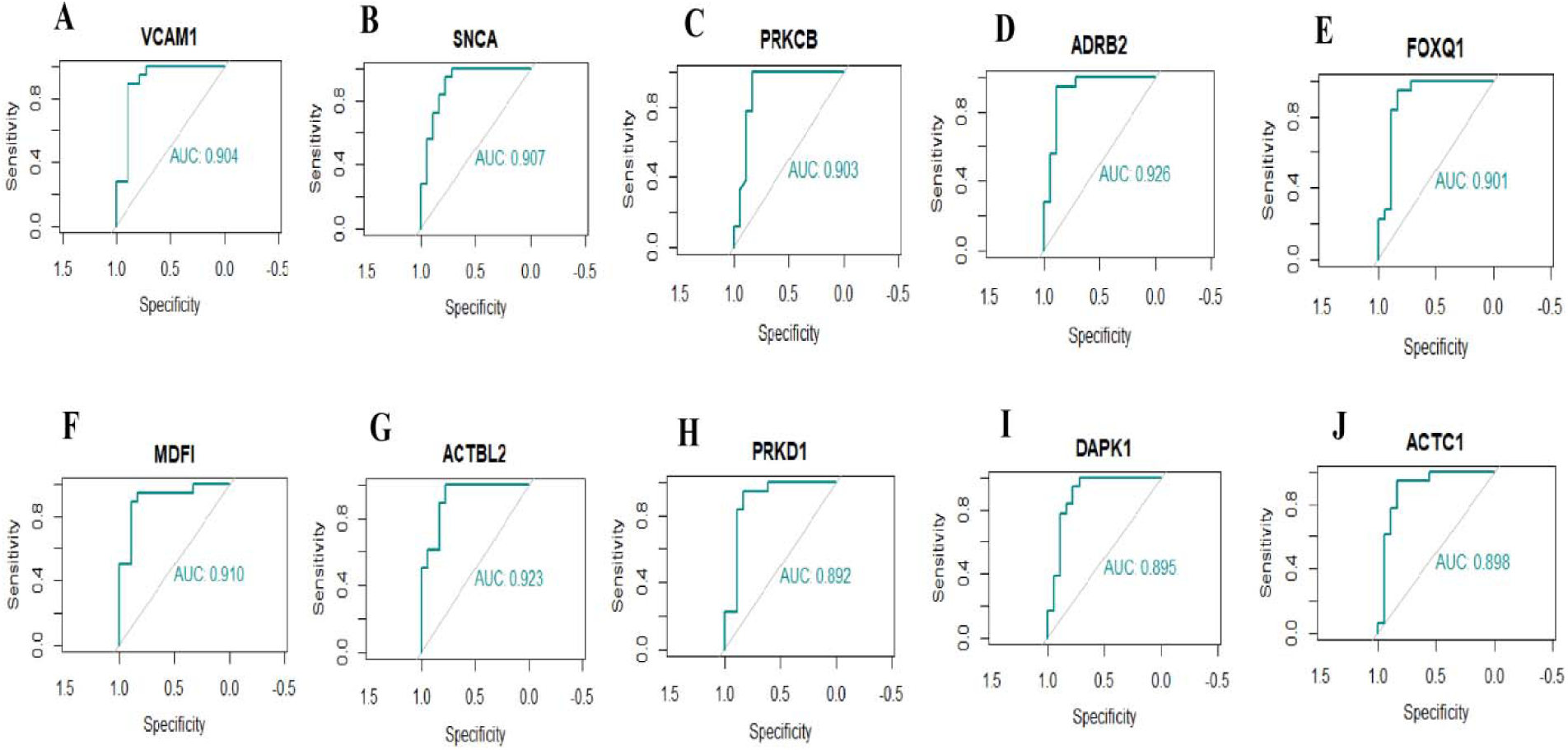
ROC curve analyses of hub genes. A) VCAM1 B) SNCA C) PRKCB D) ADRB2 E) FOXQ1 F) MDFI G) ACTBL2 H) PRKD1 I) DAPK1 J) ACTC1

## Discussion

Endometriosis is a major cause of serious reproductive disorder in the female population and leads to a great public health burden. Failure of early screening and diagnosis of endometriosis leads to progressive worsening of the dysmenorrhea, dyspareunia, chronic pelvic pain, irregular uterine bleeding and infertility. The degree of endometriosis seriously affects the recovery from female reproductive diseases. Therefore, the investigation of biomarkers of endometriosis is significant for the early diagnosis and treatment of the disease. With the advancement of bioinformatics and NGS technology, it has started to be widely applied to find the hub genes of diseases. The bioinformatics analysis of NGS data provides a convenient and comprehensive platform to reveal the molecular mechanism of endometriosis occurrence and progression, and find effective target drugs for the treatment of endometriosis.

In this investigation, we analyzed the endometriosis GSE243039 screened from the GEO database. It includes 20 normal samples and 20 endometriosis samples. Compared to normal controls, we found 958 DEGs (including 479 up regulated genes and 479 down regulated genes). PCSK9 [48], CNTN4 [49], SEMA3A [50], SFRP4 [51], MFAP5 [52], BMP6 [53], CDH6 [54], PIEZO2 [55] and PKP2 [56] are a diagnostic markers for inflammation. PCSK9 [57], SEMA3A [58] and SFRP4 [59] are associated with pain. PCSK9 [60], CNTN4 [61], SEMA3A [62], PTGIS (prostaglandin I2 synthase) [63], SFRP4 [64], MFAP5 [65], CDH6 [66], GPC6 [67] and PKP2 [68] might serve as genetic markers of ovarian cancer. The expression of PCSK9 [69], SFRP4 [70] and BMP6 [71] were altered in the polycystic ovarian syndrome. Altered expression of PCSK9 [72], CNTN4 [49], SEMA3A [73], SFRP4 [74], MFAP5 [75], BMP6 [76], PDE1C [77] and PKP2 [78] promotes the development of cardiovascular diseases. PCSK9 [79], APCDD1 [80], SFRP4 [81], MFAP5 [82] and PKP2 [83] are associated to the risk of obesity. Studies show that PCSK9 [84] and SFRP4 [85] are involved in the process of gestational diabetes mellitus. The PCSK9 [86], SEMA3A [87], SFRP4 [81] and BMP6 [88] were found as potential biomarkers for diabetes mellitus. PCSK9 [89], SEMA3A [90], PTGIS (prostaglandin I2 synthase) [91] and PIEZO2 [92] expression level is significantly altered in the hypertension. Studies have found that the ADAMTS19 [93] and BMP6 [94] plays a vital role in the development of infertility. Expression levels of SEMA3A [95], SFRP4 [96] and BMP6 [94] have been proved to be altered in endometriosis. PTGIS (prostaglandin I2 synthase) [97] and SFRP4 [98] are associated with prognosis in endometrial cancer. Altered levels of SFRP4 [99] and MFAP5 [100] have been shown to be associated with cervical cancer. This investigation might provide reference for research the connection between endometriosis and its associated complications.

In this investigation, we identified enriched genes in GO terms and signaling pathways that might be utilized as diagnostic and/or therapeutic targets in endometriosis. Signaling pathways include extracellular matrix organization [101], nervous system development [102], signal transduction [103], hemostasis [104], muscle contraction [105], signaling by retinoic acid [106] and diseases of glycosylation [107] were responsible for advancement of endometriosis. Altered expression of L1CAM [108], HSD17B2 [109], VCAM1 [110], SOX6 [111], FGF10 [112], MMP12 [113], CCR1 [114], PROK1 [115], PRL (prolactin) [116], TIMP3 [117], ADAMTS9 [118], NDNF (neuron derived neurotrophic factor) [119], LHCGR (luteinizing hormone/choriogonadotropin receptor) [120], PDGFB (platelet derived growth factor subunit B) [121], LDLR (low density lipoprotein receptor) [122], CD4 [123], FOXL2 [124], TRPA1 [125], ADRB2 [126], PLAU (plasminogen activator, urokinase) [127], EPCAM (epithelial cell adhesion molecule) [128], UCN2 [129], CYP1A1 [130], NTN1 [131], IL15 [132], BMP2 [133], APOE (apolipoprotein E) [134], CASP1 [135], ABCG2 [136], ACE (angiotensin I converting enzyme) [137], PGR (progesterone receptor) [138], ALPP (alkaline phosphatase, placental) [139], LPAR4 [140], ATRNL1 [141], HLA-C [142], MMP3 [143], PDLIM3 [144], NFASC (neurofascin) [145], IL33 [146], NGF (nerve growth factor) [147], COMP (cartilage oligomeric matrix protein) [148], FST (follistatin) [149], EFEMP1 [150], GATA6 [151], TCF21 [152], PTGS2 [153], HOXC8 [154], AKR1C3 [155], BDNF (brain derived neurotrophic factor) [119], EPHA3 [156], INHBA (inhibin subunit beta A) [157], RAP1GAP [158], TLR3 [159], NOX4 [160], TGFBI (transforming growth factor beta induced) [161], IGF2BP1 [162], DLX5 [163], VDR (vitamin D receptor) [164], FZD7 [165], ID2 [166], TLR2 [167], IL6 [168], GAS6 [169], DUSP2 [170], FGF7 [171], CCN2 [172], IGFBP3 [173], CHL1 [174], BGN (biglycan) [175], NTRK2 [176], SLIT2 [177], NOTCH2 [178], LIF (LIF interleukin 6 family cytokine) [179], CD200 [180], BST2 [181], DYSF (dysferlin) [182], DAPK1 [183], KISS1 [184], FPR1 [185] and TRH (thyrotropin releasing hormone) [186] promotes endometriosis. Transcription of L1CAM [187], AJAP1 [188], HSD17B2 [189], VCAM1 [190], GRP (gastrin releasing peptide) [191], AQP8 [192], WNT6 [193], FABP4 [194], SOX6 [195], NTRK1 [196], CNTN1 [197], MMP12 [198], LAG3 [199], SOX18 [200], CCR1 [201], FLT1 [202], PRDM1 [203], TRPC3 [204], DKK2 [205], RNF157 [206], DHCR24 [207], BMP4 [208], PRL (prolactin) [209], FOXQ1 [210], WNT5A [211], MEOX1 [212], DOCK4 [213], TIMP3 [214], ADAMTS9 [215], NDNF (neuron derived neurotrophic factor) [216], NETO1 [217], CD24 [218], LHCGR (luteinizing hormone/choriogonadotropin receptor) [219], SCD (stearoyl-CoA desaturase) [220], PDGFB (platelet derived growth factor subunit B) [221], MMRN1 [222], LDLR (low density lipoprotein receptor) [223], CD4 [224], FOXL2 [225], TRPA1 [226], EPHA5 [227], TOX (thymocyte selection associated high mobility group box) [228], CST4 [229], RSPO3 [230], MAP2K6 [231], NES (nestin) [232], TMEM119 [233], PADI2 [234], MMP8 [235], KDR (kinase insert domain receptor) [236], ADRB2 [237], MGAT3 [238], PTPRC (protein tyrosine phosphatase receptor type C) [239], PITX1 [240], KL (klotho) [241], PLAU (plasminogen activator, urokinase) [242], ZNF365 [243], PIK3R3 [244], SOX8 [245], CCND2 [246], CRABP2 [247], PCDH9 [248], EPCAM (epithelial cell adhesion molecule) [249], CLEC14A [250], CYP1A1 [251], NTN1 [252], PDGFD (platelet derived growth factor D) [253], CLDN3 [254], LEPR (leptin receptor) [255], IL15 [256], BMP2 [257], LAMA5 [258], NTNG1 [259], KRT19 [260], ROS1 [261], APOE (apolipoprotein E) [262], PTCH1 [263], ITPKA (inositol-trisphosphate 3-kinase A) [264], CASP1 [265], NID1 [266], ABCG2 [267], ACE (angiotensin I converting enzyme) [268], PGR (progesterone receptor) [269], WLS (Wnt ligand secretion mediator) [270], KLK3 [271], LRP1B [272], LY6K [273], ALPP (alkaline phosphatase, placental) [274], PRAME (PRAME nuclear receptor transcriptional regulator) [275], SLCO4A1 [276], EGFL6 [277], GPBAR1 [278], ELMO1 [279], WNK2 [280], IL2RB [281], DIRAS2 [282], GALNT14 [283], RTKN2 [284], ATRNL1 [285], S100A4 [286], MACC1 [287], MMP3 [288], COL11A1 [289], CABLES1 [290], FGFR2 [291], IL33 [292], NGF (nerve growth factor) [293], FOXC2 [294], COMP (cartilage oligomeric matrix protein) [295], FST (follistatin) [296], SORBS2 [297], EFEMP1 [298], GATA6 [299], TCF21 [300], PTGS2 [301], MTSS1 [302], DACT1 [303], HOXC8 [304], PITX2 [305], TNFSF10 [306], BDNF (brain derived neurotrophic factor) [307] KRT7 [308], NDRG2 [309], EYA2 [310], INHBA (inhibin subunit beta A) [311], SGK1 [312], SLC2A12 [313], DIO3 [314], EPB41L3 [315], TLR3 [316], ANGPTL4 [317], EPHB2 [318], FLI1 [319], THBS1 [320], ID3 [321], NOX4 [322], TGFBI (transforming growth factor beta induced) [323], IGF2BP1 [324], SALL4 [325], DLX5 [326], VDR (vitamin D receptor) [327], LZTS1 [328], FZD7 [329], EN2 [330], ENC1 [331], IFNE (interferon epsilon) [332], TNNT1 [333], ANKRD1 [334], SOX9 [335], MGP (matrix Gla protein) [336], SULF1 [337], CYP24A1 [338], DNAH11 [339], TLR2 [340], IL6 [341], NPPB (natriuretic peptide B) [342], SPINK1 [343], GPC3 [344], NTRK3 [345], AMIGO2 [346], FOXD1 [347], ADAM12 [348], DUSP2 [349], USP2 [350], KLF2 [351], SIK1 [352], SIX1 [310], FGF7 [353], MYH10 [354], IGFBP3 [355], LYVE1 [356], ACTBL2 [357], SLIT2 [358], ACTC1 [359], NNMT (nicotinamide N-methyltransferase) [360], CHI3L1 [361], RUNX1 [362], NFIB (nuclear factor I B) [363], NOTCH2 [364], PGF (placental growth factor) [365], THBS2 [366], NAV1 [367], NRG1 [368], PLK2 [369], ITGBL1 [370], CD200 [371], BST2 [372], KCNN3 [373], HMCN1 [374], VEPH1 [375], TFPI2 [376], SYTL2 [377], CCDC80 [378], DAPK1 [379], KISS1 [380], IL20RA [381], HAS3 [382], HAS1 [382], MGST1 [383], FPR1 [384] and SH3RF2 [385] were significantly altered in patients with ovarian cancer. A previous study reported that the L1CAM [386], HSD17B2 [387], GRP (gastrin releasing peptide) [388], FABP4 [389], SOX6 [390]. MMP12 [391], APOD (apolipoprotein D) [392], LAG3 [393], CST1 [394], FLT1 [395], DHCR24 [396], PRL (prolactin) [397], WNT5A [398], TIMP3 [399], CD24 [400], LHCGR (luteinizing hormone/choriogonadotropin receptor) [401], MMRN1 [402], CD4 [403], ADAMTS5 [404], ADAMTS1 [405], PADI2 [406], MARK1 [407], KL (klotho) [408], PLAU (plasminogen activator, urokinase) [409], SOX8 [410], CRABP2 [411], PTPRD (protein tyrosine phosphatase receptor type D) [412], EPCAM (epithelial cell adhesion molecule) [413], IRX2 [414], SEMA3B [415], CYP1A1 [416], PDGFD (platelet derived growth factor D) [417], LEPR (leptin receptor) [418], APOE (apolipoprotein E) [419], CASP1 [420], MGLL (monoglyceride lipase) [421], NID1 [422], ABCG2 [423], ACE (angiotensin I converting enzyme) [424], PGR (progesterone receptor) [425], HPSE2 [426], LMTK3 [427], ALPP (alkaline phosphatase, placental) [428], EGFL6 [429], CACNA2D3 [430], MCTP1 [431], HKDC1 [432], S100A4 [433], MACC1 [434], MMP3 [435], FGFR2 [436], IL33 [437], FOXC2 [438], ITGA7 [439], EFEMP1 [440], GATA6 [441], BHLHE41 [442], TCF21 [443], GDF10 [444], NKX3-1 [445], AKR1C3 [446], SGK1 [447], RAP1GAP [448], FLI1 [449], NOX4 [450], SERPINE2 [451], IGSF9 [452], IGF2BP1 [453], SALL4 [454], VDR (vitamin D receptor) [455], CELSR2 [456], ENC1 [457], SOX9 [458], CYP24A1 [338], IL6 [459], GAS6 [460], KLF2 [461], SIX1 [462], IGFBP3 [463], LYVE1 [464], CHL1 [465], BGN (biglycan) [466], SLIT2 [467], NRP2 [468], NNMT (nicotinamide N-methyltransferase) [469], RUNX1 [470], THBS2 [471], HSPB7 [472], NRG1 [473], TFPI2 [474], DAPK1 [183], HAS3 [475], HAS1 [475], STEAP1 [476] and MGST1 [477] genes were associated with the endometrial cancer. VCAM1 [478], AQP8 [479], L1CAM [480], FABP4 [481], PSG1 [482], SOX6 [483], MMP12 [484], APOD (apolipoprotein D) [485], LAG3 [486], SOX18 [487], FLT1 [488], FABP5 [489], BMP4 [490], PRL (prolactin) [491], FOXQ1 [492], WNT5A [493], FRZB (frizzled related protein) [494], CPE (carboxypeptidase E) [495], EREG (epiregulin) [496], NDNF (neuron derived neurotrophic factor) [497], CD24 [498], SCD (stearoyl-CoA desaturase) [499], LDLR (low density lipoprotein receptor) [500], CD4 [501], FOXL2 [502], KRT17 [503], NES (nestin) [504], MGAT3 [505], MARK1 [506], KL (klotho) [507], PLAU (plasminogen activator, urokinase) [508], EPHA7 [509], PIK3R3 [510], CCND2 [511], HECW1 [512], EPCAM (epithelial cell adhesion molecule) [513], BATF2 [514], CYP1A1 [515], MSTN (myostatin) [516], IL15 [517], SYT7 [518], PAK3 [519], KRT19 [520], ROS1 [521], CUBN (cubilin) [522], PTCH1 [523], CASP1 [524], ABCG2 [525], PGR (progesterone receptor) [526], HPSE2 [527], LRP1B [528], ALPP (alkaline phosphatase, placental) [529], CYP2S1 [530], DOC2B [531], MSMO1 [532], SORCS1 [533], HLA-C [534], S100A4 [535], MACC1 [536], MMP3 [537], FGFR2 [538], IL33 [539], NGF (nerve growth factor) [540], FOXC2 [541], SORBS2 [542], ITGA7 [543], EFEMP1 [544], GATA6 [545], TCF21 [546], PTGS2 [547], MTSS1 [548], DACT1 [549], SPINT2 [550], NKX3-1 [551], HOXC8 [552], AKR1C3 [Wu et al. 2014], BDNF (brain derived neurotrophic factor) [497], NDRG2 [553], EPHA3 [554], EYA2 [555], INHBA (inhibin subunit beta A) [556], ALPL (alkaline phosphatase, biomineralization associated) [557], SGK1 [558], RAP1GAP [559], EPB41L3 [560], TLR3 [561], ANGPTL4 [562], EPHB2 [563], FLI1 [564], THBS1 [565], NOX4 [566], TGFBI (transforming growth factor beta induced) [567], IGF2BP1 [568], SALL4 [569], VDR (vitamin D receptor) [570], RARB (retinoic acid receptor beta) [571], EPHA4 [572], ENC1 [573], SOX9 [574], SULF1 [575], TLR2 [576], IL6 [577], GPC3 [578], NTRK3 [579], CCNA1 [580], AMIGO2 [581], FOXD1 [582], CCNO (cyclin O) [583], ADAM12 [584], RASSF2 [585], HOXB7 [586], KLF2 [587], SIK1 [588], SIX1 [589], FGF7 [590], IGFBP3 [591], CHL1 [592], EPPK1 [593], SLIT2 [594], FLG (filaggrin) [595], NRP2 [596], NNMT (nicotinamide N-methyltransferase) [597], CHI3L1 [598], RUNX1 [599], APLN (apelin) [600], SEMA3C [601], NOTCH2 [602], THBS2 [603], PNPLA1 [604], BST2 [605], HMCN1 [606], ULBP1 [607], TFPI2 [608], DAPK1 [609], KISS1 [610], FPR1 [611] and PIK3AP1 [612] have been used as an independent biomarkers to predict prognosis in patients with cervical cancer. CBLN2 [613], SDK1 [614], VCAM1 [615], SIX2 [616], AVPR1A [617], EPHA6 [618], FABP4 [619], PSG1 [620], ANO1 [621], SOX6 [622], FGF10 [623], PLA2G7 [624], MMP12 [625], ADRA1D [626], LAG3 [627], FLT1 [628], FABP5 [629], PRDM1 [630], TRPC3 [631], IGSF3 [632], BMP4 [633], IL1RL1 [634], PRL (prolactin) [635], NEFL (neurofilament light chain) [636], WNT5A [637], TIMP3 [638], NDNF (neuron derived neurotrophic factor) [639], SNAP25 [640], CD24 [641], PDGFB (platelet derived growth factor subunit B) [642], LDLR (low density lipoprotein receptor) [643], CD4 [644], TRPA1 [645], ADAMTS1 [646], PDE4B [647], NES (nestin) [648], TH (tyrosine hydroxylase) [649], PSG9 [650], CACNA1D [651], MMP8 [652], ADRB2 [653], KL (klotho) [654], PLAU (plasminogen activator, urokinase) [655], PTPRD (protein tyrosine phosphatase receptor type D) [656], SEMA3B [657], UCN2 [658], CYP2J2 [659], CYP1A1 [660], ATP1A2 [661], CLDN3 [662], MSTN (myostatin) [663], LEPR (leptin receptor) [664], IL15 [665], CACNA1H [666], BMP2 [667], LAMA5 [668], ROS1 [669], APOE (apolipoprotein E) [670], CASP1 [671], PDE9A [672], EFNB2 [673], ABCG2 [674], ACE (angiotensin I converting enzyme) [675], PGR (progesterone receptor) [676], SLC35F3 [677], ICA1 [678], ALPP (alkaline phosphatase, placental) [679], TRPC6 [680], GPBAR1 [681], PNPLA3 [682], HLA-C [683], S100A4 [684], MACC1 [685], MMP3 [686], GDNF (glial cell derived neurotrophic factor) [687], FGFR2 [688], IL33 [689], NGF (nerve growth factor) [690], PAPPA2 [691], COMP (cartilage oligomeric matrix protein) [692], GATA6 [693], ACAN (aggrecan) [694], TCF21 [695], PTGS2 [696], PITX2 [697], AKR1C3 [698], BDNF (brain derived neurotrophic factor) [699], SGK1 [700], TLR3 [701], ANGPTL4 [702], FLI1 [703], THBS1 [704], ID3 [705], NOX4 [706], PCSK1 [707], ITGB1BP2 [708], WNK4 [709], DLX5 [710], VDR (vitamin D receptor) [711], EPHA4 [712], MGP (matrix Gla protein) [713], CYP24A1 [714], ID2 [715], TLR2 [716], IL6 [717], NPPB (natriuretic peptide B) [718], GAS6 [719], F11R [720], FOXD1 [721], ADAM12 [722], NCAM1 [723], USP2 [724], KLF2 [725], SIK1 [726], FGF7 [727], IGFBP3 [728], BGN (biglycan) [729], NTRK2 [730], NNMT (nicotinamide N-methyltransferase) [731], CHI3L1 [732], RUNX1 [733], APLN (apelin) [734], STOX2 [735], KCNQ4 [736], NOTCH2 [737], PGF (placental growth factor) [738], THBS2 [739], PDLIM5 [740], PRDM6 [741], HTR6 [742], NRG1 [743], CD200 [744], BST2 [745], KCNN3 [746], SLC2A5 [747], TFPI2 [748], DYSF (dysferlin) [749], CCDC80 [750], DAPK1 [751], KISS1 [752], SLC4A4 [753], STEAP2 [754], SORBS1 [755], ACKR2 [756], FPR1 [757], GPR143 [758] and TRH (thyrotropin releasing hormone) [759] can be used as a diagnostic markers for hypertension. ROBO2 [760], VCAM1 [761], GRP (gastrin releasing peptide) [762], FABP4 [763], ANO1 [764], SOX6 [765], TFAP2C [766], RAMP3 [767], PLA2G7 [768], MMP12 [769], FAIM2 [770], APOD (apolipoprotein D) [771], LAG3 [772], SOX18 [773], F2RL2 [774], CCR1 [775], FLT1 [776], FABP5 [629], TRPC3 [777]. THSD7A [778], DKK2 [779], PRKCB (protein kinase C beta) [780], DHCR24 [781], PDE3B [782], BMP4 [783], IL1RL1 [784], MYPN (myopalladin) [785], PLCG2 [786], PRL (prolactin) [787], WNT5A [788], MEOX1 [789], TIMP3 [790], FRZB (frizzled related protein) [791], CPE (carboxypeptidase E) [792], ADAMTS9 [793], NDNF (neuron derived neurotrophic factor) [794], PDGFB (platelet derived growth factor subunit B) [795], PIK3CG [796], LDLR (low density lipoprotein receptor) [797], CD4 [798], TRPA1 [799], F2RL3 [800], C1QL1 [801], ADAMTS5 [802], PDE4B [803], NES (nestin) [804], TH (tyrosine hydroxylase) [805], MMP8 [806], KDR (kinase insert domain receptor) [807], ADRB2 [808], ACKR3 [809], PTPRC (protein tyrosine phosphatase receptor type C) [810], KL (klotho) [811], KL (klotho) [812], PLAU (plasminogen activator, urokinase) [813], CCND2 [814], PTGS1 [815], INSIG1 [816], IRX2 [817], SIGLEC1 [818], UCN2 [819], CYP2J2 [820], CYP1A1 [821], ASTN2 [822], NTN1 [823], PDGFD (platelet derived growth factor D) [824], MSTN (myostatin) [663], LEPR (leptin receptor) [664], IL15 [825], CACNA1H [826], BMP2 [827], SYT7 [828], ZBTB46 [829], ROS1 [830], APOE (apolipoprotein E) [831], CUBN (cubilin) [832], RBM20 [833], CASP1 [834], PDE9A [835], ABCG2 [836], HMGCR (3-hydroxy-3- methylglutaryl-CoA reductase) [837], ACE (angiotensin I converting enzyme) [838], GREM2 [839], PALMD (palmdelphin) [840], LRP1B [841], ALPP (alkaline phosphatase, placental) [842], TRPC6 [843], GPBAR1 [844], MYZAP (myocardial zonulaadherens protein) [845], PRODH (proline dehydrogenase 1) [846], IL2RB [847], CDHR3 [848], PNPLA3 [849], FADS1 [850], HLA-C [851], S100A4 [852], MMP3 [853], PDLIM3 [854], GDNF (glial cell derived neurotrophic factor) [855], FGFR2 [856], IL33 [857], NGF (nerve growth factor) [858], HAPLN1 [859], FOXC2 [860], COMP (cartilage oligomeric matrix protein) [861], FST (follistatin) [862], SORBS2 [863], ITGA7 [864], PLN (phospholamban) [865], GATA6 [866], BHLHE41 [867], ACAN (aggrecan) [868], TCF21 [869], PTGS2 [870], DACT1 [871], PITX2 [872], AKR1C3 [873], BDNF (brain derived neurotrophic factor) [874], NDRG2 [875], EYA2 [876], SGK1 [877], RAP1GAP [878], DIO3 [879], TLR3 [880], ANGPTL4 [881], EPHB2 [882], THBS1 [883], TNNT2 [884], NOX4 [885], S1PR5 [886], SERPINE2 [887], PCSK1 [888], TGFBI (transforming growth factor beta induced) [889], SALL4 [890], EYA4 [891], ITGB1BP2 [708], VDR (vitamin D receptor) [892], GPC4 [893], CELSR2 [894], EPHA4 [895], TNNT1 [896], ANKRD1 [897], ZFPM2 [898], SOX9 [899], MGP (matrix Gla protein) [900], CYP24A1 [901], DNAH11 [902], TLR2 [903], IL6 [904], GAS6 [905], GPC3 [906], NTRK3 [907], AMIGO2 [908], F11R [909], ADAM12 [722], NCAM1 [910], USP2 [911], KLF2 [912], SIK1 [913], SIX1 [914], FGF7 [915], CCN2 [916], JCAD (junctional cadherin 5 associated) [917], IGFBP3 [918], LYVE1 [919], PRKD1 [920], BGN (biglycan) [921], EDA (ectodysplasin A) [922], SLIT2 [923], ACTC1 [924], NRP2 [925], CHI3L1 [926], RUNX1 [927], APLN (apelin) [928], MYOM2 [929], MYOZ1 [930], PPP1R13L [931], THBS2 [932], DES (desmin) [933], PDLIM5 [934], HSPB7 [935], NRG1 [936], PLK2 [937], ITGBL1 [938], CD200 [939], KCNN3 [940], KCNJ2 [941], EVA1A [942], TFPI2 [943], DYSF (dysferlin) [944], ADAP1 [945], CCDC80 [946], DAPK1 [947], SCN4B [948], ESYT3 [949], ABCA8 [950], HEG1 [951], FPR1 [952], SSPN (sarcospan) [953], ADH1C [954], SIRPA (signal regulatory protein alpha) [955] and TRH (thyrotropin releasing hormone) [956] might be a prognostic biomarkers and potential therapeutic targets for patients with cardiovascular diseases. HSD17B2 [109], EFNA5 [957], MMP12 [958], PROK1 [959], PRL (prolactin) [960], NLRP2 [961], NDNF (neuron derived neurotrophic factor) [962], MEI4 [963], CD24 [964], LHCGR (luteinizing hormone/choriogonadotropin receptor) [965], CD4 [966], FOXL2 [967], KDR (kinase insert domain receptor) [968], ADRB2 [969], CYP1A1 [970], NTN1 [971], MSTN (myostatin) [972], BMP2 [973], APOE (apolipoprotein E) [974], ACE (angiotensin I converting enzyme) [975], PGR (progesterone receptor) [976], GREM2 [977], ALPP (alkaline phosphatase, placental) [978], MMP3 [979], GDNF (glial cell derived neurotrophic factor) [980], FGFR2 [981], IL33 [982], NGF (nerve growth factor) [983], COMP (cartilage oligomeric matrix protein) [984], CECR2 [985], FST (follistatin) [986], GATA6 [987], PTGS2 [153], BDNF (brain derived neurotrophic factor) [988], SGK1 [989], ANGPTL4 [990], THBS1 [991], ID3 [992], NOX4 [993], IGF2BP1 [994], SALL4 [995], VDR (vitamin D receptor) [996], SULF1 [997], TLR2 [998], IL6 [999], GPC3 [1000], CCNO (cyclin O) [1001], IGFBP3 [1002], CHL1 [1003], NTRK2 [1004], SLIT2 [1005], APLN (apelin) [1006], NOTCH2 [1007], PGF (placental growth factor) [1008], LIF (LIF interleukin 6 family cytokine) [1009], CD200 [1010], TFPI2 [1011], KISS1 [752] and TRH (thyrotropin releasing hormone) [1012] were associated with a favorable prognosis for infertility. A altered expression level of VCAM1 [761], STRA6 [1013], COCH (cochlin) [1014], GRP (gastrin releasing peptide) [1015], AQP8 [1016], FABP4 [1017], ANO1 [1018], SOX6 [1019], TFAP2C [1020], NTRK1 [1021], CNTN1 [1022], FGF10 [1023], PLA2G7 [768], MMP12 [1024], LCP1 [1025], SNCA (synuclein alpha) [1026], APOD (apolipoprotein D) [1027], LAG3 [1028], CCR1 [1029], CST1 [1030], RETREG1 [1031], FLT1 [1032], FABP5 [1033], PRDM1 [1034], TRPC3 [1035], PROK1 [1036], WNT16 [1037], F13A1 [1038], DHCR24 [1039], PDE3B [1040], BMP4 [783], IL1RL1 [1041], PLCG2 [1042], PRL (prolactin) [1043], FOXQ1 [1044], NEFL (neurofilament light chain) [1045], WNT5A [1046], TIMP3 [1047], SERPINB2 [1048], FRZB (frizzled related protein) [1049], NLRP2 [1050], CPE (carboxypeptidase E) [1051], ADAMTS9 [1052], NPW (neuropeptide W) [1053], EREG (epiregulin) [1054], NDNF (neuron derived neurotrophic factor) [1055], SNAP25 [1056], SYT1 [1057], SCD (stearoyl-CoA desaturase) [1058], PDGFB (platelet derived growth factor subunit B) [1059], LDLR (low density lipoprotein receptor) [1060], CD4 [1061], GPR183 [1062], TRPA1 [125], PTGER4 [1063], ADAMTS5 [1064], RSPO3 [1065], KRT17 [1066], ADAMTS1 [1067], PDE4B [1068], NES (nestin) [804], SH2D2A [1069], TH (tyrosine hydroxylase) [1070], TMEM119 [1071], MMP8 [1072], ADRB2 [1073], ACKR3 [1074], MGAT3 [1075], TNFRSF9 [1076], TXK (TXK tyrosine kinase) [1077], KL (klotho) [811], PLAU (plasminogen activator, urokinase) [1078], EPHA7 [1079], ZNF365 [1080], PIK3R3 [1081], SOX8 [1082], CCND2 [1083], PTGS1 [1084], BATF2 [1085], UCN2 [1086], CYP2J2 [1087], CLEC14A [1088], CYP1A1 [1089], NTN1 [1090], MSTN (myostatin) [1091], LEPR (leptin receptor) [1092], CD248 [1093], IL15 [825], BMP2 [1094], ROS1 [1095], MME (membrane metalloendopeptidase) [1096], APOE (apolipoprotein E) [1097], PTCH1 [1098], CASP1 [1099], MGLL (monoglyceride lipase) [1100], EFNB2 [1101], ABCG2 [1102], HMGCR (3-hydroxy-3-methylglutaryl-CoA reductase) [1103], ACE (angiotensin I converting enzyme) [1104], PGR (progesterone receptor) [1105], GREM2 [839], RGS7 [1106], CHST1 [1107], ALPP (alkaline phosphatase, placental) [1108], TRPC6 [1109], SLCO4A1 [276], CYP4B1 [1110], GPBAR1 [844], ELMO1 [1111], DOC2B [1112], CD163L1 [1113], SLCO2A1 [1114], IL2RB [1115], B4GALNT2 [1116], SLC37A2 [1117], PNPLA3 [1118], FADS1 [1119], HLA-C [1120], ST3GAL5 [1121], S100A4 [1122], MACC1 [1123], CORT (cortistatin) [1124], MMP3 [1125], GDNF (glial cell derived neurotrophic factor) [1126], LMO3 [1127], NFASC (neurofascin) [1128], FGFR2 [1129], IL33 [1130], NGF (nerve growth factor) [1131], HAPLN1 [1132], GDF6 [1133], FOXC2 [1134], COMP (cartilage oligomeric matrix protein) [1135], FST (follistatin) [1136], ITGA7 [1137], GATA6 [1138], ACAN (aggrecan) [1139], TCF21 [1140], PTGS2 [1141], MTSS1 [1142], DHRS3 [1143], NKX3-1 [1144], TNFSF10 [1145], BDNF (brain derived neurotrophic factor) [1146], NDRG2 [1147], EPHA3 [1148], PLA2G5 [1149], MECOM (MDS1 and EVI1 complex locus) [1150], SGK1 [1151], TLR3 [1152], ANGPTL4 [1153], EPHB2 [1154], FLI1 [1155], THBS1 [1156], MBP (myelin basic protein) [1157], ID3 [1158], NOX4 [1159], S1PR5 [1160], PI16 [1161], IGF2BP1 [1162], SALL4 [1163], VLDLR (very low density lipoprotein receptor) [1164], VDR (vitamin D receptor) [1165], FAM20A [1166], EPHA4 [1167], ANKRD1 [1168], SGCA (sarcoglycan alpha) [1169], SOX9 [1170], MGP (matrix Gla protein) [1171], CYP24A1 [1172], TLR2 [1173], IL6 [1174], GAS6 [1175], NTRK3 [1176], ADAM12 [1177], NCAM1 [1178], MYOC (myocilin) [1179], USP2 [1180], KLF2 [1181], SIK1 [1182], SIX1 [1183], FGF7 [1184], CCN2 [1185], IGFBP3 [1186], LYVE1 [1187], PRKD1 [1188], BGN (biglycan) [1189], SLIT2 [1190], IRX3 [1191], ACTC1 [1192], FLG (filaggrin) [1193], NRP2 [1194], NNMT (nicotinamide N-methyltransferase) [1195], CHI3L1 [1196], RUNX1 [1197], NFIB (nuclear factor I B) [1198], APLN (apelin) [1199], PLP1 [1200], NAV2 [1201], NOTCH2 [1202], PGF (placental growth factor) [1203], THBS2 [1204], NRG1 [1205], LIF (LIF interleukin 6 family cytokine) [1206], PLK2 [1207], NALCN (sodium leak channel, non-selective) [1208], CD200 [1209], KCNN3 [1210], EVA1A [1211], TFPI2 [1212], DYSF (dysferlin) [1213], SYTL2 [1214], TLR1 [1215], CCDC80 [946], DAPK1 [947], KISS1 [1216], GEM (GTP binding protein overexpressed in skeletal muscle) [1217], IL20RA [1218], HAS3 [1219], HAS1 [1220], SLC4A4 [1221], SIRPB1 [1222], STEAP1 [1223], ACKR2 [1224], FPR1 [1225], GNG7 [1226], IGFBPL1 [1227], PIK3AP1 [1228], ADH1C [1229], LXN (latexin) [1230] and TRH (thyrotropin releasing hormone) [1231] have been detected in the inflammation. VCAM1 [1232], AQP8 [1233], FABP4 [1234], FABP5 [1235], BMP4 [1236], PRL (prolactin) [960], WNT5A [1237], ADAMTS9 [1238], NDNF (neuron derived neurotrophic factor) [1239], LHCGR (luteinizing hormone/choriogonadotropin receptor) [1240], LDLR (low density lipoprotein receptor) [1241], CD4 [1242], ADAMTS5 [1243], MAP2K6 [1244], ADAMTS1 [1245], PDE4B [1246], TH (tyrosine hydroxylase) [1247], MMP8 [1248], ADRB2 [969], KL (klotho) [1249], EPHA7 [1250], UCN2 [1251], CYP1A1 [970], LEPR (leptin receptor) [1252], IL15 [1253], BMP2 [1254], APOE (apolipoprotein E) [1255], CASP1 [1256], ACE (angiotensin I converting enzyme) [1257], PGR (progesterone receptor) [1258], GREM2 [1259], SORCS1 [1260], HKDC1 [1261], FADS1 [1262], S100A4 [1263], IL33 [1243], NGF (nerve growth factor) [1264], COMP (cartilage oligomeric matrix protein) [1265], FST (follistatin) [1266], GATA6 [1267], ACAN (aggrecan) [1268], AKR1C3 [1269], BDNF (brain derived neurotrophic factor) [1270], ANGPTL4 [1271], NOX4 [1272], VDR (vitamin D receptor) [1273], GPC4 [1274], TLR2 [1275], IL6 [1276], IGFBP3 [1277], TNIK (TRAF2 and NCK interacting kinase) [1278], APLN (apelin) [1279], PGF (placental growth factor) [1280], NRG1 [1281], LIF (LIF interleukin 6 family cytokine) [1282], ANGPTL1 [1283], KISS1 [1284], SORBS1 [1285] and TRH (thyrotropin releasing hormone) [1286] are a potential targets for polycystic ovarian syndrome. VCAM1 [1287], STRA6 [1288], AQP8 [1289], FABP4 [1290], SOX6 [1291], RAMP3 [1292], MMP12 [1293], FAIM2 [770], CCR1 [1294], ISM1 [1295], FLT1 [1296], FABP5 [1297], THSD7A [1298], SCTR (secretin receptor) [1299], WNT16 [1300], PRKCB (protein kinase C beta) [1301], PDE3B [1302], IL1RL1 [1041], PRL (prolactin) [1303], WNT5A [1304], HTR1B [1305], TIMP3 [1306], CPE (carboxypeptidase E) [1051], EREG (epiregulin) [1307], NDNF (neuron derived neurotrophic factor) [1308], SNAP25 [1309], CD24 [1310], SCD (stearoyl-CoA desaturase) [1058], PDGFB (platelet derived growth factor subunit B) [1311], LDLR (low density lipoprotein receptor) [1312], CD4 [1313], TRPA1 [1314], MAP2K6 [1315], PDE4B [1316], TH (tyrosine hydroxylase) [1317], MMP8 [1318], ADRB2 [1319], KL (klotho) [1320], PLAU (plasminogen activator, urokinase) [1321], PTGS1 [1084], INSIG1 [1322], BMP8A [1323], UCN2 [1324], NTN1 [1090], PDGFD (platelet derived growth factor D) [1325], MSTN (myostatin) [1326], LEPR (leptin receptor) [1092], IL15 [1327], BMP2 [1328], APOE (apolipoprotein E) [1329], CASP1 [1330], MGLL (monoglyceride lipase) [1331], NID1 [1332], ABCG2 [1333], ACE (angiotensin I converting enzyme) [1334], PGR (progesterone receptor) [1335], GREM2 [1336], LRP1B [1337], ALPP (alkaline phosphatase, placental) [1338], TRPC6 [1339], EGFL6 [1340], GPBAR1 [1341], AIF1L [1342], GPAT3 [1343], SORCS1 [1260], SLC37A2 [1344], FADS1 [1345], ACSL5 [1346], PTPRN2 [1347], S100A4 [1348], MACC1 [1349], CORT (cortistatin) [1350], MMP3 [1351], GDNF (glial cell derived neurotrophic factor) [1352], LMO3 [1353], CABLES1 [1354], IL33 [1355], NGF (nerve growth factor) [1356], FOXC2 [1357], FST (follistatin) [1136], PLN (phospholamban) [1358], ACAN (aggrecan) [1359], PTGS2 [1360], GDF10 [1361], CPNE5 [1362], DGAT2 [1363], BDNF (brain derived neurotrophic factor) [1364], RGS4 [1365], EPHA3 [1366], PLA2G5 [1367], SGK1 [700], TLR3 [1368], ANGPTL4 [1369], EPHB2 [1370], THBS1 [1371], ID3 [1372], NOX4 [1373], PCSK1 [1374], WNK4 [1375], VLDLR (very low density lipoprotein receptor) [1376], VDR (vitamin D receptor) [1377], GPC4 [1378], IFNE (interferon epsilon) [1379], ZFPM2 [1380], TLR2 [1381], IL6 [1382], SPINK1 [1383], GAS6 [1384], F11R [1385], SIGLEC15 [1386], ADAM12 [1387], MYOC (myocilin) [1388], USP2 [1389], SIK1 [1390], CCN2 [1391], IGFBP3 [1392], LYVE1 [1393], BGN (biglycan) [1394], EDA (ectodysplasin A) [1395], NTRK2 [1396], SLIT2 [1397], IRX3 [1191], NNMT (nicotinamide N-methyltransferase) [1398], CHI3L1 [1399], RUNX1 [1400], APLN (apelin) [1401], PGF (placental growth factor) [1402], HTR6 [742], NRG1 [1403], NPY4R [1404], CCDC80 [1405], KISS1 [1406], SLC6A15 [1407], ESYT3 [1408], SORBS1 [1409], SLC38A3 [1410], LXN (latexin) [1411] and TRH (thyrotropin releasing hormone) [1412] have been identified as a target for obesity. Studies have shown that VCAM1 [1413], STRA6 [1414], AQP8 [1415], FABP4 [1416], FLT1 [1417], BMP4 [633], PRL (prolactin) [1418], ADAMTS9 [1419], NDNF (neuron derived neurotrophic factor) [1420], DTX1 [1421], CD4 [1422], ADAMTS5 [1423], MMP8 [1424], ADRB2 [1425], PTPRD (protein tyrosine phosphatase receptor type D) [1426], INSIG1 [1427], LEPR (leptin receptor) [1428], IL15 [1429], APOE (apolipoprotein E) [1430], ACE (angiotensin I converting enzyme) [1431], LRP1B [1432], ALPP (alkaline phosphatase, placental) [1433], HKDC1 [1434], PNPLA3 [1435], FADS1 [1436], MMP3 [1437], FGFR2 [1438], IL33 [1439], FOXC2 [1440], FST (follistatin) [1441], HOXC8 [1442], BDNF (brain derived neurotrophic factor) [1443], NDRG2 [1444], ANGPTL4 [1445], TGFBI (transforming growth factor beta induced) [1446], VDR (vitamin D receptor) [1447], GPC4 [1448], CYP24A1 [1449], TLR2 [1450], IL6 [1451], KLF2 [1452], IGFBP3 [1453], SLIT2 [1454], APLN (apelin) [1455], NOTCH2 [1456], PGF (placental growth factor) [1457], NRG1 [1458], TLR1 [1459], CCDC80 [1460] and KISS1 [1461] play an important role in promoting the development of gestational diabetes mellitus. VCAM1 [1462], STRA6 [1013], WNT6 [1463], FABP4 [1464], SOX6 [1465], PLA2G7 [1466], MMP12 [1467], FAIM2 [770], SNCA (synuclein alpha) [1468], APOD (apolipoprotein D) [1469], LAG3 [1470], PREX1 [1471], FLT1 [1472], FABP5 [1473], TRPC3 [1474], THSD7A [1475], PRKCB (protein kinase C beta) [1476], PDE3B [1477], BMP4 [1478], PRL (prolactin) [1479], NEFL (neurofilament light chain) [1480], WNT5A [1481], TIMP3 [1306], CPE (carboxypeptidase E) [1482], ADAMTS9 [1483], NDNF (neuron derived neurotrophic factor) [1484], SNAP25 [1485], SCD (stearoyl-CoA desaturase) [1058], LDLR (low density lipoprotein receptor) [1486], CD4 [1487], TRPA1 [1488], RSPO3 [1489], PDE4B [1490], TH (tyrosine hydroxylase) [1491], CACNA1D [1492], MMP8 [1318], KDR (kinase insert domain receptor) [1493], ADRB2 [653], KL (klotho) [1494], PLAU (plasminogen activator, urokinase) [1321], CCND2 [1495], PTPRD (protein tyrosine phosphatase receptor type D) [1496], SIGLEC1 [1497], UCN2 [1324], CYP2J2 [1498], CYP1A1 [1499], NTN1 [1090], MSTN (myostatin) [1326], LEPR (leptin receptor) [1500], IL15 [1501], BMP2 [1502], APOE (apolipoprotein E) [1503], CUBN (cubilin) [1504], CASP1 [1505], MGLL (monoglyceride lipase) [1506], EFNB2 [1507], NID1 [1508], ABCG2 [1102], ACE (angiotensin I converting enzyme) [1509], STMN2 [1510], ICA1 [1511], TRPC6 [1512], GPBAR1 [1513], ELMO1 [1514], DOC2B [1515], ANK1 [1516], SORCS1 [1517], HKDC1 [1518], PNPLA3 [1519], FADS1 [1520], ACSL5 [1521], HLA-C [1522], S100A4 [1523], CORT (cortistatin) [1524], MMP3 [1525], GDNF (glial cell derived neurotrophic factor) [1526], CABLES1 [1354], IL33 [1527], NGF (nerve growth factor) [1528], FOXC2 [1529], COMP (cartilage oligomeric matrix protein) [1530], FST (follistatin) [1531], SORBS2 [1532], GATA6 [1533], PTGS2 [1141], DACT1 [1534], DGAT2 [1535], BDNF (brain derived neurotrophic factor) [1484], NDRG2 [1536], SGK1 [1537], TLR3 [1152], ANGPTL4 [1538], EPHB2 [1539], THBS1 [1540], MBP (myelin basic protein) [1541], NOX4 [1542], PI16 [1543], PCSK1 [888], TGFBI (transforming growth factor beta induced) [1544], IGF2BP1 [1545], WNK4 [1546], VLDLR (very low density lipoprotein receptor) [1547], VDR (vitamin D receptor) [1548], GPC4 [1274], PTPRN (protein tyrosine phosphatase receptor type N) [1549], EPHA4 [712], SOX9 [1550], MGP (matrix Gla protein) [1551], CYP24A1 [1552], TLR2 [1553], IL6 [1554], NPPB (natriuretic peptide B) [1555], SPINK1 [1556], GAS6 [1557], F11R [1558], FOXD1 [1559], ADAM12 [1560], KLF2 [1561], SIK1 [1562], FGF7 [1563], IGFBP3 [918], LYVE1 [1393], EDA (ectodysplasin A) [1395], SLIT2 [1564], IRX3 [1565], NNMT (nicotinamide N-methyltransferase) [1398], CHI3L1 [1566], RUNX1 [1567], APLN (apelin) [1568], COL4A3 [1569], NOTCH2 [1570], PDLIM5 [740], NRG1 [1571], DMRT2 [1572], NPY4R [1573], CD200 [1574], BST2 [1575], TFPI2 [1576], KISS1 [1406], MPP7 [1577], SORBS1 [1409], SLC38A3 [1578], CHN2 [1579] and TRH (thyrotropin releasing hormone) [1580] play an important regulatory role in the pathogenesis of diabetes mellitus. A previous bioinformatics study suggested that GRP (gastrin releasing peptide) [1581], AVPR1A [1582], ANO1 [1583], NTRK1 [1584], FGF10 [1585], MMP12 [1586], SNCA (synuclein alpha) [1587], CCR1 [1588], FLT1 [1589], FABP5 [1590], TRPC3 [1591], BMP4 [1592], PRL (prolactin) [1593], WNT5A [1594], TIMP3 [1595], SERPINB2 [1596], NLRP2 [1050], NPW (neuropeptide W) [1053], EREG (epiregulin) [1597], NDNF (neuron derived neurotrophic factor) [1598], SNAP25 [1056], SYT1 [1599], CD4 [1600], GPR183 [1601], TRPA1 [1602], PDE4B [1603], TH (tyrosine hydroxylase) [1604], MMP8 [1605], ADRB2 [1606], MGAT3 [1607], PLAU (plasminogen activator, urokinase) [1608], ASTN2 [1609], IL15 [1610], BMP2 [1611], APOE (apolipoprotein E) [1612], PDE9A [1613], MGLL (monoglyceride lipase) [1100], EFNB2 [1101], HMGCR (3-hydroxy-3-methylglutaryl-CoA reductase) [1614], ACE (angiotensin I converting enzyme) [1615], SYT9 [1616], TRPC6 [1617], XCR1 [1618], S100A4 [1619], MMP3 [1620], GDNF (glial cell derived neurotrophic factor) [1621], IL33 [1622], NGF (nerve growth factor) [1623], GDF6 [1133], COMP (cartilage oligomeric matrix protein) [1624], ACAN (aggrecan) [1625], PTGS2 [1626], GDF10 [1627], BDNF (brain derived neurotrophic factor) [1628], NDRG2 [1629], SGK1 [1630], TLR3 [1631], EPHB2 [1632], MBP (myelin basic protein) [1633], NOX4 [1619], SHANK2 [1634], PI16 [1635], DLX5 [1636], VDR (vitamin D receptor) [1637], ZFHX2 [1638], EPHA4 [1639], CYP24A1 [1640], ID2 [1641], TLR2 [1642], IL6 [1643], SPINK1 [1644], GAS6 [1645], KLF2 [1646], SIX1 [1647], CHL1 [1648], SLIT2 [1649], RUNX1 [1650], NOTCH2 [1651], PGF (placental growth factor) [1652], NRG1 [1653], NALCN (sodium leak channel, non-selective) [1654], LXN (latexin) [1655] and TRH (thyrotropin releasing hormone) [1656] could be used as diagnostic markers of pain. Therefore, studying the enriched genes involved in the regulation of endometriosis might be helpful to clarify the incidence or molecular pathogenic mechanisms of ovarian cancer, endometrial cancer, cervical cancer, hypertension, cardiovascular diseases, infertility, inflammation, polycystic ovarian syndrome, obesity, gestational diabetes mellitus, diabetes mellitus and pain.

Establishing PPI network and module analysis is friendly for researchers to investigate the underlying molecular mechanism of endometriosis for the reason that the DEGs would be grouped and ordered in the network judging by their interactions. PPI network and module analyses could help to find hub genes involved in the regulation of endometriosis. VCAM1 [110], ADRB2 [126], and DAPK1 [183] might serve as genetic markers of endometriosis. The expression of VCAM1 [190], ADRB2 [237], FOXQ1 [210], ACTBL2 [357], DAPK1 [379], ACTC1 [359], CST4 [229], NFIB (nuclear factor I B) [363], NFIX (nuclear factor I X) [1657] and ERG (ETS transcription factor ERG) [1658] were altered in the ovarian cancer. VCAM1 [478], FOXQ1 [492], DAPK1 [609] and ERG (ETS transcription factor ERG) [1659] promotes the development of cervical cancer. VCAM1 [615], ADRB2 [653] and DAPK1 [751] are associated to the risk of hypertension. VCAM1 [761], PRKCB (protein kinase C beta) [780], ADRB2 [808], PRKD1 [920], DAPK1 [947], ACTC1 [924] and NFIX (nuclear factor I X) [1660] are found to be associated with cardiovascular diseases. Studies show that VCAM1 [761], SNCA (synuclein alpha) [1026], ADRB2 [1073], FOXQ1 [1044], PRKD1 [1188], DAPK1 [947], ACTC1 [1192], CST1 [1030], NFIB (nuclear factor I B) [1198] and ERG (ETS transcription factor ERG) [1661] are involved in the process of inflammation. VCAM1 [1232] and ADRB2 [969] are an important regulator of the polycystic ovarian syndrome. VCAM1 [1287], PRKCB (protein kinase C beta) [1301] and ADRB2 [1319] might be regarded as a valuable biomarkers for diagnosis, treatment and prognosis of obesity. Altered expression of VCAM1 [1413] and ADRB2 [1425] are associated with gestational diabetes mellitus. Altered expression of VCAM1 [1462], SNCA (synuclein alpha) [1468], PRKCB (protein kinase C beta) [1476] and ADRB2 [653] are involved in the development and progression of diabetes mellitus. ADRB2 [969] has been known to be involved in cancer progression infertility. ADRB2 [1606] is molecular marker for pain. The expression levels of DAPK1 [183], CST1 [394] and NFIX (nuclear factor I X) [1662] have been proved to be altered in endometrial cancer patients. MDFI (MyoD family inhibitor), TNFRSF19 and FOXL1 served as novel biomarkers for endometriosis diagnosis and prognosis. This investigation identified the possible hub genes that were highly correlated with the PPI network to find the novel biomarkers associated in the pathogenesis of endometriosis.

In this investigations, the miRNA-hub gene regulatory network and TF-hub gene regulatory network of the hub genes in endometriosis were analyzed by using miRNet and NetworkAnalyst database. These analyses could help to find some miRNAs, TFs and hub genes involved in the regulation of endometriosis. CCND2 [246], VCAM1 [190], PDGFB (platelet derived growth factor subunit B) [221], PTCH1 [263], FOXQ1 [210], IGF2BP1 [324], ACTC1 [359], EPB41L3 [315], DAPK1 [379], hsa-mir-17-5p [1663], TCF3 [1664], RNF2 [1665], CLOCK (clock circadian regulator) [1666], SMARCA4 [1667] and TRIM28 [1668] provided a clear picture of the prognosis of patients with ovarian cancer. CCND2 [511], VCAM1 [478], PTCH1 [523], FOXQ1 [492], IGF2BP1 [568], EPB41L3 [560], DAPK1 [609], hsa-mir-17-5p [1669], TCF3 [1670] and TRIM28 [1671] as a biomarkers for cervical cancer. CCND2 [814], VCAM1 [615], PDGFB (platelet derived growth factor subunit B) [795], PRKCB (protein kinase C beta) [780], ACTC1 [924], DAPK1 [947], hsa-mir-17-5p [1672], hsa-mir-2110 [1673], TCF3 [1674] and SMARCA4 [1675] are associated with developing cardiovascular diseases. CCND2 [1083], VCAM1 [761], PDGFB (platelet derived growth factor subunit B) [1059], PTCH1 [1098], FOXQ1 [1044], IGF2BP1 [1162], ACTC1 [1192], DAPK1 [947], hsa-mir-2110 [1676], hsa-mir-10b-5p [1677], TCF3 [1678], NR1I2 [1679] and TRIM28 [1680] have been shown to be associated with inflammation. Studies have shown that CCND2 [1495], VCAM1 [1462], PTPRD (protein tyrosine phosphatase receptor type D) [1496], PRKCB (protein kinase C beta) [1476], IGF2BP1 [1545], hsa-mir-200a-3p [1681] and hsa-mir-10b-5p [1682] are an important biomarkers of diabetes mellitus. VCAM1 [110], PDGFB (platelet derived growth factor subunit B) [121], IGF2BP1 [162], DAPK1 [183] and hsa-mir-17-5p [1683] are a potential markers for the detection and prognosis of endometriosis. VCAM1 [1232], hsa-mir-17-5p [1684] and hsa-mir-2110 [1685] are a key initiator of polycystic ovarian syndrome. A previous study reported that VCAM1 [1287], PDGFB (platelet derived growth factor subunit B) [1311], PRKCB (protein kinase C beta) [1301], hsa-mir-17-5p [1686], hsa-mir-10b-5p [1687] and TRIM28 [1688] are altered expressed in obesity. VCAM1 [1413], PTPRD (protein tyrosine phosphatase receptor type D) [1426] and hsa-mir-17-5p [1689] are associated with prognosis in patients with gestational diabetes mellitus. IGF2BP1 [453], DAPK1 [183], PTPRD (protein tyrosine phosphatase receptor type D) [412], TCF3 [1690], SMARCA4 [1691] and TRIM28 [1692] biomarkers are vital for endometrial cancer. Research have shown that PTPRD (protein tyrosine phosphatase receptor type D) [656], PDGFB (platelet derived growth factor subunit B) [642], DAPK1 [751], hsa-mir-4432 [1693], SMARCA4 [1694] and TRIM28 [1695] participates in hypertension. IGF2BP1 [994], hsa-mir-17-5p [1696] and SMARCA4 [1697] plays an important role in the infertility. TRIM28 [1698] expression have been observed in pain. We identified ST8SIA4, MDFI (MyoD family inhibitor), KRT18, STX11, hsa-mir-3143, hsa-mir-6888-5p, hsa-mir-3122, hsa-mir-556-3p, hsa-mir-1229-5p, PHC1, HOXC9, PRDM14 and HTT (huntingtin) might serve as novel biomarkers for endometriosis. We suggest that exercise can regulate the expression of these miRNAs, TFs and hub genes, thereby inhibiting the occurrence and development of endometriosis.

## Conclusions

The current investigation identified biomarkers and pathways which might be involved in endometriosis progression through the integrated analysis of NGS dataset. These results might contribute to a better understanding of the molecular mechanisms which underlie endometriosis and provide a series of potential biomarkers. However, further experiments are required to verify the findings of the current investigations. Therefore, further experiments with additional patient cohorts are also required to confirm the results of this investigations. In vivo and in vitro investigation of gene and pathway interaction is essential to delineate the specific roles of the identified biomarkers, which might help to confirm biomarker functions and reveal the molecular mechanisms underlying endometriosis.

## Acknowledgement

I thanks very much to Peixin Jiang, Baylor College of Medicine, Houston, TX, USA,the author who deposited their NGS dataset GSE243039, into the public GEO database.

## Ethical approval

This article does not contain any studies with human participants or animals performed by any of the authors.

## Informed consent

No informed consent because this study does not contain human or animals participants.

## Availability of data and materials

The datasets supporting the conclusions of this article are available in the GEO (Gene Expression Omnibus) (https://www.ncbi.nlm.nih.gov/geo/) repository.

[(GSE243039) https://www.ncbi.nlm.nih.gov/geo/query/acc.cgi?acc=GSE243039]

## Consent for publication

Not applicable.

## Author Contributions

1. B. V. - Writing original draft, and review and editing
2. C. V. - Software and investigation

